# Mapping the biosynthetic pathway of a hybrid polyketide-nonribosomal peptide in a metazoan

**DOI:** 10.1101/2021.05.28.446193

**Authors:** Likui Feng, Matthew T. Gordon, Ying Liu, Kari B. Basso, Rebecca A. Butcher

## Abstract

Hybrid polyketide synthase (PKS) and nonribosomal peptide synthetase (NRPS) systems typically use complex protein-protein interactions to facilitate direct transfer of intermediates between megasynthases. In the nematode *Caenorhabditis elegans*, PKS-1 and NRPS-1 produce the nemamides, the only known hybrid polyketide-nonribosomal peptides in animals, through a poorly understood mechanism. Here, we use genome editing and mass spectrometry to map the roles of individual PKS-1 and NRPS-1 enzymatic domains in nemamide biosynthesis. Furthermore, we show that nemamide biosynthesis requires at least five additional stand-alone enzymes that are encoded by genes distributed across the worm genome. We identify the roles of these enzymes in the biosynthetic pathway and discover a novel mechanism of trafficking intermediates between a PKS and an NRPS. Specifically, we show that the enzyme PKAL-1 activates an advanced polyketide intermediate as an adenylate and directly loads it onto a carrier protein in NRPS-1. This trafficking provides a means by which a PKS-NRPS system can expand its biosynthetic potential and is likely important for the regulation of nemamide biosynthesis.

## Introduction

In the past 15 years, it has become clear that animal genomes encode biosynthetic pathways for many microbial-like secondary metabolites.^1–11^ Although in some cases these pathways were acquired from microorganisms through horizontal gene transfer, in the majority of cases these pathways are thought to have evolved independently in animals.^1,3,7,9–11^ Investigating the rich biochemistry of animals thus has the potential to reveal many new chemical insights and biosynthetic strategies. Furthermore, these studies are poised to reveal how the animal biosynthetic machinery is integrated with the higher order complexity found in animals, including multiple organelles and tissues, integrated signaling pathways, and complex life history traits.

Nematodes, in particular, have been shown to have a rich biosynthetic repertoire.^12^ Infact, many nematode genomes encode multi-module Type I PKSs and NRPSs for assembly-linetype biosynthesis of polyketides and nonribosomal peptides.^3–5,8^ Using the megasynthetases PKS-1 and NRPS-1, the model nematode *C. elegans* has been shown to produce a remarkable class of hybrid polyketide-nonribosomal peptides known as the nemamides in two essential neurons, the canal-associated neurons (CANs) (**Fig. 1a**).^8^ The nemamides promote survival of the worm during starvation in a manner that is independent of the transcription factor DAF-16/FOXO.^8,13^ Since PKS-1 and NRPS-1 homologs are found in most nematode species, including parasitic ones, it is likely that other nematode species produce nemamide-like molecules and that these natural products play a conserved role in nematode biology.^8^ However, the biosynthesis of the nemamides, as well as how it is regulated, is poorly understood.

**Figure 1.**
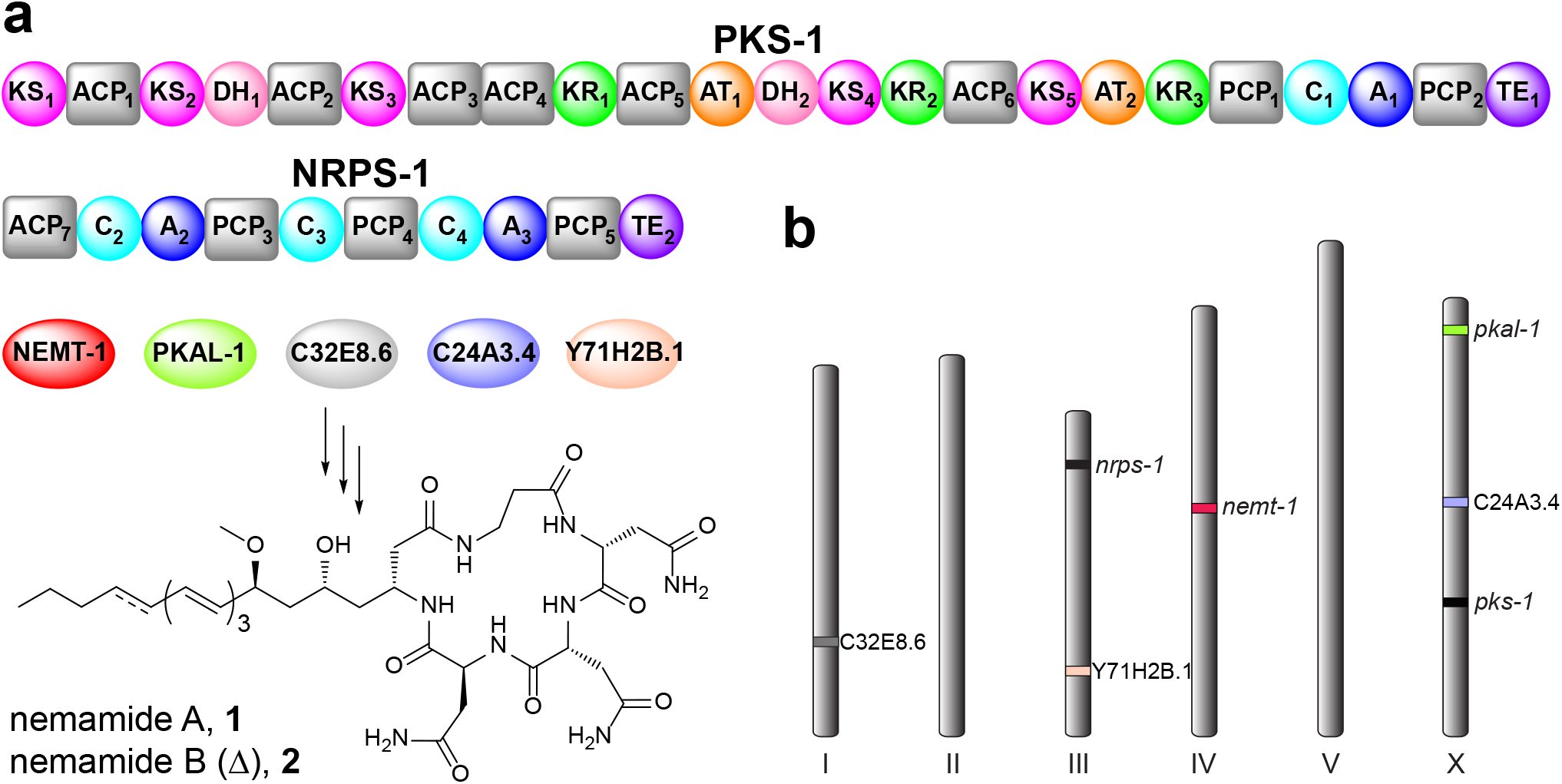
Enzymes required for nemamide biosynthesis. (**a**) The domain organization of PKS-1 and NRPS-1 is shown, along with five additional free-standing enzymes (NEMT-1, PKAL-1, C32E8.6, C24A3.4, and Y71H2B.1) that were demonstrated in this study to be required for nemamide biosynthesis. To facilitate annotation of the worm mutant strains generated in this study, domains have been numbered according to the order of their appearance in PKS-1 and NRPS-1. The ACP_7_ domain was identified and its functional role was confirmed in this study. (**b**) The approximate chromosomal location of *pks*-*1*, *nrps*-*1* and the five additional genes demonstrated to be required for nemamide biosynthesis in this study.

In Type I PKSs and NRPSs, each module is responsible for incorporating a different building block, such as malonyl- or methylmalonyl-CoA in the case of PKSs and L-, D-, or nonproteinogenic amino acids in the case of NRPSs.^14,15^ Acyltransferase (AT) domains are responsible for loading building blocks onto acyl carrier proteins (ACPs) in PKSs, while adenylation (A) domains are responsible for loading building blocks onto peptidyl carrier proteins (PCPs) in NRPSs. The building blocks are then linked together by ketosynthase (KS) domains in PKSs and condensation (C) domains in NRPSs. Additional domains can be present in the modules, such as ketoreductase (KR) and dehydratase (DH) domains in PKSs that control the oxidation state of the β-carbon, as well as methyltransferase and aminotransferase domains in PKSs and NRPSs that can further modify the natural product. Once synthesis of the natural product on a PKS or NRPS is complete, it is typically cleaved from the synthetase by a thioesterase (TE) domain. Further complexity can be introduced through hybrid systems that combine PKS and NRPS modules. These two types of modules are sometimes encoded in the same protein, but they are more often encoded in separate proteins. In this latter case, protein protein interactions facilitate the direct passage of biosynthetic intermediates between the two proteins.^16^

Domain analysis of PKS-1 and NRPS-1 provides a number of tantalizing clues indicating that the biosynthesis of the nemamides deviates from a canonical pathway in several respects.^8^ Both PKS-1 and NRPS-1 contain stretches of protein sequence with no homology to known enzymatic domains, and the enzymatic domains that can be identified are out of order and/or have diverged significantly from those found in other systems.^8^ For example, the substrate preferences of the A domains cannot be predicted based on the presence of key residues present in their active sites.^17^ Furthermore, there are no obvious domains that would enable certain structural features present in the nemamides, including the methoxy and amino groups, to be incorporated. Given that biosynthetic genes are not clustered in animals as they are in microorganisms, the identification of these missing domains in the worm represents a challenge. Additionally, while there is a TE domain at the C-terminus of NRPS-1, which presumably cleaves the natural product from the synthetase, there is also, strangely, a TE domain at the C-terminus of PKS-1 of unknown function.

Here, we map the biosynthesis of a complex metabolite in an animal system through genetic manipulation of the biosynthetic genes *in vivo* followed by comparative metabolomics. We show that the biosynthesis of the nemamides requires at least seven genes distributed across the genome that are united by their common expression in the CANs (**Fig. 1a,b**). Furthermore, we uncover the biosynthetic roles of these genes and show that two of them encode a methyltransferase (NEMT-1) and polyketide-ACP ligase (PKAL-1) that act *in trans* and are required for the trafficking of intermediates between PKS-1 and NRPS-1. PKAL-1 represents a unique enzyme in that it loads a complex polyketide intermediate onto an NRPS for further chemical elaboration.

## Results

### NRPS-1 is responsible for incorporating all of the amino acid components of the nemamides

Assuming a linear, assembly-line mechanism in nemamide biosynthesis, we predicted that the β-Ala moiety in the nemamides is installed by the C-terminal NRPS module of PKS-1 and that the two D-Asn and final L-Asn moieties are installed by the NRPS modules of NRPS-1. However, the lack of sequence homology of the A domains in PKS-1 and NRPS-1 to bacterial and fungal A domains precluded predictions of the amino acid substrate specificities of these domains that might have supported our model.^8,17^ Furthermore, our inability to express in *E. coli* any of the PKS-1 or NRPS-1 A domains (either as excised domains or as part of multi-domain constructs) prevented us from analyzing the substrate preferences of the A domains *in vitro*. Therefore, we decided to inactivate specific domains in PKS-1 and NRPS-1 in the worm in order to map their biosynthetic role. First, we used CRISPR-Cas9 to inactivate in the worm the TE domain in NRPS-1 by replacing the catalytic serine with an alanine to generate strain *nrps*-*1[TE2_S2803A]* (**Supplementary Fig. 1**). This worm strain does not make the nemamides, consistent with the role of the NRPS-1 TE domain in cleaving nemamide from the synthetase through the formation of a macrolactam (**Supplementary Fig. 2**).

Unexpectedly, the *nrps*-*1[TE2_S2803A]* strain accumulates a number of intermediates in the biosynthesis of the nemamides, one with no amino acids incorporated (**3**), one with β-Ala incorporated (**4**), one with βAla-D-Asn incorporated (**5**), and one with β-Ala-D-Asn-D-Asn incorporated (**6**) (**Fig. 2a,b; Supplementary Fig. 3**). Given that the nemamides are produced in very low amounts in *C. elegans*, and given that the biosynthetic intermediates are produced at even lower amounts, the intermediates had to be partially purified to enable identification. 2-3L of worms grown in high-density axenic culture enabled the production of 3-5g of worms, which were used to generate extracts that were that were then purified through two chromatographic steps. Intermediates were followed based on the characteristic UV spectrum of the triene or tetraene moiety that is present in them. Fortunately, this UV signature appears to be quite unique to nemamide and nemamide intermediates in *C. elegans* extracts. We verified the accumulation of intermediates **3**, **4**, **5**, and **6** in the NRPS-1 TE domain mutant strain using high resolution LCMS/MS (**Supplementary Fig. 4-7**). The amounts of the intermediates detected in this mutant were less than 10% of the mean amount of nemamide A in wild-type worms (**Supplementary Fig. 3**). The fact that the NRPS-1 TE domain mutant strain does not accumulate the linear form of nemamide A with the last L-Asn incorporated, suggests that this intermediate likely remains covalently attached to NRPS-1 terminal PCP (PCP_5_) in the mutant.

**Figure 2.**
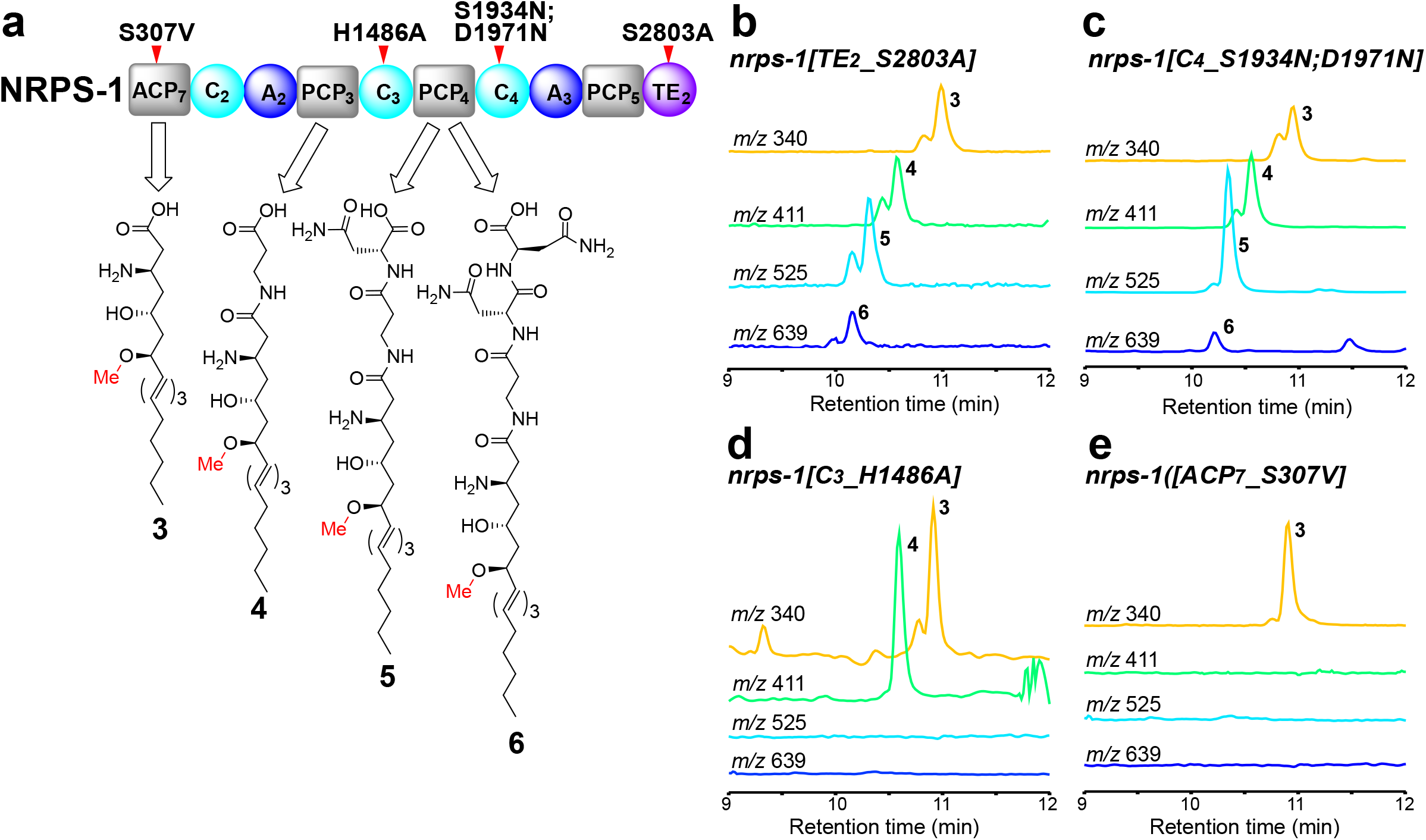
Analysis of biosynthetic intermediates in mutant strains in which specific NRPS-1 domains have been inactivated. (**a**) The structures of the intermediates that were identified in the NRPS-1 mutant strains and the proposed carrier proteins that carry them as the corresponding thioesters. The domains that were mutated are indicated with a red tick mark. The chemical structures of the intermediates were confirmed through high resolution LC-MS/MS (**Supplementary Fig. 4-7**). (**b**-**e**) Representative extracted ion chromatograms of intermediates **3** ([M+H]^+^ *m/z* 340), **4** ([M+H]^+^ *m/z* 411), **5** ([M+H]^+^ *m/z* 525), and **6** ([M+H]^+^ *m/z* 639) in the NRPS-1 TE2 domain mutant, *nrps*-*1(reb12[TE2_S2803A])* (**b**), the NRPS-1 C4 domain mutant, *nrps*-*1(gk186409[C4_S1934N];gk186410[C4_D1971N])* (**c**), the NRPS-1 C3 domain mutant, *nrps*-*1(reb10[C3_H1486A])* (**d**), and the NRPS-1 ACP_7_ domain mutant, *nrps*-*1(reb8[ACP_7__S307V])* (**e**). Note that for the *nrps*-*1_C4* domain mutant contains a mutation in an active site Ser, as well as an additional mutation (D1971N) that is not expected to affect the domain’s function.

We hypothesized that we might be able to map the biosynthesis of the nemamides by NRPS-1 by targeting individual domains in NRPS-1 and analyzing the intermediates that accumulate. We began with domains acting at the end of the nemamide biosynthetic pathway and worked our way backward. The *nrps*-*1[C4_S1934N;D1971N]* strain, in which an active site Ser of the last NRPS-1 C domain was mutated, does not make the nemamides and accumulates intermediates **3**, **4**, **5**, and **6** (**Fig. 2a,c**; **Supplementary Fig. 1-3**). This result suggests that the last C domain in NRPS-1 incorporates the L-Asn into the growing chain before cyclization by the NRPS-1 TE domain to form the nemamides. The *nrps*-*1[C3_H1486A]* strain, in which the catalytic histidine of the second to last C domain in NRPS-1 was mutated, does not make the nemamides and accumulates intermediates **3** and **4** (**Fig. 2a,d**; **Supplementary Fig. 1-3**). This result suggests that this C domain incorporates the two D-Asn moieties into the nemamides. Certain C domains catalyze both epimerization and condensation of the amino acids that they incorporate, but none of the C domains in NRPS-1 have the characteristic features of these domains (**Supplementary Fig. 8**).^18^ Thus, it is unclear whether incorporation of the D-Asn moieties into the nemamides involves loading of D-Asn by an A domain or loading of L-Asn, followed by epimerization to D-Asn by an unidentified epimerase.

The N-terminus of NRPS-1 contains a stretch of unannotated sequence of 400 amino acids that has no obvious homology to PKS or NRPS domains, based on anti-SMASH or BLAST analysis.^19^ Using the modeling program SWISS-model, however, we were able to identify an ACP domain between residues 259-361 of this sequence (annotated as ACP_7_).^20^ The *nrps*-*1[ACP_7__S307V]* strain, in which the predicted site of phosphopantetheinylation in ACP_7_ is mutated, does not make the nemamides, demonstrating that this ACP domain is functionally relevant (**Supplementary Fig. 1-2**). Surprisingly, this strain specifically accumulates intermediate **3**, but not **4**, revealing that the β-alanine residue in the nemamides is not incorporated by the C-terminal NRPS module of PKS-1, as we had originally proposed, but by the N-terminal NRPS module of NRPS-1 (**Fig. 2a,e**; **Supplementary Fig. 3**). Since disruption of the NRPS-1 ACP_7_, C3, C4, and TE2 domains leads to the accumulation of intermediate **3**, this intermediate is most likely generated by PKS-1 and transferred onto NRPS-1 ACP_7_ for further elongation. Furthermore, because intermediate **3** contains the amino and methoxy groups, incorporation of these groups must precede the biosynthetic steps carried out by NRPS-1.

### Role of the C-terminal NRPS module and TE domain in PKS-1

Given that the β-alanine moiety is incorporated by NRPS-1, it was unclear whether the C-terminal NRPS module in PKS-1 would be required for nemamide biosynthesis. The domain organization of this NRPS module in PKS-1, as well as the sequence of the A domain in this module, are highly conserved across nematode evolution, suggesting that the NRPS module of PKS-1 does play an important role in the biosynthesis (**Supplementary Table 1**).^8^ Three mutant worm strains containing mutations in either the A1, C1, or PCP2 domains of the NRPS module in PKS-1 were generated (Fig. 3; Supplementary Fig. 1). While the C1 domain mutant could still produce some nemamides (less than 40%), the A1 domain mutant made only very minor amounts of nemamides and the PCP1 domain mutant did not make any at all (**Fig. 3**; **Supplementary Fig. 9**). Thus, although the C-terminal NRPS module of PKS-1 is not involved in incorporating β-alanine, it is required for nemamide biosynthesis. None of these strains accumulated any intermediates with UV signatures characteristic of trienes or tetraenes, suggesting that if any such intermediates do accumulate, they remain covalently linked to the synthetase.

**Figure 3.**
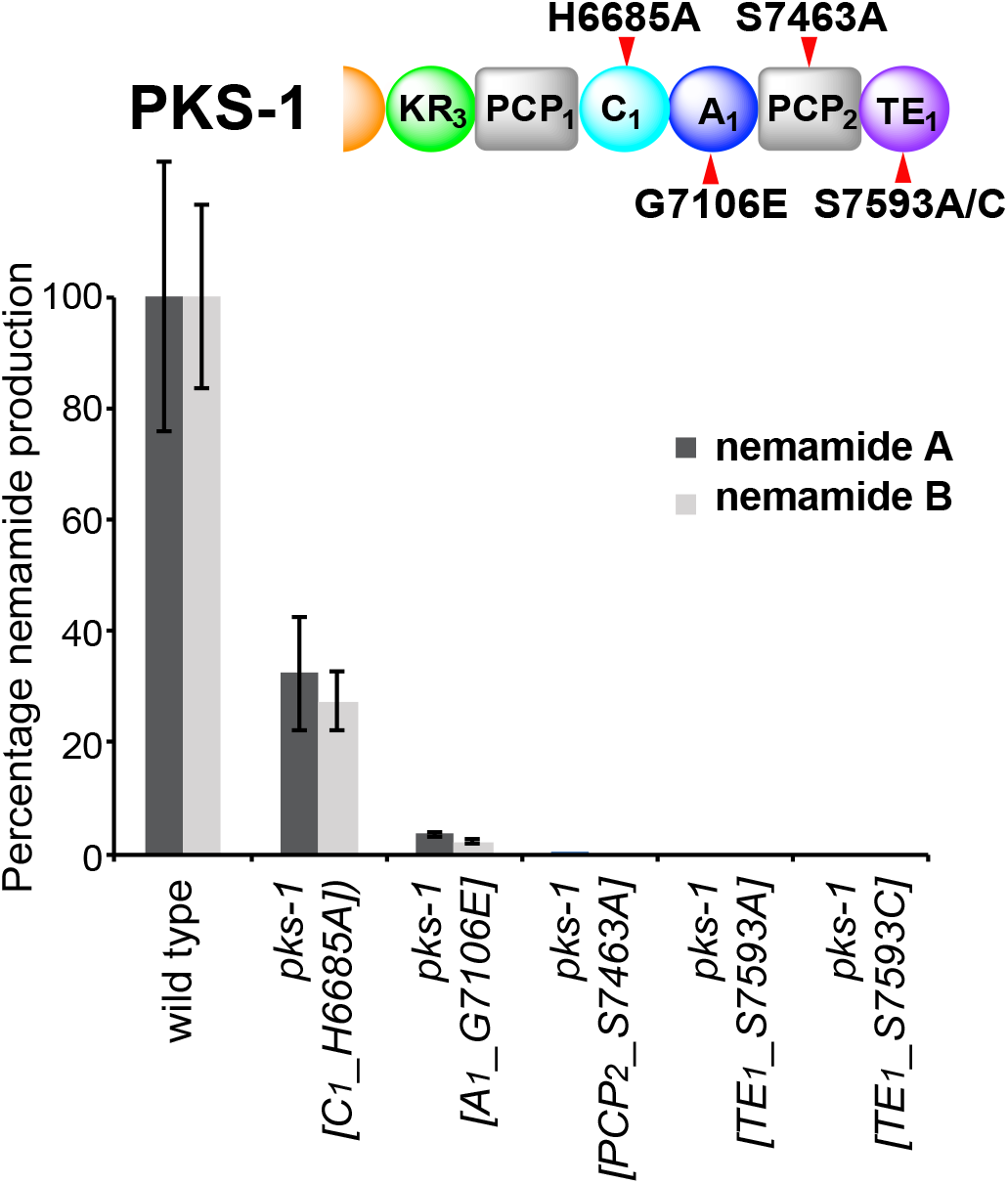
Analysis of nemamide production in mutant strains in which different domains in the C-terminal NRPS module of PKS-1 have been inactivated. The C-terminal NRPS module was inactivated by mutating essential residues in each domain, which is predicted to be active. In the C1 domain, the catalytic residue His 6685 was mutated to Ala. In the A1 domain, Gly 7106, which is essential for cofactor binding, was mutated to Glu. In the PCP2 domain, the site of phosphopantheinylation, Ser 7463, was mutated to Ala. In the TE1 domain, Ser 7593, which is part of the catalytic triad, was mutated to either Ala or Cys. The domains that were mutated are indicated with a red tick mark. The amount of nemamide A or B in each strain relative to the amount of nemamide A in wild-type worms was based on the UV absorbance of the compounds and was normalized by the total mass of dried worms used for extraction. Data represent the mean ± standard deviation of at least two to three independent experiments.

The NRPS module in PKS-1 is followed by a C-terminal TE domain of unknown function. Comparison of this TE domain to bacterial TEI domains, which are involved in the cleaving of final products from the synthetase, and bacterial TEII domains, which facilitate synthesis by cleaving misloaded intermediates from carrier proteins, suggests that the PKS-1 TE domain is most similar to bacterial TEI domains found in hybrid PKS-NRPS systems.^8,21,22^ We generated two mutant worm strains, *pks*-*1[TE1_S7593A]* and *pks*-*1[TE1_S7593C]*, in which the catalytic serine residue of the PKS-1 TE domain was mutated to either an alanine or a cysteine, and showed that these strains do not make any nemamides or accumulate any biosynthetic intermediates (**Fig. 3**; **Supplementary Fig. 1**; **Supplementary Fig. 9**). This result suggests that the PKS-1 TE domain is not functioning in an editing fashion as a TEII domain, since such an editing enzyme would typically not be essential for biosynthesis, but only facilitate it.^22^ Instead, this result suggests that the PKS-1 TE domain is playing an essential role, most likely in the trafficking of intermediates between PKS-1 and NRPS-1.

### Identification of *trans*-acting enzymes involved in nemamide biosynthesis

We hypothesized that certain steps in nemamide biosynthesis would require additional enzymes beyond PKS-1 and NRPS-1 and that these enzymes would likely also be expressed in the CANs. Single-cell gene expression profiling has been performed in multiple cell types in *C. elegans*, including the CANs.^23,24^ 38 genes, including *pks*-*1* and *nrps*-*1*, show selective expression in the CANs that is at least 5-fold higher than the next tissue in which they are expressed.^23^ Based on their homology and predicted enzymatic activities, we screened a number of these genes by analyzing nemamide production in loss-of-function mutant worm strains. These data show that at least five additional genes are required for nemamide biosynthesis, F49C12.10, T20F7.7, C32E8.6, C24A3.4, and Y71H2B.1 (**Fig. 4a**). F49C12.10, which we named NEMT-1 (<u>NE</u>mamide *O*-<u>M</u>ethyl<u>T</u>ransferase-1), is predicted to be an *O*-methyltransferase since its closest non-nematode homolog is phthiotriol methyltransferase (with 31% identity), which methylates a precursor to cell surface-associated apolar lipids in mycobacteria.^25,26^ T20F7.7 and C32E8.6 are homologous to acyl-CoA synthetases, but failure of the T20F7.7 and C32E8.6 genes to inter-rescue suggests that they play distinct roles in nemamide biosynthesis (**Supplementary Fig. 10**). T20F7.7 was previously named ACS-9 due to its homology, but for reasons described below, we have renamed it PKAL-1 (<u>P</u>oly<u>K</u>etide-<u>A</u>CP <u>L</u>igase-1). Lastly, C24A3.4 is annotated as a methylacyl-CoA racemase or CoA transferase, and Y71H2B.1 is annotated as a fatty-acyl CoA binding protein. For all five genes, nemamide production could be rescued in loss-of-function strains by complementing the genes by expressing them under the control of their own promoter (**Supplementary Fig. 11**). By generating translational GFP reporter strains, we were able to verify that all of the additional genes required for nemamide biosynthesis are primarily expressed in the CANs, although C24A3.4 is also expressed in the intestine (**Supplementary Fig. 12**). Interestingly, unlike natural product biosynthetic genes in bacteria and fungi, which are often clustered, the nemamide biosynthetic genes are scattered across the *C. elegans* genome with C32E8.6 on chromosome I, *nrps*-*1* and Y71H2B.1 on different arms of chromosome III, *nemt*-*1* on chromosome IV, and *pks*-*1*, *pkal*-*1*, and C24A3.4 spread across the X chromosome (**Fig. 1b**).

**Figure 4.**
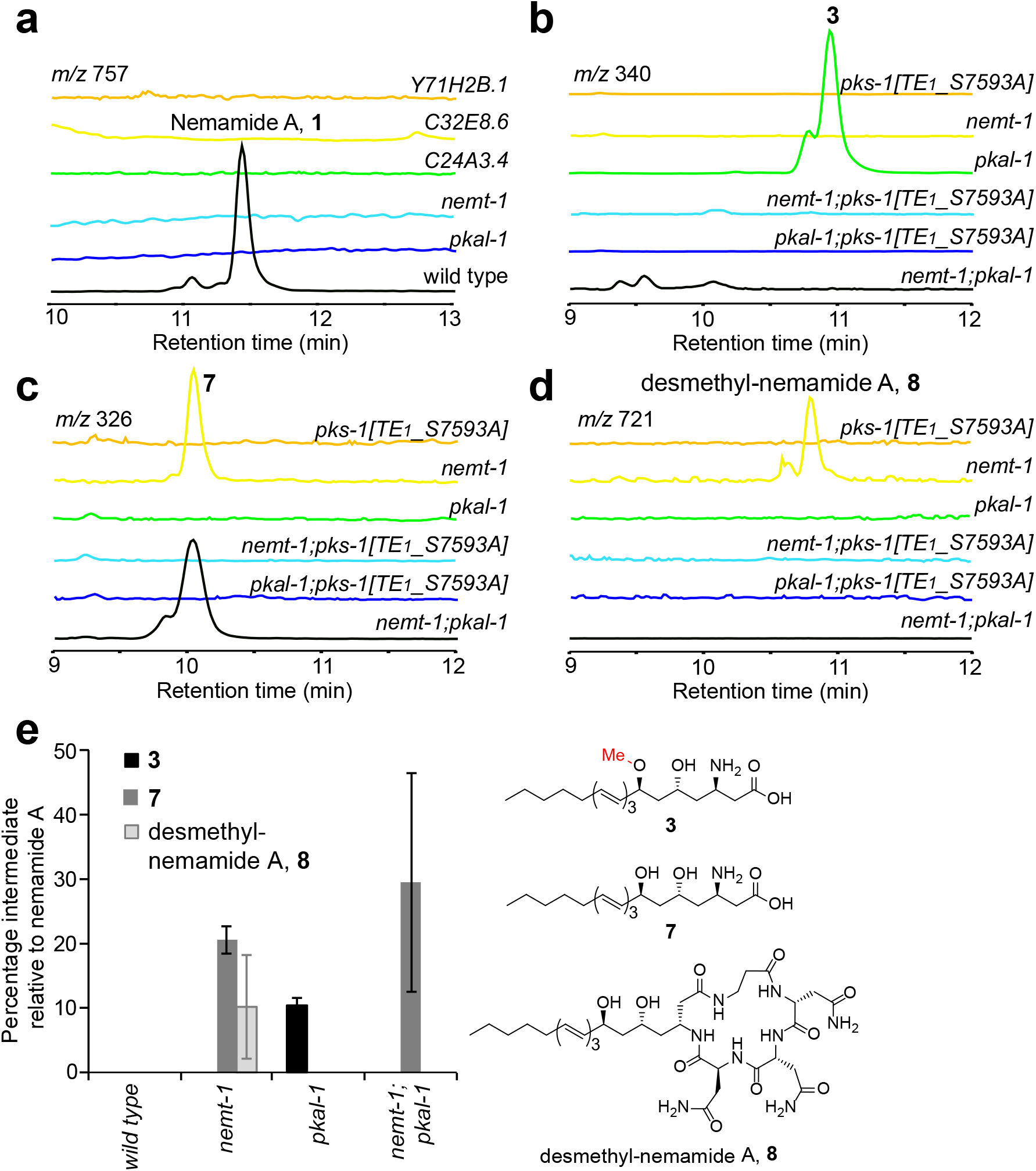
Analysis of nemamides and biosynthetic intermediates in mutant strains in which *trans*-acting enzymes have been inactivated. (**a-d**) Representative extracted ion chromatograms of nemamide A, **1**, (**a**) and intermediate **3** (**b**), intermediate **7** (**c**), and desmethyl-nemamide **8** (**d**) in the *pks*-*1(reb11[TE1_S7593A])*, *pkal*-*1(reb28)*, *nemt*-*1(reb15)*, and double mutants. (**e**) The amounts of intermediates **3**, **7**, and **8** in mutant strains relative to the amount of nemamide A in wild-type worms was based on the UV absorbance of the compounds and was normalized by the total mass of dried worms used for extraction. Data in (**e**) represent the mean ± standard deviation of three independent experiments.

### NEMT-1 and PKAL-1 traffic intermediates between PKS-1 and NRPS-1

To investigate the role of the *trans*-acting enzymes in nemamide biosynthesis, we characterized the intermediates that accumulate in the corresponding mutant worm strains. The *pkal*-*1* mutant strain accumulates **3**, which is the same intermediate that accumulated in the strain with the ACP_7_ domain of NRPS-1 mutated (**Fig. 4b,e**). On the other hand, the *pkal*-*1*; *pks*-*1[TE1_S7593A]* double mutant strain does not accumulate **3**, indicating that PKAL-1 functions downstream of the PKS-1 TE1 domain (**Fig. 4b,e**). We hypothesized that PKAL-1 might be involved in loading **3** onto the ACP_7_ domain of NRPS-1 for further extension.

The *nemt*-*1* mutant strain accumulates intermediate **7**, which is similar to **3**, but lacks the methoxy group (**Fig. 4c,e**; **Supplementary Fig. 13**). This result indicates that NEMT-1 is the *O*-methyltransferase that installs the methoxy group that is present in the nemamides. The *nemt*-*1* mutant strain also produces **8**, which is similar to nemamide A, but lacks the methoxy group (**Fig. 4d,e; Supplementary Fig. 14**). The fact that the *nemt*-*1* mutant can make desmethyl-nemamide **8**, suggests that PKAL-1 can load **7** onto the ACP_7_ domain of NRPS-1, even though it is missing the methoxy group. However, because the *nemt*-*1* mutant accumulates intermediate **7**, PKAL-1 and/or NRPS-1 likely prefers methylated substrates over unmethylated ones (**Fig. 4e**). Unlike the *nemt*-*1* single mutant, the *nemt*-*1*; *pks*-*1_TE1* double mutant does not accumulate intermediate **7** or desmethyl-nemamide **8**, suggesting a possible model in which NEMT-1 functions downstream of the PKS-1 TE1 domain. Thus, our genetic data suggest that NEMT-1 methylates intermediate **7** to form **3**, and then PKAL-1 is involved in loading **3** onto the ACP_7_ domain of NRPS-1 for further extension.

No biosynthetic intermediates were found in Y71H2B.1, C24A3.4, or C32E8.6 loss-of-function mutant worm strains. Given that intermediates also do not accumulate in *pks*-*1* mutant worm strains, these three genes may function in earlier stages of nemamide biosynthesis, potentially either in initiation or in other steps involving PKS-1.

### Biochemical activity of PKAL-1

Based on its sequence homology, we hypothesized that PKAL-1 might function similar to a fatty acyl-CoA ligase (FACL). That is, PKAL-1 might activate 3 as the adenylate and then react that intermediate with CoA to form a CoA-thioester that is subsequently loaded onto the ACP_7_ domain of NRPS-1. To biochemically characterize the role of PKAL-1 in trafficking between PKS-1 and NRPS-1, we cloned the enzyme from a cDNA library, expressed it in *E. coli*, and purified it for biochemical characterization. We then incubated PKAL-1 with fatty acids of various lengths, ATP, and CoA. PKAL-1 can activate a variety of medium and long-chain fatty acids as the corresponding fatty acyl-AMP but cannot further convert them to the corresponding fatty acyl-CoA (**Fig. 5a, Supplementary Fig. 15**). A negative control, PKAL-1(K488A), in which a lysine predicted to be important for catalysis has been mutated,^14^ has no activity towards fatty acids. These results suggest that PKAL-1 is analogous to a fatty acyl-AMP ligase (FAAL) rather than an FACL. FAAL enzymes activate fatty acids as the corresponding fatty acyl-AMP but then transfer the fatty acyl group to the phosphopantetheinyl arm of a carrier protein instead of CoA.^27–29^ FAAL enzymes typically have an insertion motif that prevents the movement between the larger N-terminal domain and the smaller C-terminal domain that occurs between the adenylation and CoA-ligase reactions.^27,28^ Certain FAAL enzymes lack the insertion motif, but instead have additional interactions between the N-terminal and C-terminal domains that are thought to prevent the CoA-ligase reaction from occurring.^30^ Although sequence alignment of PKAL-1 demonstrates that it is missing the insertion motif (**Supplementary Fig. 16**), structural modeling of PKAL-1 indicates that it might lack CoA ligase activity due to the absence of an effective binding site for CoA (**Supplementary Fig. 17**).

**Figure 5.**
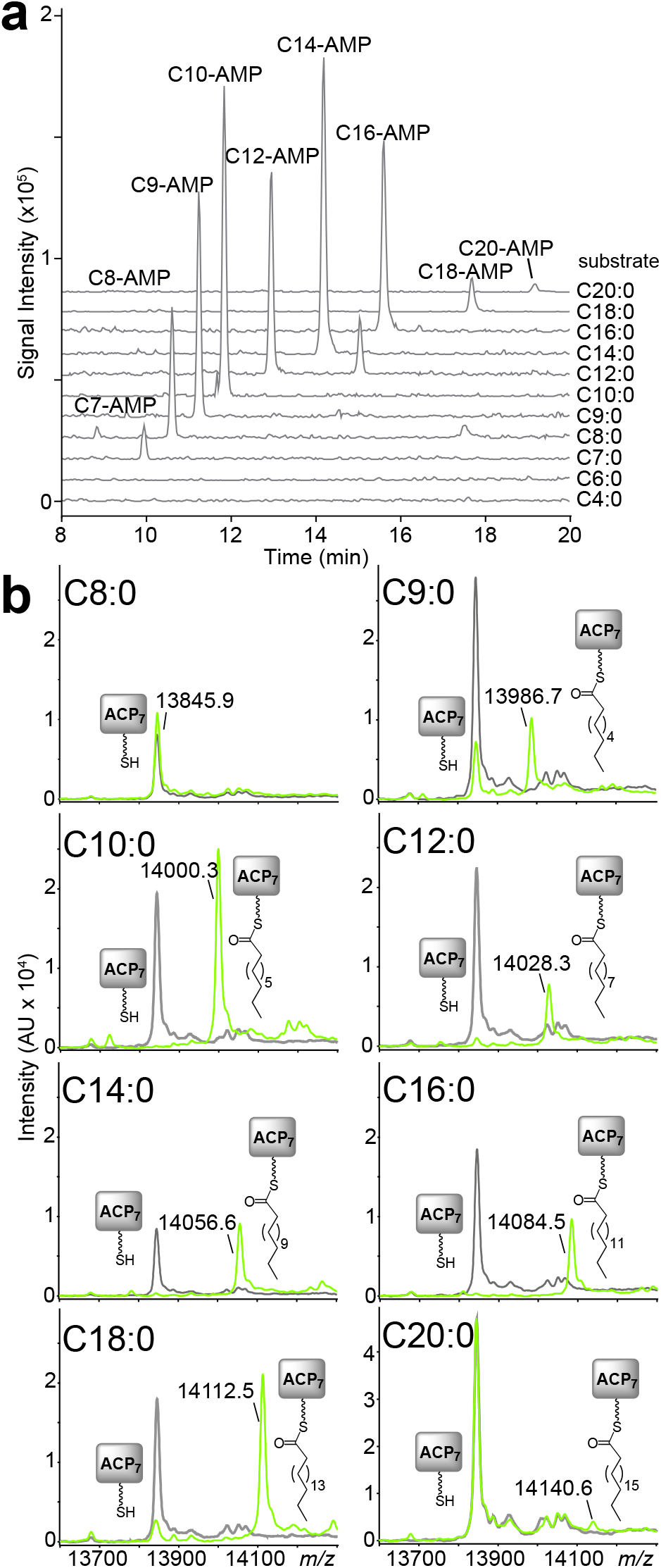
*In vitro* activity of PKAL-1 against fatty acid substrates. (**a**) Activity of PKAL-1 towards fatty acids of various length in the presence of ATP and CoA. The identities of the peaks were verified based on their *m/z* in both positive and negative modes, as well as their UV spectrum. The catalytic mutant PKAL-1(K488A) did not show any product formation. The inability of PKAL-1 to form CoA-thioesters was verified with synthetic standards (**Supplementary Fig. 15**). (**b**) MALDI-TOF analysis of the holo-ACP_7_ domain from NRPS-1 incubated with PKAL-1, fatty acids of various lengths, and ATP.

FAAL enzymes play an important role in the biosynthesis of lipid-modified polyketides and nonribosomal peptides, such as mycolic acids that are essential for mycobacterial growth, isonitrile lipopeptides that promote mycobacterial virulence, and diverse lipopeptides that are widespread in cyanobacteria.^27–32^ In order to determine whether PKAL-1 can load fatty acids that are similar in length to intermediate **3** onto the ACP_7_ domain of NRPS-1, the excised ACP_7_ domain was first co-expressed in *E. coli* with the promiscuous phosphopantetheinyl transferase Sfp,^33^ enabling an efficient purification of the holo-ACP_7_ domain. This holo-ACP_7_ domain was incubated with PKAL-1, ATP and fatty acids of various lengths, and the products were analyzed by MALDI-TOF. PKAL-1 could efficiently load fatty acids that were 9 to 18 carbons in length and could load a minor amount of C20 fatty acids. (**Fig. 5b**). This result is consistent with a role for PKAL-1 in activating **3**, which is 18 carbons in length. PKAL-1 was unable to load fatty acids onto a different ACP, ACP_1_ from PKS-1 (**Supplementary Fig. 18**). Thus, PKAL-1 must engage in specific interactions with the ACP_7_ domain of NRPS-1 in order to load fatty acids onto it. While FAAL enzymes have been shown to activate fatty acids to initiate polyketide and nonribosomal peptide biosynthesis,^27,28,32,34,35^ PKAL-1 is unusual in that it activates an advanced polyketide intermediate for loading onto an NRPS for further elaboration.

## Discussion

Using comparative metabolomics, we have mapped the biosynthetic pathway to the nemamides, a remarkable family of hybrid polyketide-nonribosomal peptides biosynthesized in the CANs of *C. elegans*. Furthermore, we have determined the biosynthetic roles of five additional enzymes from *C. elegans* that function *in trans* in nemamide biosynthesis. Although the different genes that are required for nemamide biosynthesis are encoded in disparate locations across the worm genome, these genes share a common feature that they are expressed in the CANs. Our work has shown that at least 7 of the 38 genes that have enriched expression in the CANs^23^ are involved in nemamide biosynthesis, suggesting that nemamide biosynthesis may be a primary function of these enigmatic neurons.

Our data show that all of the amino acids in the nemamides, including the β-alanine, are incorporated by NRPS-1 and that the C-terminal NRPS module of PKS-1 has other functions in nemamide biosynthesis (**Fig. 6**). The A1 and PCP2 domains of PKS-1 are required for nemamide production, so the NRPS module of PKS-1 does play a role in nemamide biosynthesis. Furthermore, both the domain organization of this NRPS module in PKS-1 and the sequence of the A domain have been conserved across nematode species.^8^ As we have not be able to identify an aminotransferase for incorporation of the amino group into the nemamides, this module may be involved in this process. A possible mechanism for the incorporation of an amino group into the nemamides is suggested by the biosynthesis of the β-amino fatty acid starter unit in the macrolactam family of antibiotics that includes BE-14106 and ML-449.^36–39^ The biosynthesis of the β-amino fatty acid is thought to require a free-standing A domain, a free-standing PCP domain, and a free-standing glycine oxidase to incorporate the amino group from glycine into an - unsaturated fatty acyl precursor.^36,37^ However, we have not yet identified a candidate glycine oxidase in the worm genome, and thus, it is unclear whether the amino group in the nemamides is incorporated in an analogous fashion.

**Figure 6.**
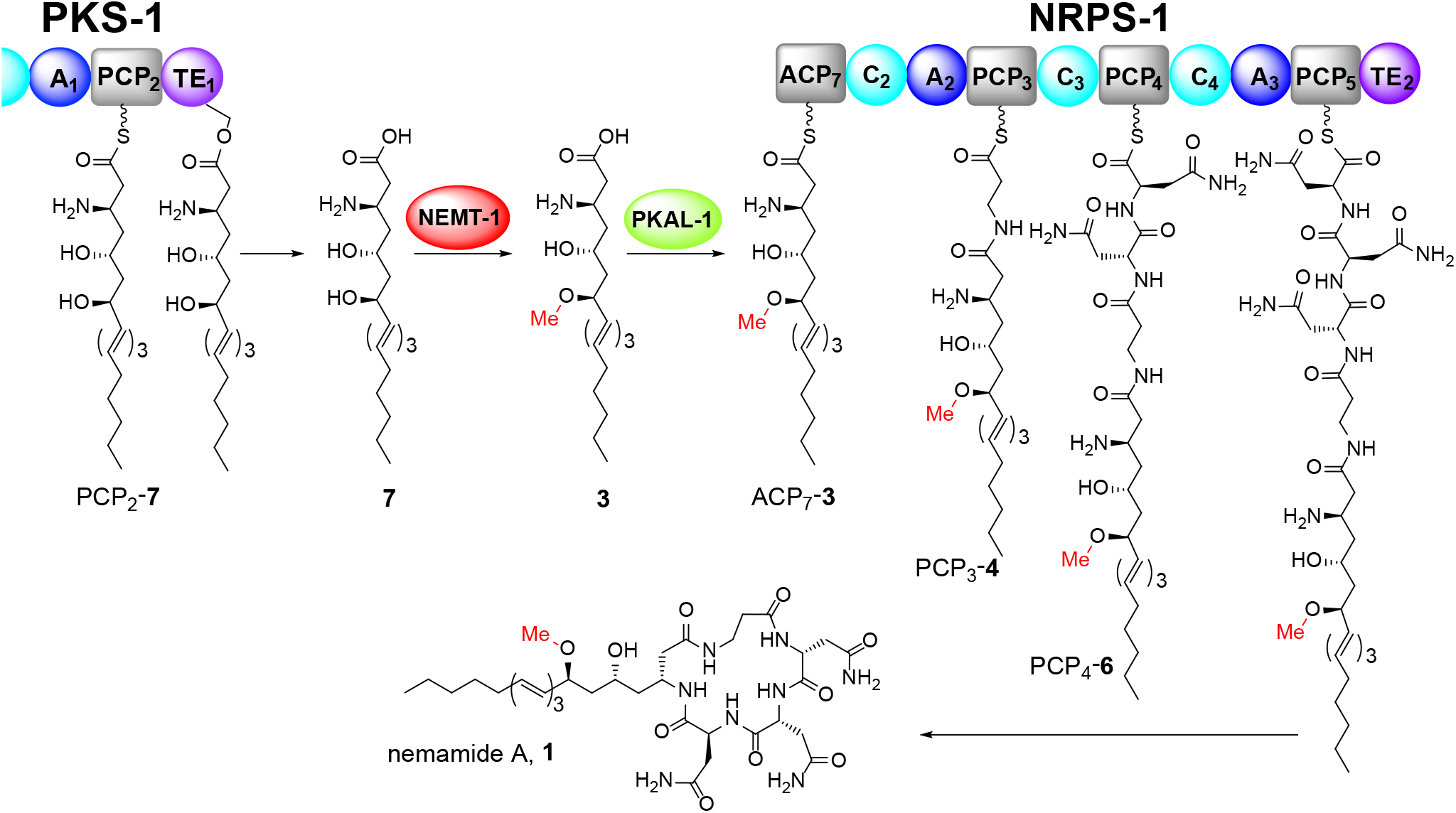
Proposed model for the biosynthesis of the nemamides. Intermediate **7** is released from PKS-1 and methylated by NEMT-1. The methylated intermediate **3** is activated by PKAL-1 as the adenylate and loaded onto the ACP_7_ domain of NRPS-1 in order to biosynthesize nemamide A, **1**. An analogous pathway biosynthesizes nemamide B, **2**.

Trafficking between PKS and NRPS enzymes typically involves protein-protein interactions between the megasynthetases that enable direct passage of intermediates from the carrier domain on an upstream PKS or NRPS to the KS or C domain on the downstream PKS or NRPS, respectively.^16^ Here, we have shown that trafficking of intermediates between PKS-1 and NRPS-1 in the biosynthesis of the nemamides in *C. elegans* involves reloading of a complex polyketide intermediate. According to our model, NEMT-1 methylates the polyketide product of PKS-1 (e.g., **7**), and PKAL-1 activates these *O*-methylated intermediates (e.g., **3**) and loads them onto NRPS-1 to subsequently produce the nemamides (**Fig. 6**). Although it is possible that NEMT-1 acts earlier in the pathway and methylates an intermediate attached to PKS-1, we do not favor this model over the one shown in **Figure 6** for the following reasons: (1) The *nemt1*-*1* mutant strain is able to produce desmethyl-nemamide A, suggesting that PKAL-1 and NRPS-1 can process desmethyl intermediates, reducing the likelihood that intermediate **7** is simply accumulating because PKAL-1 and NRPS-1 cannot process it; (2) NEMT-1 is most similar to *O*-methyltransferases that act on free (non-protein bound) lipids (i.e., phthiotriol methyltransferase);^26^ (3) To our knowledge, all known examples of free-standing *O*-methyltransferases are thought to act as tailoring enzymes after natural products have been released from the assembly line. The mechanism proposed in **Figure 6** may be important for trafficking of intermediates in nemamide biosynthesis across membranes or between cellular compartments. Indeed, the C-terminus of PKS-1 has predicted transmembrane domains that may localize it to specific membranes and/or compartments.

The PKAL-1 enzyme is unique in that it activates an advanced polyketide intermediate as the adenylate and loads it onto ACP of the next megasynthetase in the assembly line, NRPS-1. There are many examples of FAAL enzymes activating fatty acids as AMP-esters for loading onto a carrier protein for extension by either a PKS or NRPS.^27,28,30^ However, to our knowledge, PKAL-1 is unique in that it activates an advanced polyketide substrate and loads it onto a carrier protein for further assembly-line processing on an NRPS. Furthermore, PKAL-1 is unusual in that it does not contain the insertion motif that characterizes most FAAL enzymes.^27–32^ Thus, other structural features, such as an inability to bind CoA, must explain its failure to act as a CoA ligase. Interestingly, PKAL-1 is upregulated under starvation conditions in a manner that is independent of the transcription factor DAF-16, which promotes survival during starvation-induce larval arrest.^40,41^ Previously, we showed that the nemamides promote survival during starvation-induced larval arrest in a DAF-16-independent manner and bacterial food suppresses the production of the nemamides.^8^ Thus, the requirement of PKAL-1 for nemamide production may help to couple nemamide production to food availability.

In summary, we have shown that nemamide biosynthesis requires at least seven different genes expressed in the CANs. Two of these genes encode enzymes that are required for the trafficking of intermediates between PKS-1 and NRPS-1, and this unique mechanism may facilitate the regulation of nemamide biosynthesis.

## Acknowledgements

We thank Patrick McGrath and Yi Sheng for kindly providing plasmids for CRISPR-Cas9, and Steven Bruner for providing the pACYCDuet-sfp plasmid. We thank Qingyao Shou for advice and Manasi Kamat for help with mass spectrometry. Mass spectrometry analysis was performed at the University of Florida’s Mass Spectrometry Research and Education Center, which is supported in part by the NIH (S10 OD021758-01A1). We acknowledge the CGC, which is supported by the NIH Office of Research Infrastructure Programs (P40 OD010440), for providing some worm strains. This work was supported by the NIH (GM118775) and the NSF (Career-1555050).

## Author Contributions

R.A.B. and L.F. designed the study, analyzed the data and wrote the paper, which was reviewed by all authors; L.F. and M.T.G. performed experiments; L.F., Y.L. and K.B.B. performed LCMS/MS analysis.

## Online Methods

### Worm strains

Worms were maintained on OP50 using standard methods. Strains used in this study were obtained from Caenorhabditis Genetics Center or were generated through genome editing by CRISPR-Cas9 or through transgenesis (**Supplementary Table 2**). Some strains were backcrossed with wild-type strain (N2). The double mutants were generated from single mutants, using standard genetic crossing methods. The presence of alleles was verified through single worm PCR using primers in **Supplementary Table 3** and/or followed by restriction site digestion, as shown in **Supplementary Table 4**.

### Single worm PCR and CRISPR-Cas9

Some mutants containing deletions and point mutations were generated based on the Fire lab’s CRISPR-Cas9 protocol with modifications^42,43^. Concentration of Cas9 vector used was 50 ng/μL. The plasmid for expressing the *dpy*-*10* sgRNA was used at 25 ng/μL, and the other plasmids for expressing the sgRNA sequences for target genes were used at 50-100 ng/μL (see **Supplementary Table 5** for sequences). *dpy*-*10(cn64)* donor oligonucleotides were used at a concentration of 500 nM, and other donor oligos for generating desired mutations were used at 500-750 nM (see **Supplementary Table 6** for sequences). To generate deletion mutants, no donor oligonucleotide was used for either *dpy*-*10* or targeted genes. After injection, F1 worms with *dpy* (for deletions) and *roller* (for point mutations) phenotypes were picked for single worm PCR using the primers listed in **Supplementary Table 3** and restriction digestion (only for mutants with point mutations) of the PCR products listed in **Supplementary Table 4**. The PCR products of positive candidates were sequenced to confirm their alleles, and *dpy* worms were backcrossed with wild-type worms.

### Transgenesis

Translational reporters for each gene containing gene promoter plus genomic DNA were inserted into pBS77-SL2-mCherry using primers containing restriction sites listed in **Supplementary Table 7**. The translational reporter plasmids were injected into the corresponding mutant worm strains at 50 ng/μL. The *coel::dsred* plasmid was used as a coinjection marker at 25 ng/μL, and the total concentration of DNA injected was 100 ng/μL including pUC18. At least three independent transgenic lines were analyzed.

### Small-scale worm extraction for nemamide production

*C. elegans* wild-type and mutant worm strains were grown at room temperature on two NGM agar plates (10 cm) spread with 0.75 mL 25X OP50 until the food on the plates was almost gone. Then, the worms were transferred to 1 L Erlenmeyer flasks containing S medium (350 mL). The worm cultures were grown at 22.5°C for 3-5 d until no food was left and were fed with 3.5 mL of 25X OP50 every day. For sample collection, the culture flasks were placed in an ice-bath for 30 min to 1 h to settle the worms. Then, the worms were transferred from the bottom of the flasks to a 50 mL centrifuge tube and were centrifuged (1000 rpm for 5 min) to separate the worms from the worm medium. The process was repeated until most of the worms were removed from the flasks. The collected worms were washed with water three times and centrifuged (1000 rpm for 5 min), and then they were soaked in 10 mL of water for 1 h in a shaking incubator (22.5°C, 225 rpm) to remove bacteria from their digestive tract. The worms were collected by centrifugation and were freeze-dried. The dried worm pellets were ground with sea sand (2 g sand per 200 mg dried worms) using a mortar and pestle. The ground worms were extracted with 15 mL of 190 proof ethanol for 3.5 h, and the extract was centrifuged (3500 rpm for 20 min). The supernatant was collected and dried using a speedvac. The dried worm samples were each resuspended in 100 μL of methanol, sonicated (if needed), and centrifuged (15000 rpm for 1 min) before analysis by LCMS. The samples were analyzed using a Luna 5μm C18 (2) column (100 × 4.6 mm; Phenomenex) coupled with an Agilent 6130 single quad mass spectrometer operating in single ion monitoring (SIM) mode for nemamide A ([M+Na]^+^,*m/z* 757) and nemamide B (([M+Na]^+^, *m/z* 755). The following solvent gradient was used with a flow rate of 0.7 mL/min: 95% buffer A, 5% buffer B, 0 min; 0% buffer A, 100% buffer B, 20 min; 0% buffer A, 100% buffer B, 22min; 95% buffer A, 5% buffer B, 23 min; 95% buffer A, 5% buffer B, 26 min (buffer A, water with 0.1% formic acid; buffer B, acetonitrile with 0.1% formic acid). Two *pkal*-*1* mutants, RAB58 (*reb21* allele) and RAB59 (*reb28* allele), were analyzed in small scale, and neither of them produced any nemamides. Two C32E8.6 mutants, RAB60 (*reb23* allele) and RAB61 (*reb24* allele), were also analyzed, and neither produced any nemamides.

### Large-scale worm extraction for biosynthetic intermediates

The extraction method was similar to one previously reported but with modifications.^8^ 50 mL of wild-type or mutant strain cultures were inoculated and shaken at 180 rpm for 7-10 d at 20 °C in 2.8 L baffled flasks containing 500 mL of CeHR medium with 20% cow’s milk.^44^ Worms were collected by centrifugation, washed with water, shaken in water for 30 min to clear their intestines, and washed again with water. Worms were stored frozen at −20 Large-scale worm extraction for biosynthetic intermediatesC until needed. For the extraction and fractionation process, worms from 2 L-worth of culture (around 5-10 g of dried worm) were processed at a time. After freeze drying, worms were ground for 15 min with 50-100 g of sand using a mortar and pestle. The pulverized worms were transferred to a 1 L Erlenmeyer flask, and 300 mL of 190-proof ethanol was added to the flask. The flask was shaken at 300 rpm for 3.5 h. The extract was filtered using a Buchner funnel and filter paper and evaporated with a rotavap at 25 βC. The extract was then subjected to silica gel chromatography (50-100 g silica gel) and eluted with a gradient of hexane, ethyl acetate, ethyl acetate/methanol (1:1) and methanol (350 mL each) to give four fractions (A – D). Fractions C and D were processed separately, evaporated with a rotavap at 27 βC, redissolved in 12 mL of methanol, and centrifuged at 3500 rpm for 10 min. The supernatant was dried and dissolved in 10 mL 70% methanol/water. The resulting cloudy sample was then applied to an HP-20ss column (100 g HP-20ss resin), eluting with methanol/water (7:3 to 9:1 to 10:0) to give twelve subfractions (1 to 12, 125 mL each). Each fraction was dried by rotavap and analyzed by LC-MS for peaks with the same characteristic triene/tetraene UV spectrum found in the nemamides. The amount of nemamide or intermediate was calculated based on the total UV area of HPLC traces (UV at 280 nm for nemamide A-like molecules and UV at 320 nm for nemamide B-like molecules), normalized by sample dilutions and the total amount (g) of dried worms used for extraction. UV peaks were verified by masses in both positive and negative modes. Percentage for each compound in each sample was compared to the mean amount of nemamide A or nemamide B in wild type. *pkal*-*1* or C32E8.6 mutants containing different alleles were analyzed, and they showed very similar intermediate patterns. For the *pkal*-*1* mutant, the *reb28* allele was used for most analyses of single and double mutants. For the C32E8.6 mutant, the *reb23* allele was used for most analyses of single mutants. At least two or three independent experiments were performed for each worm strain. Extracts above were analyzed using a Luna 5μm C18 (2) column (100 × 4.6 mm; Phenomenex) coupled with an Agilent 6130 single quad mass spectrometer operating in both positive and negative mode under full scan and single ion monitoring (SIM) modes. The following solvent gradient was used with a flow rate of 0.7 mL/min: 95% buffer A, 5% buffer B, 0 min; 0% buffer A, 100% buffer B, 20 min; 0% buffer A, 100% buffer B, 22min; 95% buffer A, 5% buffer B, 23 min; 95% buffer A, 5% buffer B, 26 min (buffer A, water with 0.1% formic acid; buffer B, acetonitrile with 0.1% formic acid). Intermediate **3**, ESI (*m/z*): [M+H]^+^ 340, [M-OCH4+H]^+^ 308, [M-H]^−^ 338; Intermediate **4**, ESI (*m/z*): [M+H]^+^ 411, [M-OCH4+H]^+^ 379, [M-H]^−^ 409; Intermediate **5**, ESI (*m/z*): [M+H]^+^ 525, [M-H]^−^ 523; Intermediate **6**, ESI (*m/z*): [M+H]^+^ 639, [M-H]^−^ 637; Intermediate **7**, ESI (*m/z*): [M+H]^+^ 326, [M-OCH4+H]^+^ 308, [M-H]^−^ 324; desmethyl-nemamide A, **8**, ESI (*m/z*): [M+H]^+^ 743, [M+H]^+^ 721, [M-OCH4+H]^+^ 703, [M-H]^−^ 719.

### Identification of nemamide biosynthetic intermediates by qTOF

Extracted nemamide biosynthetic intermediates were analyzed by HR-LC-MS and HR-LC-MS/MS in positive ion mode. 5 μL of sample was analyzed on a Capillary LCMS Solutions 3 μm 200Å (0.3 × 150 mm) ProtoSIL C18AQ+ column attached to an UltiMate 3000 capillary RSLC System and a Bruker Impact II QTOF mass spectrometer equipped with an Apollo II ion funnel ESI source (Bruker) operated in positive ion mode with the following settings: mass range was *m/z* 80 – 1300 at a rate of 1 Hz, capillary voltage was 2.5 kV, source temperature 200 °C, drying gas 4.0 L/min, and nebulizer gas at 0.3 bar. Mobile phase A was water with 10 mM ammonium formate and 0.1% formic acid, and mobile phase B was acetonitrile with 0.1% formic acid. The following solvent gradient was used for separation with a flow rate of 5 μL / min: 98% buffer A, 2% buffer B, 0 min; 98% buffer A, 2% buffer B, 5 min; 40% buffer A, 60% buffer B, 35min; 5% buffer A, 95% buffer B, 55 min; 5% buffer A, 95% buffer B, 67 min; 98% buffer A, 2% buffer B, 70 min; 98% buffer A, 2% buffer B, 75 min. HR-LC-MS/MS was performed for each sample with a collision energy of 30 eV or 15 eV. Intermediate **3**, HR-ESIMS (*m/z*): [M+H]^+^ calcd. for C19H34NO4 340.2482, found 340.2483; intermediate **4**, HR-ESIMS (*m/z*): [M+H]^+^ calcd. for C22H39N2O5 411.2853, found 411.2849; intermediate **5**, HR-ESIMS (*m/z*): [M+H]^+^ calcd. for C26H45N4O7 525.3283, found 525.3281; intermediate **6**, HR-ESIMS (*m/z*): [M+H]^+^ calcd. for C30H51N6O9 639.3712, found 639.3716; intermediate **7**, HR-ESIMS (*m/z*): [M+H]^+^ calcd. for C18H32NO4 326.2326, found 326.2326; desmethyl-nemamide A, **8**, HR-ESIMS (*m/z*): [M+Na]^+^ calcd. for C33H52N8O10Na for 743.3699, found 743.3694, [M+H]^+^ calcd. for C33H53N8O10 for 721.3879, found 721.3879, [M-H2O+H]^+^ calcd. for C33H51N8O9 for 703.3774, found 703.3769.

### Plasmid construction, protein overexpression and purification

All genes and excised domains were amplified by PCR using Phusion polymerase (New England Biolabs) from a *C. elegans* cDNA library. Specifically, *pkal*-*1* was amplified with CATG<u>CCATGG</u>GGGCGAAATATTATCCAGAAAC and CATG<u>GCGGCCGC</u>ATAGTACATTAGCCTATTC, *pks*-*1_ACP_1_* was amplified with GCGC<u>CCATGG</u>GGCTTTCTGATGCGGAAATTGAGTC and CATG<u>GCGGCCGC</u>AGTTGTTGCTTTAGTAACTGGAAC, and *nrps*-*1_ACP_7_* was amplified with CATG<u>CCATGG</u>GGAGTGAAGACTCCGATGAAGAAGT and CATG<u>GCGGCCGC</u>TCCGGACCCCAGCGCTTTCTCAC. The genes were inserted using *Nco*I and *Not*I into the pET16b-KH01 vector (a modified version of pET-16b)^45^ such that they were expressed with C-terminal His tag. All of the sequences were verified by sequencing. The PKAL-1(K488A) mutation was generated via Q5 site-directed mutagenesis kit (New England Biolabs), using the primer pair GTCAAGTGGGGCCATTCAAAAGAATAG and GATTTTGGCATCTCTTTTATAATTG and verified through sequencing. Additional mutations were introduced into ACP_1_ and ACP_7_ to allow visualization of the carrier proteins by UV/vis, enabling purification by FPLC and concentration estimation by NanoDrop. The second and third residues (after the start codon) in ACP_1_ in were modified to Tyr and Trp, respectively, and the second residue (after the start codon) in ACP_7_ was modified to Trp and Tyr, via the Q5 site directed mutagenesis kit. Specifically, primers ATATACCATGTACTGGTCTGATGCGGAAATTGAG and CTCCTTCTTAAAGTTAAACAAAATTATTTC were used to mutate ACP_1_, and primers GTACAGTGAAGACTCCGATGAAG and CACATATATCTCCTTCTTAAAGTTAAAC were used to mutate ACP_7_. The PKAL-1 construct was transformed into BL21 (DE3) cells, and the cells were grown in LB broth with 150 mg/L ampicillin at 37 °C to OD600 0.6-0.8, cooled down on ice for 20 min, and induced with 0.3 mM IPTG at 16 °C for 20 h. The ACP_1_ and ACP_7_ constructs were each co-transformed with pACYCDuet-sfp into BL21 (DE3) for co-expression. The cells were grown in LB broth with 150 mg/L ampicillin and 34 mg/L chloramphenicol at 37 °C to OD600 0.3-0.4, and temperature was lowered to 16 °C for expression. 30 min prior to induction, cultures were supplemented with 2.5mM calcium pantothenate, and once cells reached OD600 0.6-0.8, protein expression was induced with 0.6 mM IPTG at 16 °C for 20 h. All purification steps were carried out at 4 °C. Briefly, cells were collected by centrifugation at 3700 rpm for 10 min, and resuspended in lysis buffer (20 mM Tris, 500 mM NaCl, pH 7.5). The cells were then lysed by microfluidizer three times and centrifuged at 18,000 rpm for 20 min. The supernatant was incubated with 1 mL of pre-equilibrated Nickel-resin (Thermo Scientific) for 1 h by shaking on ice. The resin was washed with 15 mL of lysis buffer, 15 mL of wash buffer (20 mM Tris, 500 mM NaCl, 20 mM imidazole, pH 7.5), and eluted with wash buffer containing 250 mM imidazole. For PKAL-1, the eluted sample was concentrated and loaded onto an FPLC connecting to a Superdex 200 gel filtration column (GE healthcare) with buffer (20 mM Tris, 100 mM NaCl, pH 7.5). Protein concentrations were determined by using Quick Start Dye reagent (Bio-Rad) with 2 mg/mL bovine serum albumin used as a standard (for PKAL-1) or by Nanodrop (for the carrier proteins). Purified proteins were flash frozen in 10% glycerol and stored at −80 °C. Mass analysis of the carrier proteins expressed individually compared to co-expression with Sfp showed complete conversion of the carrier proteins from the *apo* to the *holo* form.

### LC-MS-based PKAL-1 activity assay

To examine the activity of PKAL-1 and its catalytic mutant, a LC-MS-based assay was used as previously described.^46^ Generally, 50 μL reaction mixture contained 100 mM potassium phosphate at pH 7.0, 5 mM MgCl_2_, 5 mM CoA, 5 mM ATP, 0.1% Triton X-100, and 1 mM fatty acid substrate. The reaction was initiated by adding 2 μL of 2 mg/mL purified PKAL-1 or PKAL-1(K488A) enzyme at 25 °C for 2 h. 50 μL methanol was added to quench the reaction, and the reaction was vortexed and centrifuged. 5 μL supernatant was used for LC-MS analysis on an Agilent 6130 single quadrupole mass spectrometer in both positive and negative full-scan modes, mass range 150-1500, 125 V fragmentor voltage, 0.15 min peak width and 2.20 sec cycle length. Mobile phases A was water with 10 mM ammonium acetate, and mobile phase B was acetonitrile. The LC gradient was started from 95% A for 2 min and then ramped up to 100% B over 24 min.

### MALDI-TOF MS analysis

A 15 μL reaction mixture contained 100 mM Tris buffer at pH 7.8, 10 mM MgCl_2_, 1 mM TCEP, 1 mM ATP, 100 μM fatty acid substrate, and 100 μM *holo-*ACP_7_ or 100 μM *holo*-ACP_1_. The reactions were initiated by adding 2 μL of 2 mg/mL purified PKAL-1 or PKAL-1(K488A) at 25 °C for 2 h. Samples were diluted 1:10 in ultrapure water and spotted onto a ground plate 1:1 with a matrix containing saturated sinapinic acid in 70% acetonitrile. A Bruker AutoFlex LRF MALDI-TOF (Bruker Daltonics, Bremen, Germany) equipped with a Smartbeam-II UV laser was used to analyze the ACP masses, using the positive linear mode at a mass range between 5,000-20,000 Da. Laser power was used at the threshold level required to generate signal until suitable data was obtained. The instrument was calibrated with Protein Calibration Standard II (Bruker Daltonics) bracketing the molecular weights of the samples (typically, mixtures of apo myoglobin and bovine serum albumin using doubly charged, singly charged, and dimer peaks, as appropriate). All data was analyzed using flexAnalysis software (Bruker). Each single spectrum was the average of 500 laser shots, and the final spectra were generated using the sum of at least 3 single spectra.

