## Supplementary Information for "Mapping the biosynthetic pathway of a hybrid polyketide-nonribosomal peptide in a metazoan"

**a** Thioesterase (TE) domains

```
PKS-1_TE1     .....DLKYPSNDLRELAHFYAEEIAA...HAGNKRIFVMGHSMGGIMSREIVAELKIWG
NRPS-1_TE2    ..........GDTVDEVAKLYRLQIEESAENIETSKLVFIGASSAGTFAFSTSQLFADDD
Pik_TE        GTGTGTALLPADLDTALDAQARAILR....AAGDAPVVLLGHSGGALLAHELAFRLERAH
Sur_TE        NQ.TS....AIEDLEELTDLYKQELN....LRPDRPFVLFGHSMGGMITFRLAQKLEREG
Ybt_TE        LRHLE....PLRSITQLAALLANELEA..SVSPDTPLLLAGHSMGAQVAFETCRLLEQRG
Rif_TE        RRHEP....PVDSIGGLTNRLLEVLR....PFGDRPLALFGHSMGAIIGYELALRMPEAG
                                                     *
```

**b** Condensation (C) domains

```
NRPS-1_C2     .............EKSSVSNPLSAI...QCSKLSPETLQLIFHHISIDGRSLAIFYQQFK
NRPS-1_C3     ..............PSGASRPKDQFSIQLWMSSKNKLLTISIHHLICDGRSLQILEHQLQ
NRPS-1_C4     IR.LLCDEPINV....LEGSPMIRASFI..SSPEKHVAFLHLHHLISDARSTQLTNSTMK
PKS-1_C1      .........NHLFE..IGKSTPLRVRVAEDCDNSRIHIVFNQHHILTDGWSMTVLSDTVS
ArfA-C2       FSARR....Y...RLDVSQAPLMRLVYARDPALDRVVGILLFHHLAMDHIALEV....MR
VibH_C        QIEQDLQRSSTLIDAPITSHQVYRLSHS......EHLIYTRAHHIVLDGYGMMLFEQRLS
CDA_C1        WMDRDRATPLPLDRPGLSSHALFTLGGG......RHLYYLGVHHIVIDGTSMALFYERLA
                                                         *    *
```

**c** Acyl-carrier protein (ACP) domains

```
PKS-1_ACP3    ....KTLQMAVRHKVCLAVGDVIESGLDIDESQLSTGFSE.LGIDSLATVDLLNRLNQKY
PKS-1_ACP4    ESDATVDRTEIRRKVSLAVFDLATETLSAEDLQ.SKGFTE.LGMDSLSIVDFVNRLNDKY
PKS-1_ACP6    ....KVKEEIKKKSLNFEEIFFEIVGITDISSKLNIPFMD.LGIDSLCMENLRYSLNKN.
Bac_ACP       ..........ADTLERVTKIIVDRLGVDEADVKLEASFKEDLGADSLDVVELVMELEDE.
PKS-1_ACP5    ...MNFSVEDEEEVLELIKEKVSSILMCSPTKLKNNKNIMDMGLDSKLIVEFLNFINST.
NRPS-1_ACP7   .....CTARPSRSMEILISILKDQMKLLSTSHEVETTPLPYLGIDSLRLAELEYHVASH.
PKS-1_ACP1    .NIVDEQTNSSLSDAEIESTVRTIVKQFLDIEEDDINLLETGAVDSLTSIEMVEAFGTAV
PKS-1_ACP2    .KITKKVENEDQKRASKNMLHVWFEENFGWTDIDNTTGFFDLGLTSIQAVKLRNAIKSN.
                                                            *
```

**d** Adenylation (A) domains

```
EntE_A        FFALLKLGVAPVL..ALFSHQRSELNAYASQIEPALLIADRQH....ALFSGDDFLNTFV
PKS-1_A1      LLACVFLGLPYAPIDPTWPEPRQLFVKSKVSFTLE....N...............CF...
SidN_A3       IVGIMKSGNTYVPIEAGLPNDRKSFLLRDSRAAMAFVCDN........NFDGVELPP...
Grs_A         ILAVLKAGGAYVPIDIEYPKERIQYILDDSQARMLLTQKHLVHLIHNIQFNGQVEIF...
NRPS-1_A2     VLAAWEAGLYPVPMHKDSKEAQIEKTLEALGIEEA.............FDSKDLCQ...
NRPS-1_A3     LAVQFT.GAAYLPIDASYPEERKKTILKDSVFNFE..............YNGRVDQE...

EntE_A        TEHSSIRVVQLLNDSGEHNLQDAINHPAEDFTATPSPADEVAYFQLSGGTTGTPKLIPRT
PKS-1_A1      ..........................SCNLKLRNFNSRTQFGSIYSIFTSGSTGVPKGVLMA
SidN_A3       .ET...KV...LD.TKNQSFIENLSTQDTSDILNNYPENLDAYLLYTSGSTGTPKGVRVS
Grs_A         .EE...DT...IK............IREGTNLHVPSKSTDLAYVIYTSGTTGNPKGTMLE
NRPS-1_A2     ............................TKLRIFNKSILYDLAYVTSTSGSTGTPKLVGTS
NRPS-1_A3     ............................PRHRHFAISTDYCLSYIITTSGTTGTPKSVAIG
                                                             *

EntE_A        HNDYYYSVRRSVE........ICQFTQQTRYLCAIPAAHNYAMSSPGSLGVFLAGGTVVL
PKS-1_A1      EQSVSSFMTSASK........QCMFRSNIRVLDSVKQVFD..VSVSNIIGSVLNGGVLIS
SidN_A3       RHNLSSFSDAWGKLIGNVAPKSLELGGVGKFLCLASRAFD..VHIGEMFLAWRFGLCAVT
Grs_A         HKGISNLKVFFEN........SLNVTEKDRIGQFASISFD..ASVWEMFMALLTGASLYI
NRPS-1_A2     FEGHSNLARQYTT........TYQISSRDTVGQVVDPSFD..IFFADIVK......TLVN
NRPS-1_A3     AKSLLNLFLSSTL........TMKCSSSSRTYQFTNFVFD..NSVLEVSMSIASQGTLVY
```

**e** Peptidyl-carrier protein (PCP) domains

```
NRPS-1_PCP3   .LPHPTLPNSLESDLSSIWTSLLNCPEPSPSDH.FFLIGGHSLLLVRLRHLIETKLGISL
NRPS-1_PCP1   KAAISIARLSLTSSLKHWLEHYSNLEIPSDDDD.IFTLGVDSIAVMLAMQKLRSEN.IEI
PKS-1_PCP2    KREIVVMKNSLEEKVINVFSKILGR.NVAPTDK.FESIGGNSLNAIQIAHRLAEELKIEI
NRPS-1_PCP2   KLLGLVEVRKKLELIIAAFKKFLDNTDVTKSTD.FFQAGGHSLTAMRLIDHLSDLLEVEI
Yer_PCP1      .AEADLPQGDIEKQVAALWQQLLSTGNVTRETD.FFQQGGDSLLATRLTGQLHQAG.YEA
PKS-1_PCP1    ESLKLPKSTSCEFVIAEIWKETLGISILNDANPNFFSLGGDSLSALQVVWKVQKKTDRIV
                                                     *
```

**Supplementary Figure 1. Domain sequence alignments of PKS-1 and NRPS-1 with known functional domains.** (**a**) Sequence alignment of TE domains with Pik_TE (Pikromycin TE domain, PDB ID: 2H7Y), Sur_TE (Surfactin TE domain, PDB ID: 2RON), Yer_TE (Yersiniabactin TE domain, PDB ID: 6BA8), and Rif_TE (Rifamycin TE domain, PDB ID: 3FLA). Sequences used for TE domains are PKS-1_$TE_1$ (7559-7610) and NRPS-1_$TE_2$ (2771-2820). The catalytic serine residue is labeled with an asterisk. (**b**) Sequence alignment of C domains with ArfA_$C_2$ (Arthrofactin module A $C_2$ domain), VibH_C (Vibriobactin free-standing C domain VibH, PDB ID: 1L5A), CDA_$C_1$ (Calcium-dependent antibiotic synthase $C_1$ domain, PDB ID: 4JN3). Sequences used for C domains are PKS-1_$C_1$ (6653-6701), NRPS-1_$C_2$ (520-563), NRPS-1_$C_3$ (1457-1502), and NRPS-1_$C_4$ (1895-1947). In the HHxxxDG motif, the second histidine (asterisk) serves as a catalytic base, and the aspartate (asterisk) is critical for the structural integrity of the active site. (**c**) Sequence alignment of NRPS-1_$ACP_7$ with PKS-1_ACP domains and *Bacillus subtilis* ACP (GenBank accession no. P80643, PDB ID: 1HY8). The conserved active site serine is marked with asterisk. Protein sequences used are PKS-1_$ACP_1$ (719-776), PKS-1_$ACP_2$ (1776-1833), PKS-1_$ACP_3$ (2792-2846), PKS-1_$ACP_4$ (2940-2997), PKS-1_$ACP_5$ (3471-3526), PKS-1_$ACP_6$ (5154-5207), and NRPS-1_$ACP_7$ (267-320). (**d**) Sequence alignment of A domains with EntE_A (Enterobactin module E, PDB ID: 3RG2), SidN_$A_3$ ($A_3$ domain in Siderophore N synthetase, PDB ID: 3ITE), Grs_A (Gramicidin S, PDB ID: 1AMU). Sequences used for A domains are PKS-1_$A_1$ (7044-7167), NRPS-1_$A_2$ (900-1019), and NRPS-1_$A_3$ (2274-2398). The conserved glycine residue in the flexible loop involved in the interaction with the pyrophosphate leaving group during amino acid loading is marked with an asterisk.[1] (**e**) Sequence alignment of PKS-1_PCP domains, NRPS-1_PCP domains, and Yer_$PCP_1$ (Yersiniabactin synthetase PCP1 domain, GenBank accession no. Q7CI41, PDB ID: 5U3H). The serine residue for phosphopantetheinyl postranslational modification is located in the conserved PCP motif DXFFXLGGDSL and is marked with an asterisk. Protein sequences are PKS-1_$PCP_1$ (6462-6521), PKS-1_$PCP_2$ (7424-7481), NRPS-1_$PCP_3$ (1289-1346), NRPS-1_$PCP_4$ (1700-1758), and NRPS-1_$PCP_5$ (2648-2705). Alignment was generated by Clustal Omega and ESPRipt 3.0.[2]

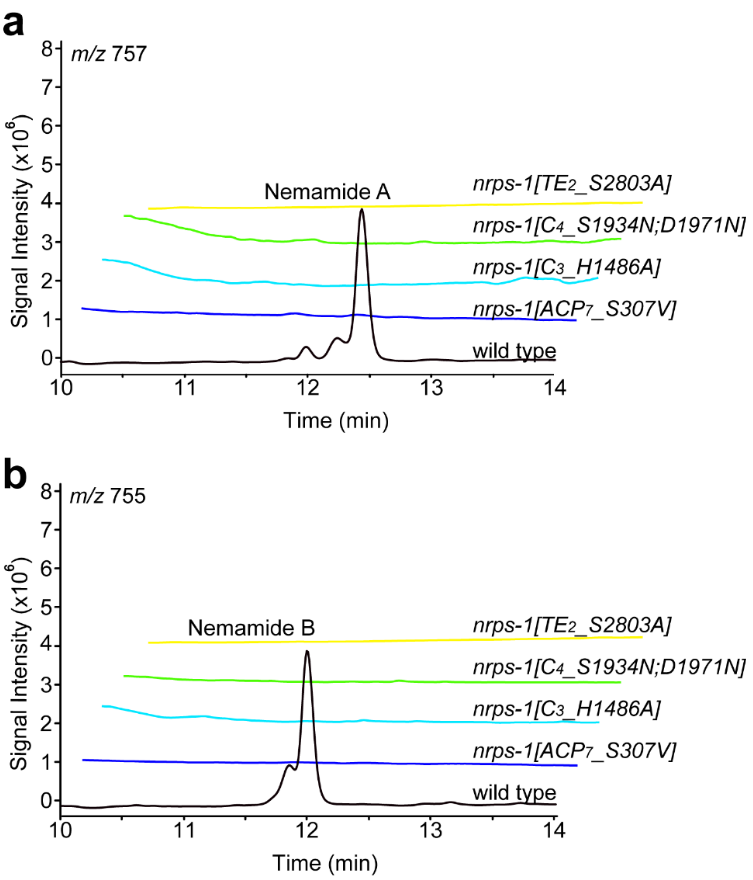


**Supplementary Figure 2. Nemamide production in wild-type and *nrps-1* domain mutants**. Extracted ion chromatograms for nemamide A (**a**) and nemamide B (**b**) in wild type, *nrps-1(reb12[TE$_2$_S2803A])*, *nrps-1(gk186409[C$_4$_S1934N]; gk186410[C$_4$_D1971N])*, *nrps-1(reb10[C$_3$_H1486A])*, and *nrps-1(reb8[ACP$_7$_S307V])* worms. The *nrps-1(gk186409[C$_4$_S1934N]; gk186410[C$_4$_D1971N])* mutant was obtained from the Caenorhabditis Genetics Center and backcrossed four times with wild type. Note that the retention times of the nemamides in the Supplementary Information are different than in the main text due to the fact that the samples were analyzed on different columns.

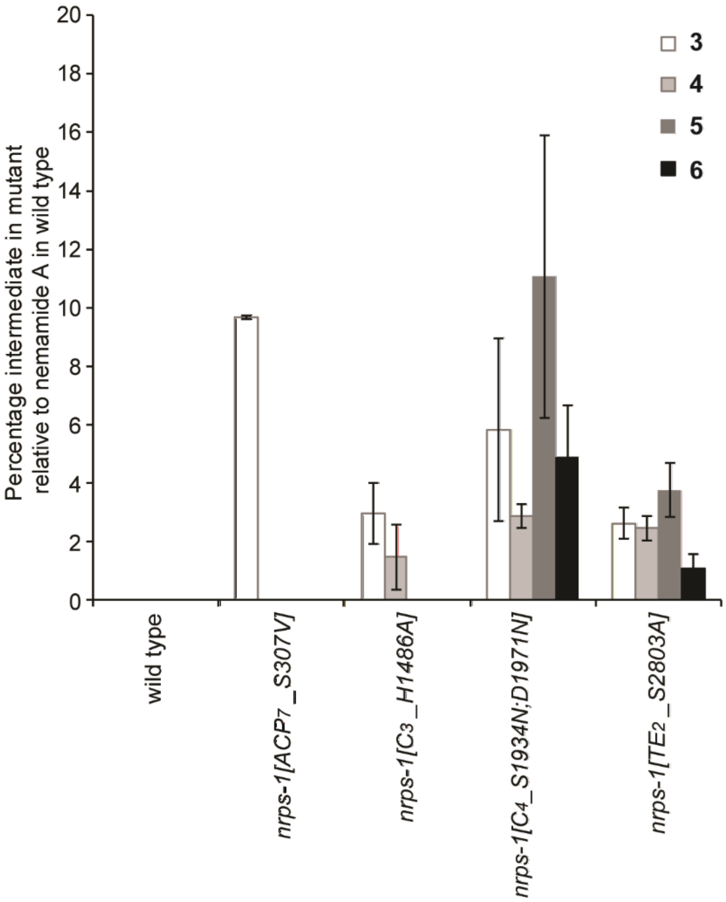


**Supplementary Figure 3. Production of intermediates in wild type and *nrps-1* domain mutants**. The production of intermediates **3**, **4**, **5**, and **6** in wild type, *nrps-1(reb12[TE$_2$_S2803A])*, *nrps-1(gk186409[C$_4$_S1934N]; gk186410[C$_4$_D1971N]), nrps-1(reb10[C$_3$_H1486A])*, and *nrps-1(reb8[ACP$_7$_S307V])* worms. The *nrps-1(gk186409[C$_4$_S1934N]; gk186410[C$_4$_D1971N])* mutant was obtained from the Caenorhabditis Genetics Center and backcrossed four times with wild type. Each strain was grown in large-scale culture for extraction of nemamide (in wild type) and intermediates (in mutants). Percentage for each intermediate in each strain was determined by comparing the amount of the intermediate (as gauged by UV at 280 nm) relative to the mean amount of nemamide A (as gauged by UV at 280 nm) in wild type. Data represent the mean ± standard deviation of at least two to three independent experiments.

**a**

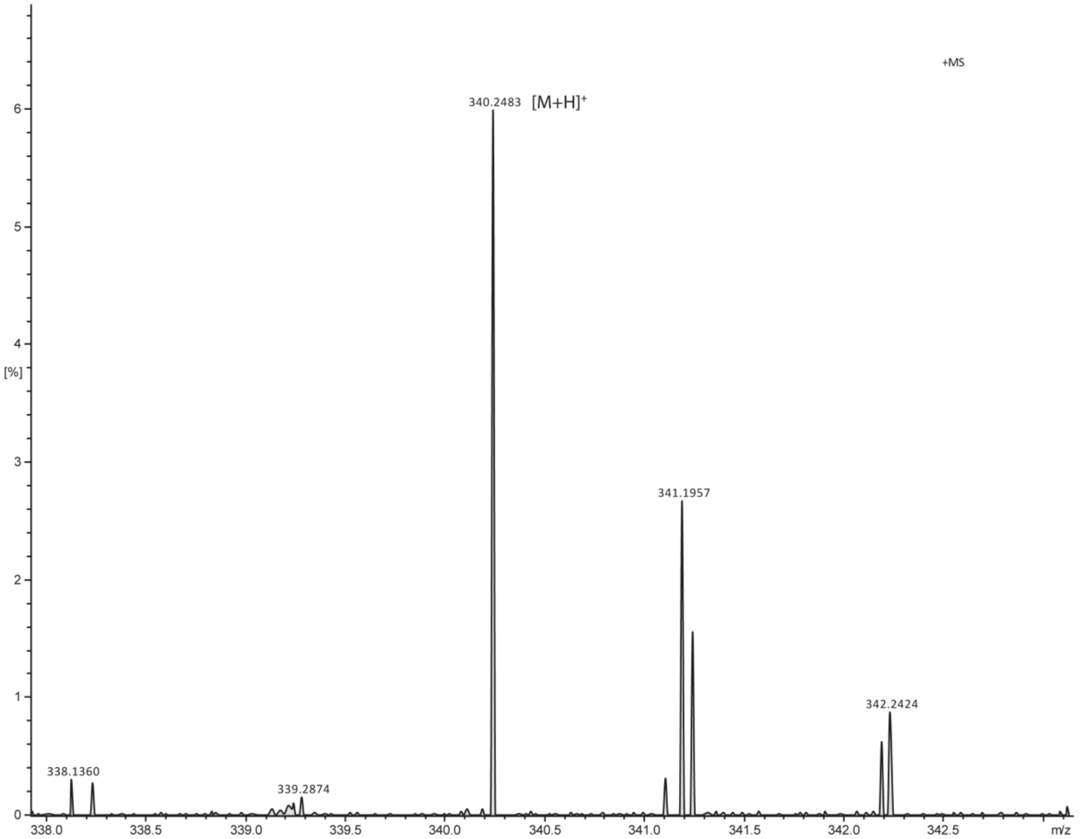
+MS
340.2483 [M+H]+
341.1957
342.2424
338.1360
339.2874
[%]
m/z

**b**

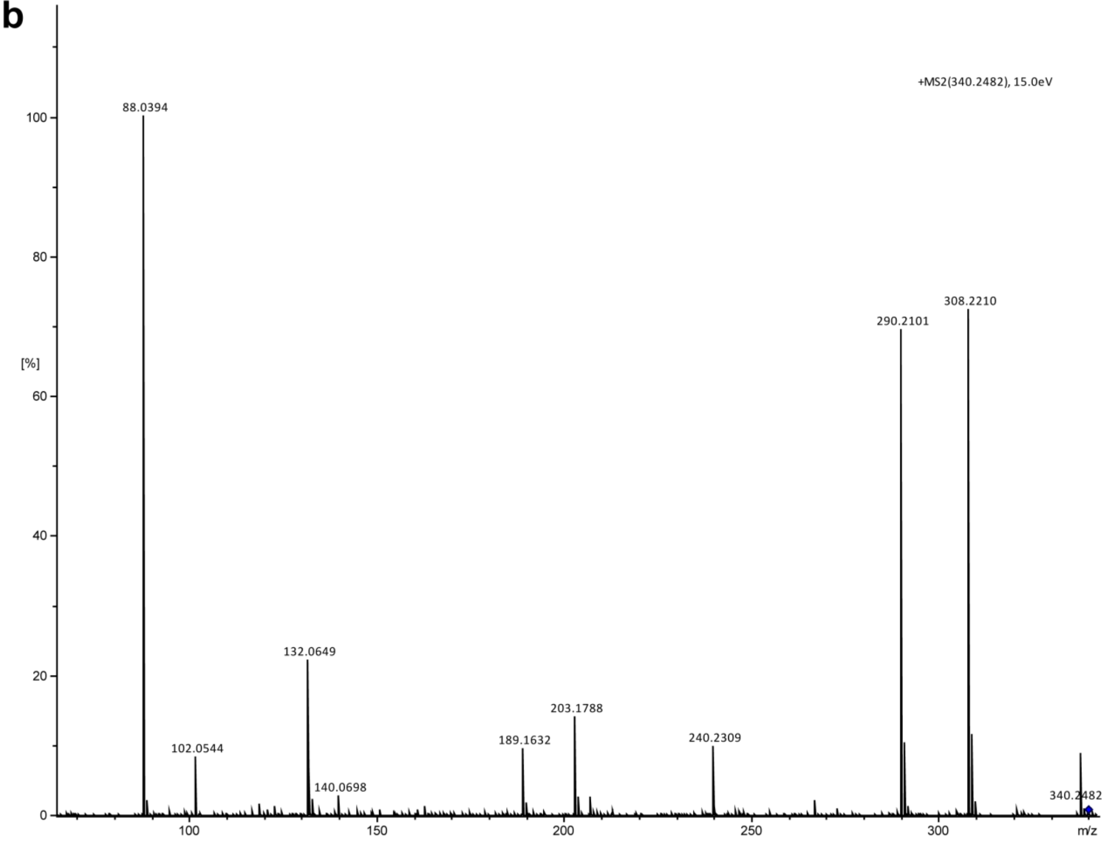
+MS2(340.2482), 15.0eV
88.0394
290.2101
308.2210
132.0649
102.0544
140.0698
189.1632
203.1788
240.2309
340.2482
[%]
m/z

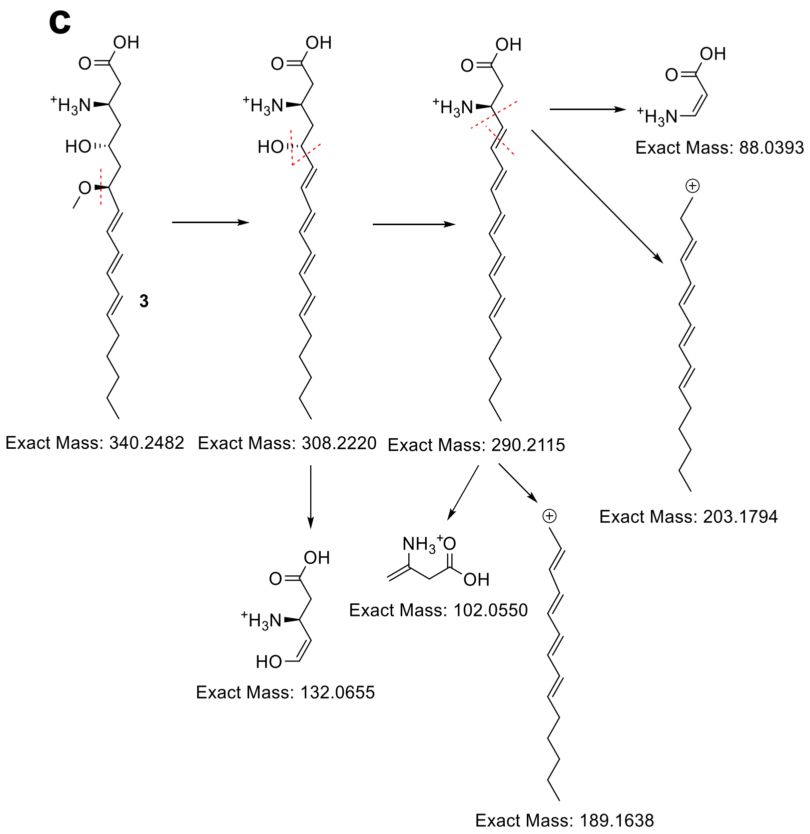


**Supplementary Figure 4. Mass spectrometry analysis of intermediate 3**. (**a**,**b**) HR-LC-MS/MS of intermediate **3**. (**c**) Analysis of the fragmentation patterns.

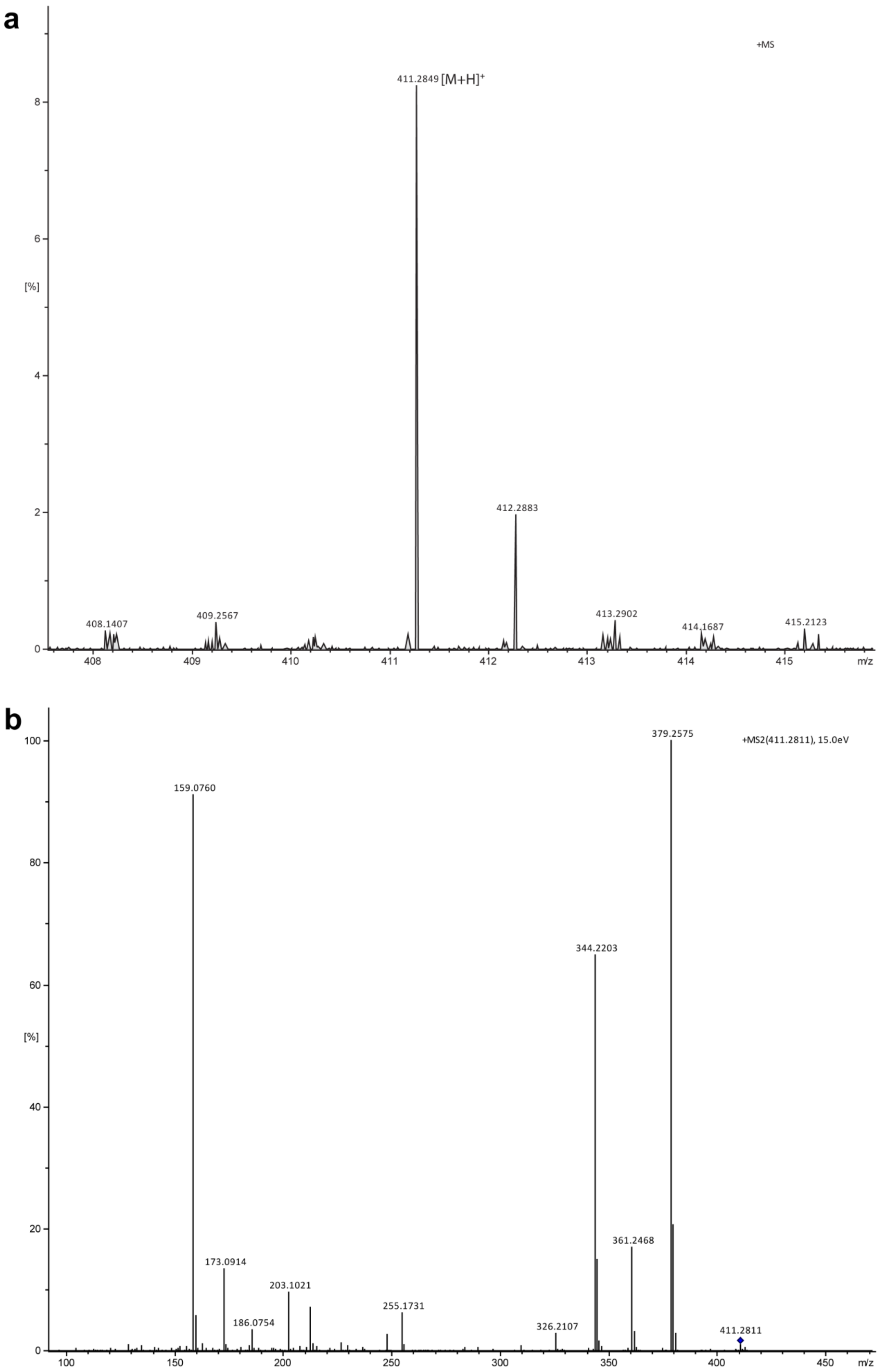

a
+MS
411.2849 [M+H]+
412.2883
408.1407
409.2567
413.2902
414.1687
415.2123
[%]
m/z
b
+MS2(411.2811), 15.0eV
379.2575
159.0760
344.2203
361.2468
173.0914
203.1021
186.0754
255.1731
326.2107
411.2811
[%]
m/z

c

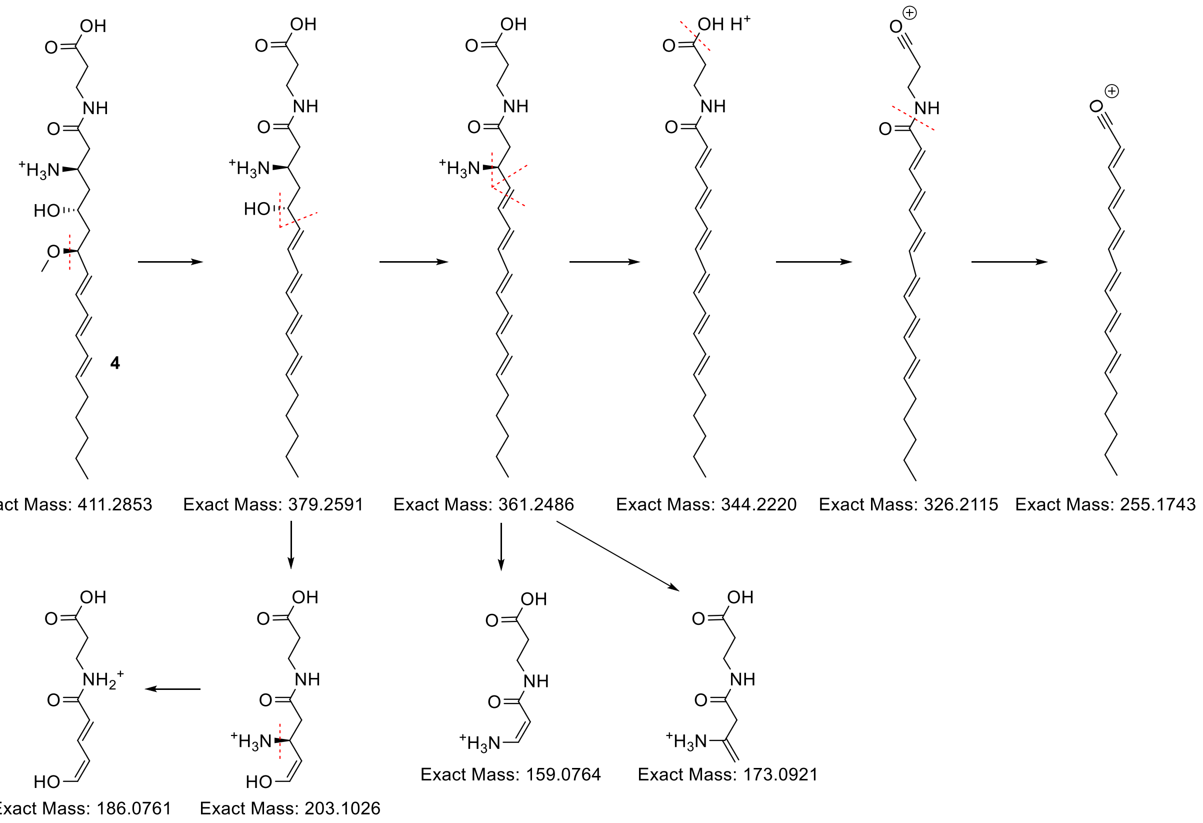


**Supplementary Figure 5. Mass spectrometry analysis of intermediate 4**. (**a**,**b**) HR-LC-MS/MS of intermediate **4**. (**c**) Analysis of the fragmentation patterns.

a

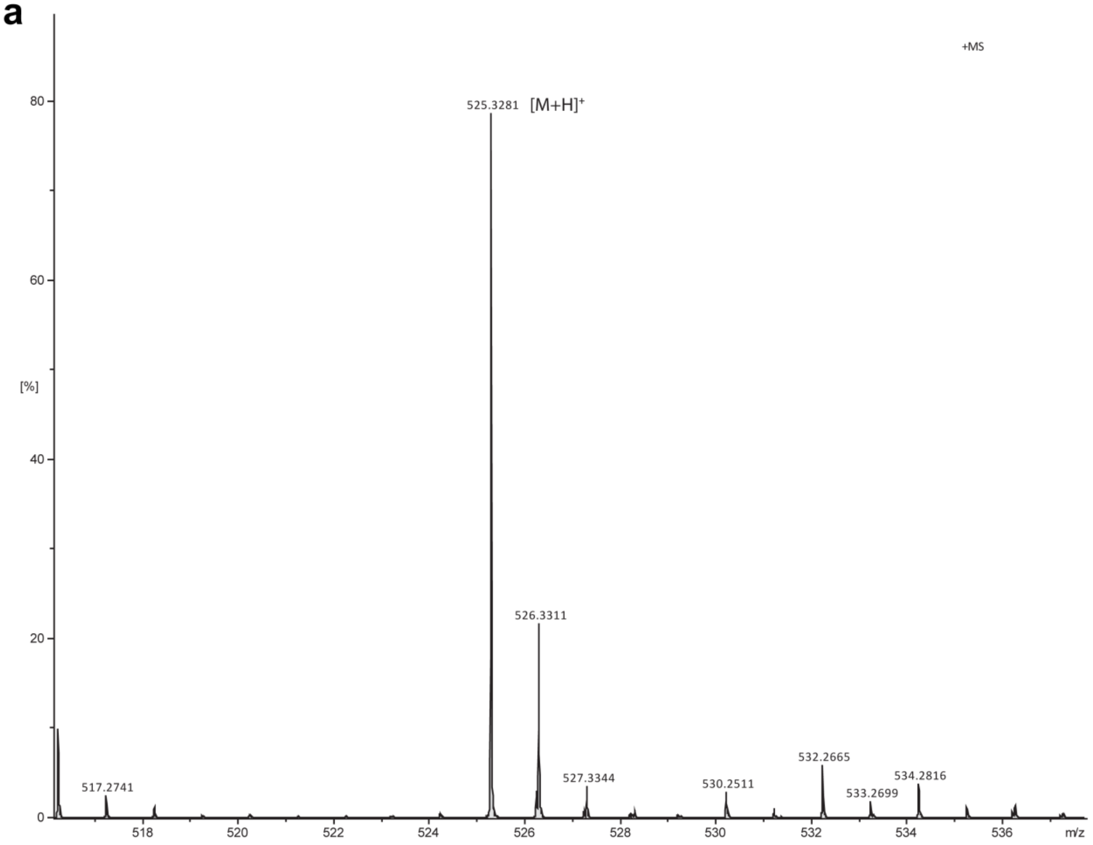
+MS
525.3281 [M+H]$^+$
526.3311
517.2741
527.3344
530.2511
532.2665
533.2699
534.2816
[%]
m/z

b

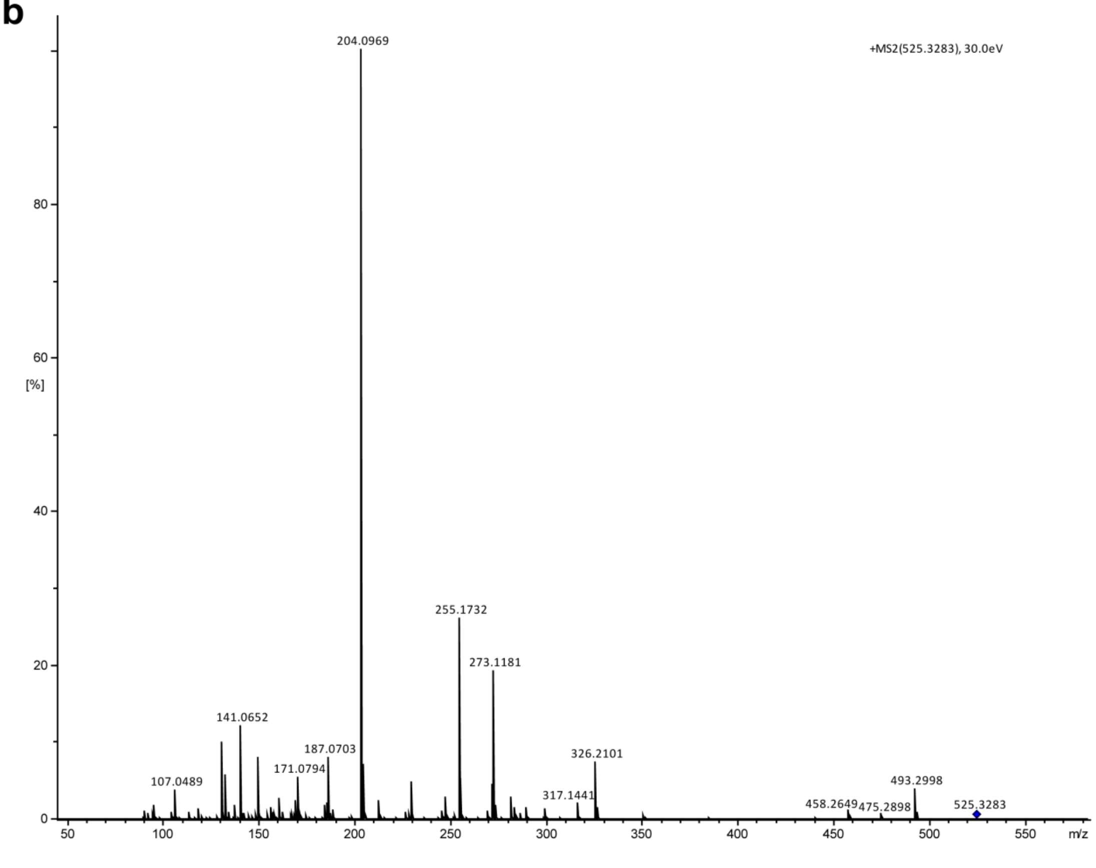
+MS2(525.3283), 30.0eV
204.0969
255.1732
273.1181
141.0652
187.0703
171.0794
107.0489
326.2101
317.1441
458.2649
475.2898
493.2998
525.3283
[%]
m/z

**c**

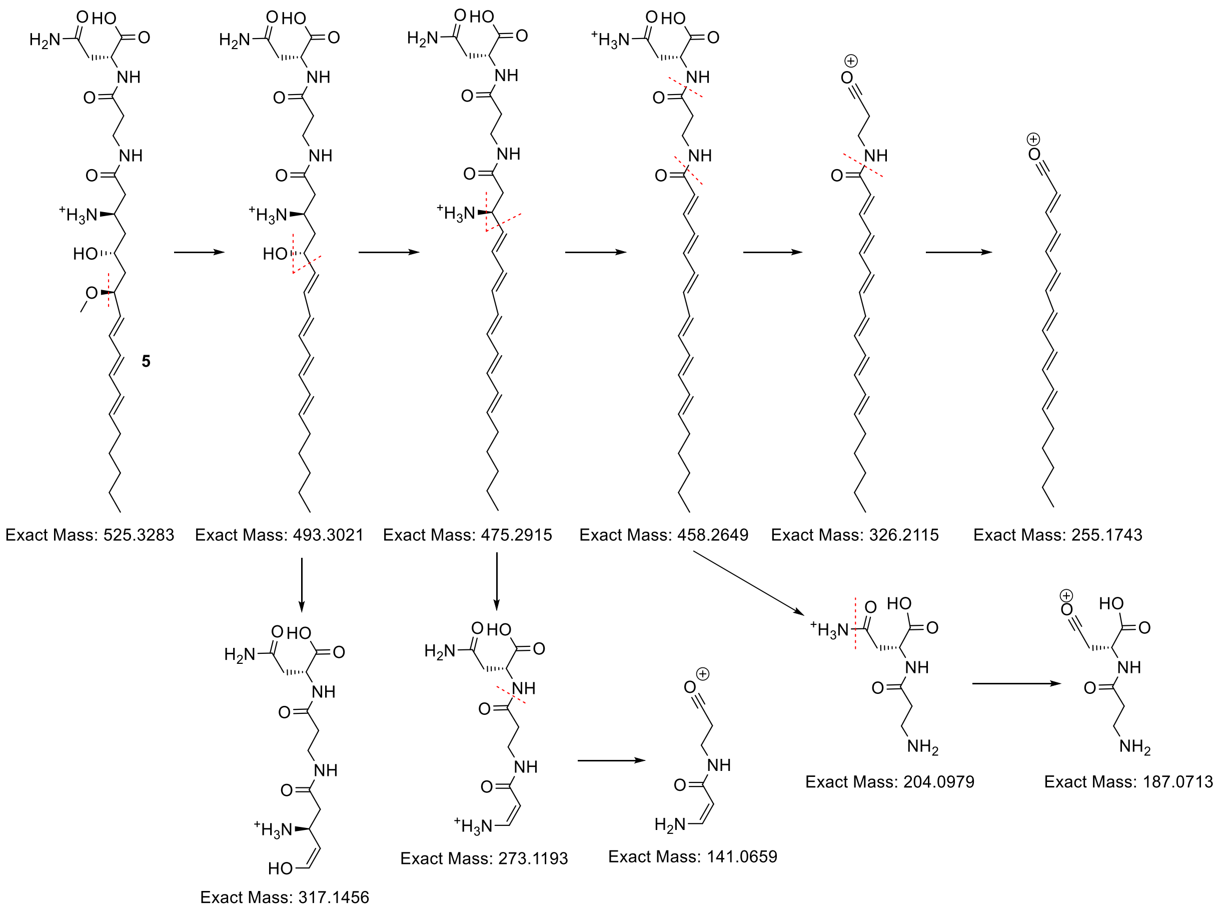


**Supplementary Figure 6. Mass spectrometry analysis of intermediate 5**. (**a**,**b**) HR-LC-MS/MS of intermediate **5**. (**c**) Analysis of the fragmentation patterns.

a

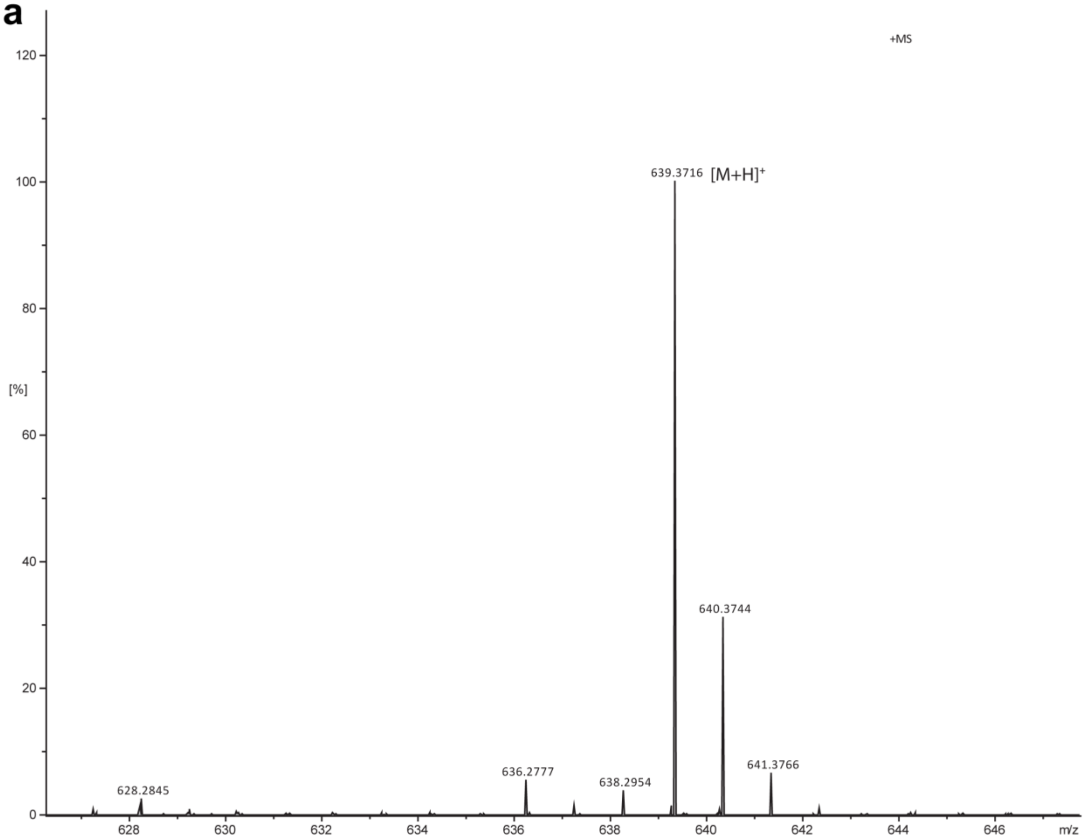

+MS
639.3716 [M+H]+
640.3744
641.3766
628.2845
636.2777
638.2954
[%]
m/z


b

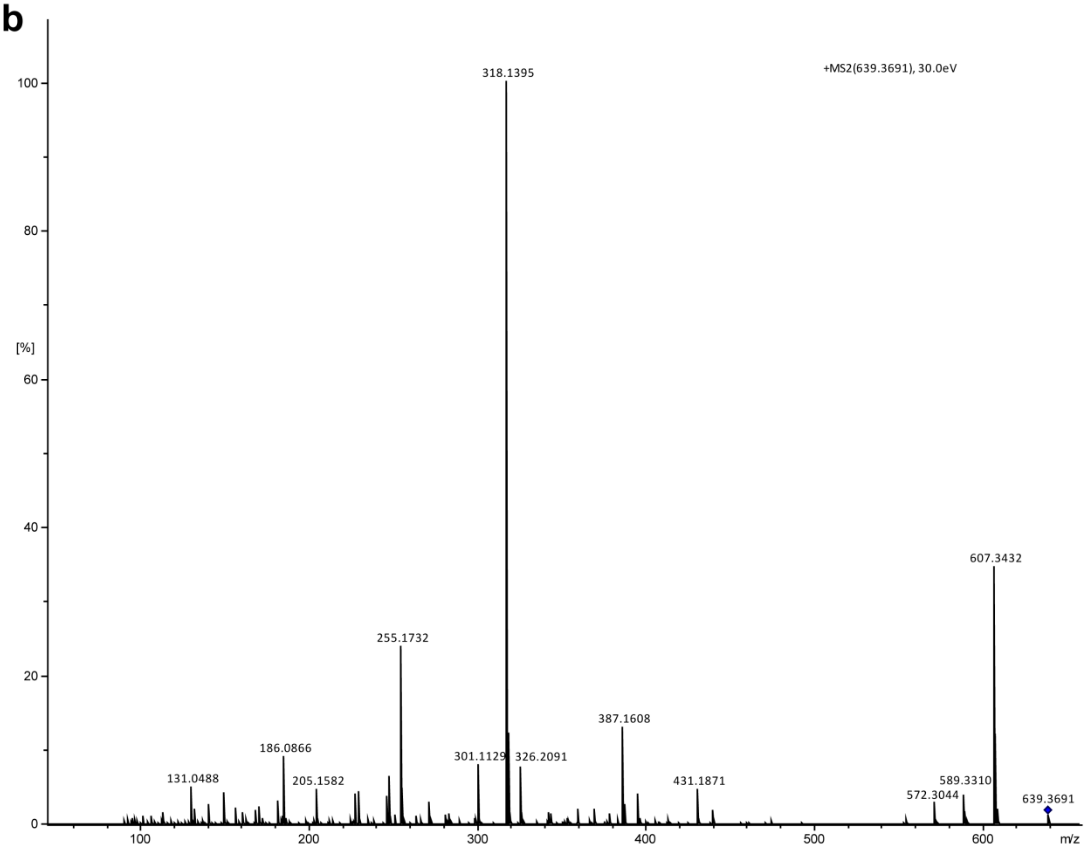

+MS2(639.3691), 30.0eV
318.1395
607.3432
255.1732
387.1608
186.0866
131.0488
205.1582
301.1129
326.2091
431.1871
572.3044
589.3310
639.3691
[%]
m/z

**c**

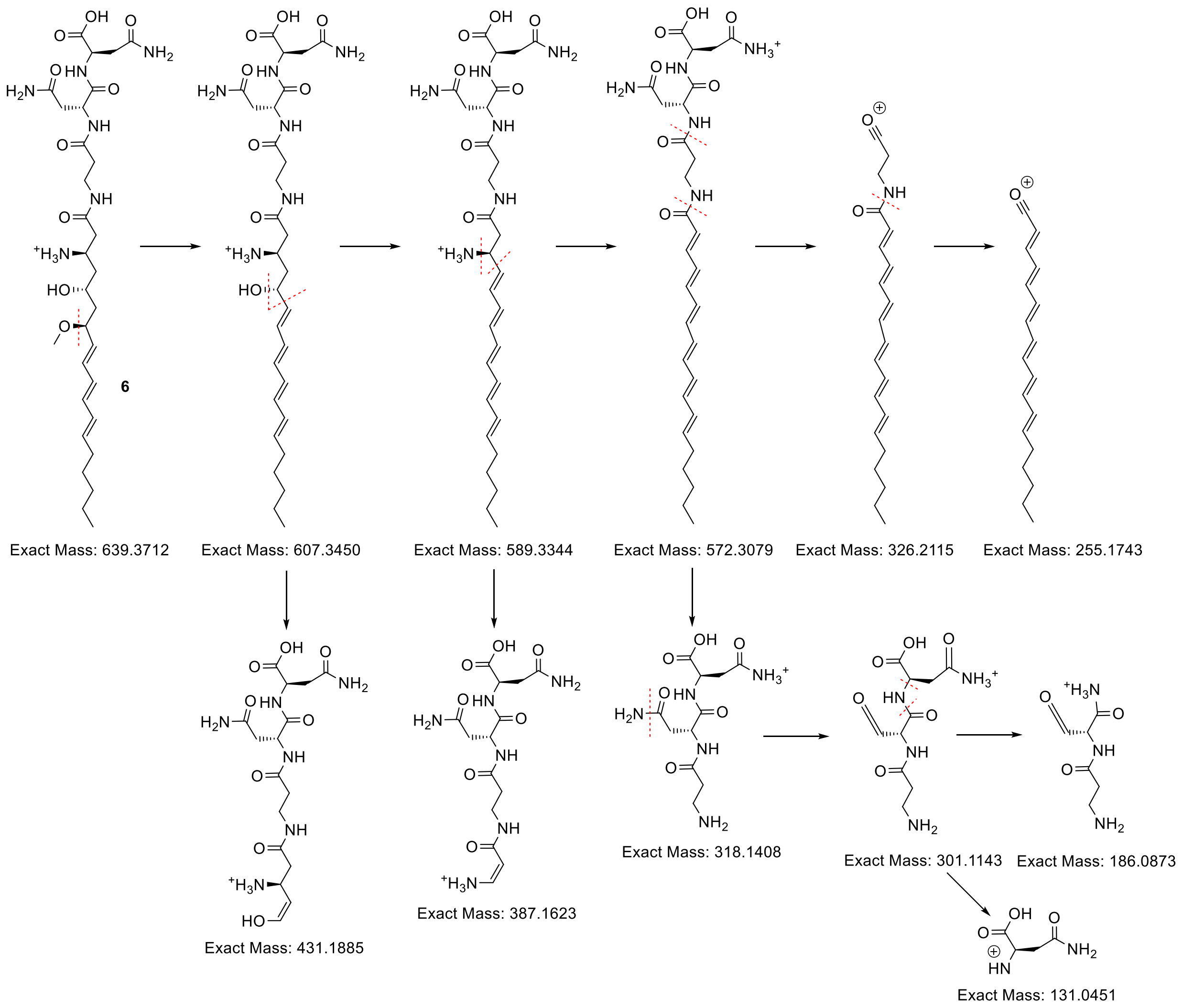


**Supplementary Figure 7. Mass spectrometry analysis of intermediate 6**. (**a**,**b**) HR-LC-MS/MS of intermediate **6**. (**c**) Analysis of the fragmentation patterns.

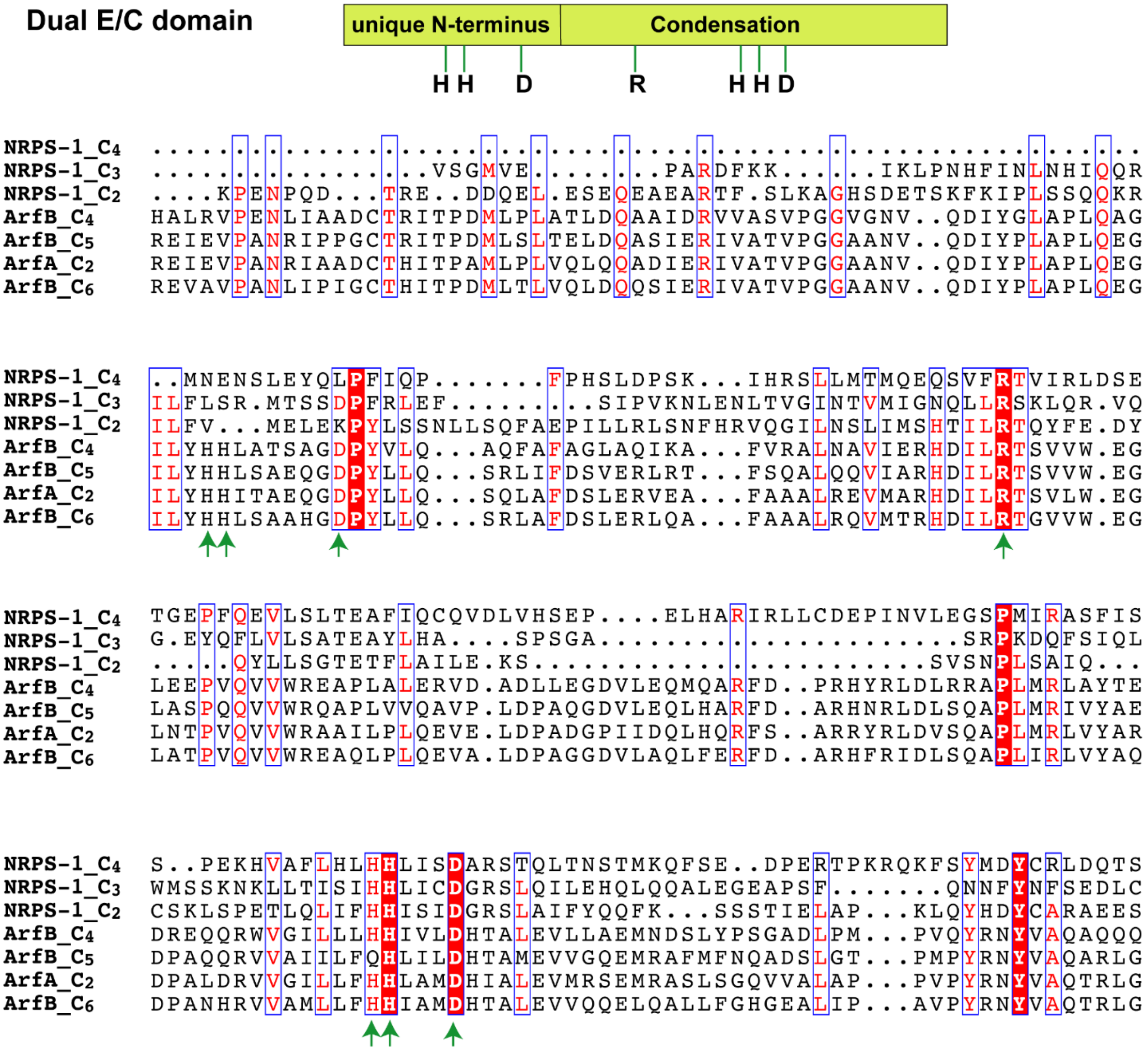


**Supplementary Figure 8. Domain sequence alignments of the C domains in NRPS-1 with the dual epimerase (E)/C domains from the arthrofactin biosynthetic pathway.** Dual E/C domains have a unique N-terminus with catalytic residues HH-D, followed by a common C domain with a catalytic motif R-HHXXXD.[3] The sequences of the $C_2$ domain from NRPS-1 (residues 401-586), the $C_3$ domain from NRPS-1 (residues 1361-1524), and the $C_4$ domain from NRPS-1 (residues 1815-1974) were compared to the sequences of the C domains of ArfB from the arthrofactin biosynthetic pathway. Essential residues for catalysis are indicated by arrows. The C domains in NRPS-1 are missing the catalytic residues in the N-terminus that are characteristic of dual E/C domains.

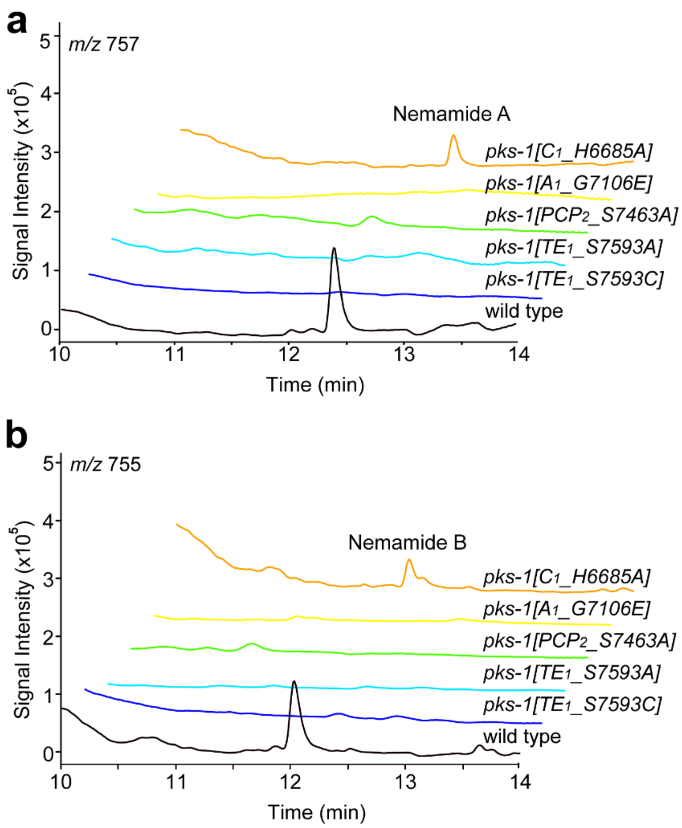


**Supplementary Figure 9. Nemamide production in wild type and *pks-1* mutant strains.** Extracted ion chromatogram for nemamide A (**a**) and nemamide B (**b**) in wild type and *pks-1(reb29[PCP$_2$_S7463A])*, *pks-1(reb22[A$_1$_G7106E])*, *pks-1(reb9[C$_1$_H6685A])*, *pks-1(reb11[TE$_1$_S7593A]),* and *pks-1(reb13[TE$_1$_S7593C])* mutant worms, containing mutations in the C-terminal NRPS module of PKS-1. Note that the nemamides could not be detected in the crude extracts of small-scale cultures of *pks-1(reb22[A$_1$_G7106E])* (as shown here), but they could be detected in very small amounts in the partially purified extracts of large-scale cultures of *pks-1(reb22[A$_1$_G7106E])* (as shown in **Fig. 3**). Note that the retention times of the nemamides in the Supplementary Information are different than in the main text due to the fact that the samples were analyzed on different columns.

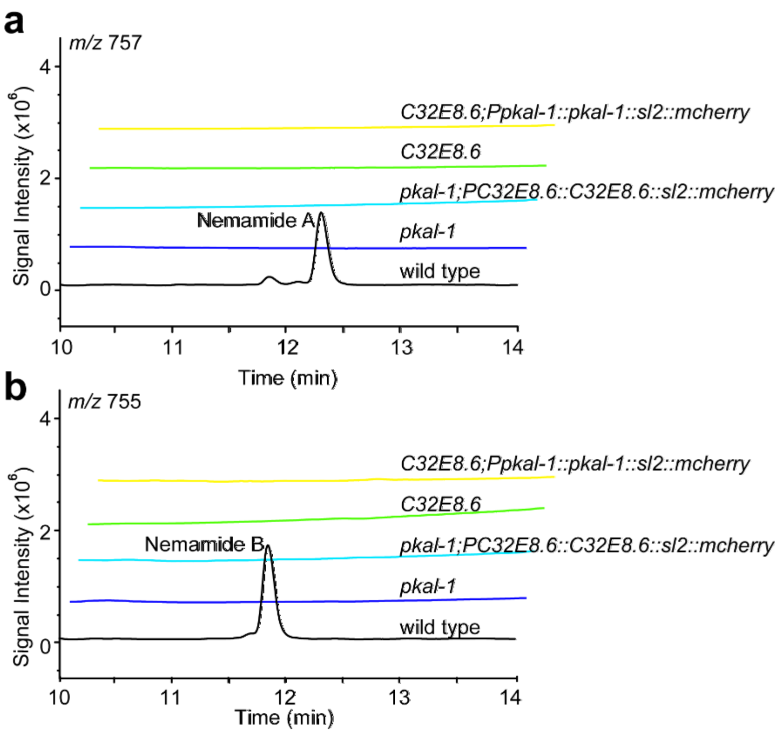


**Supplementary Figure 10. Failure of T20F7.7 (*pkal-1*) and C32E8.6 to rescue each other.** Extracted ion chromatogram for nemamide A (**a**) and nemamide B (**b**) in wild-type, the *pkal-1* mutant, the *pkal-1* mutant in which C32E8.6 was expressed under the control of its own promoter, the C32E8.6 mutant, and the C32E8.6 mutant in which *pkal-1* was expressed under the control of its own promoter. Note that the retention times of the nemamides in the Supplementary Information are different than in the main text due to the fact that the samples were analyzed on different columns.

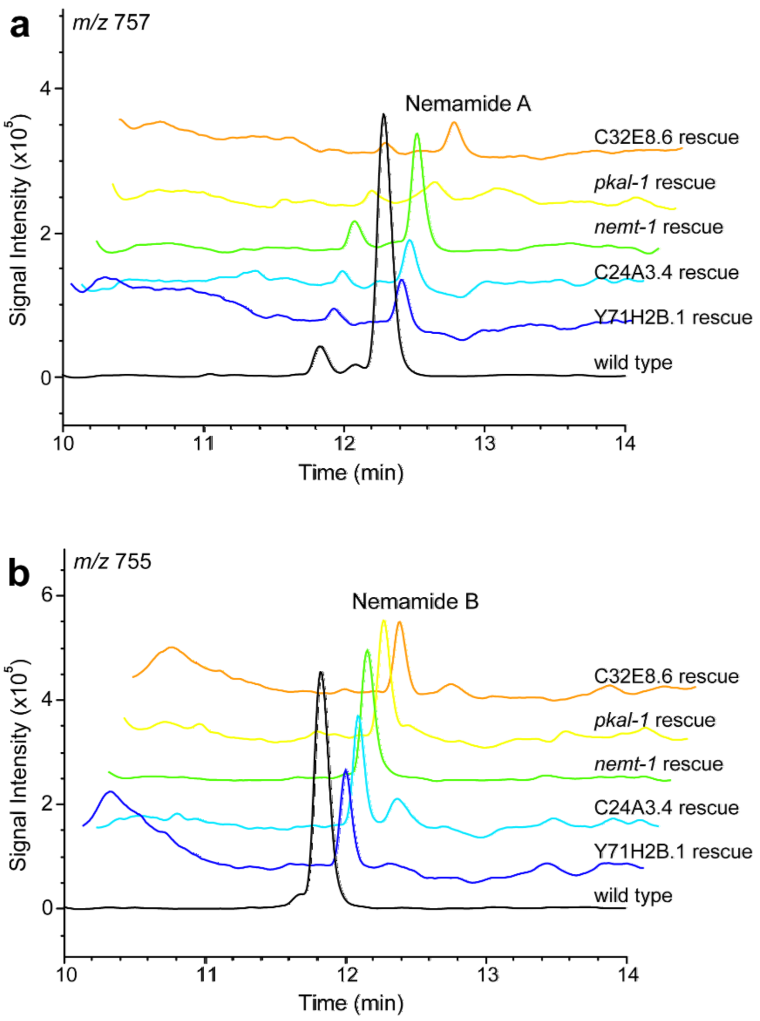


**Supplementary Figure 11. Nemamide production in wild type and mutant rescue strains**. Extracted ion chromatograms for nemamide A (**a**) and nemamide B (**b**). Mutants were rescued by complementing with *sl2::mCherry* plasmids under control of gene promoters::genes. Note that the retention times of the nemamides in the Supplementary Information are different than in the main text due to the fact that the samples were analyzed on different columns.

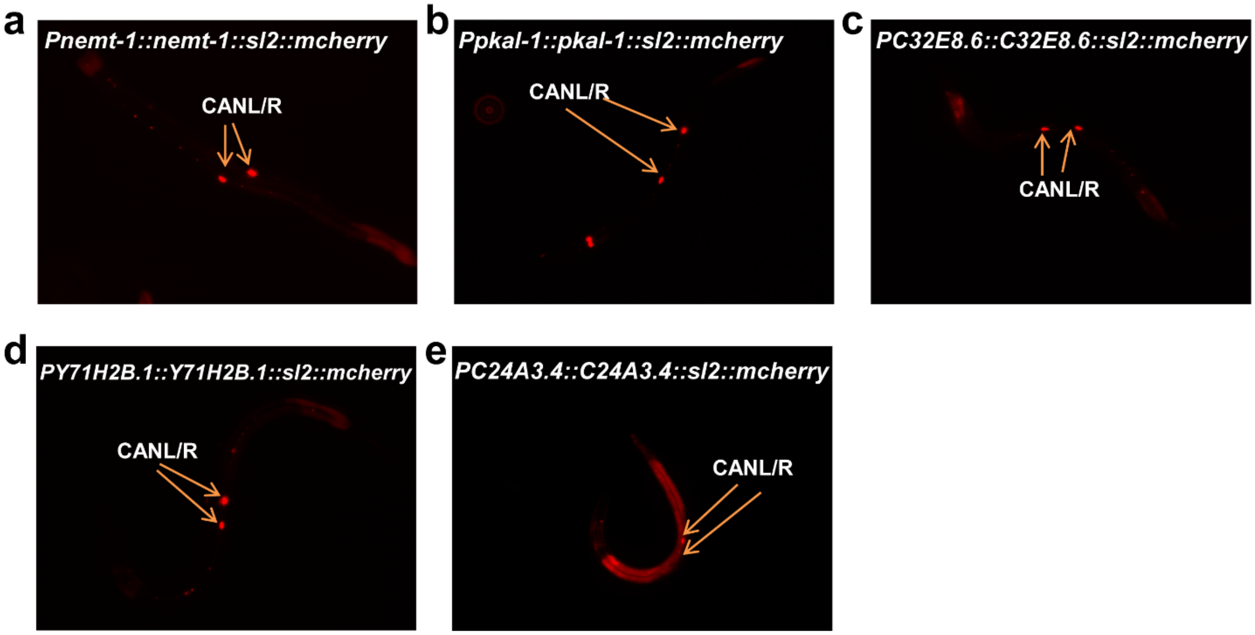


**Supplementary Figure 12. Expression of nemamide biosynthetic genes in the CANs.**

**a**

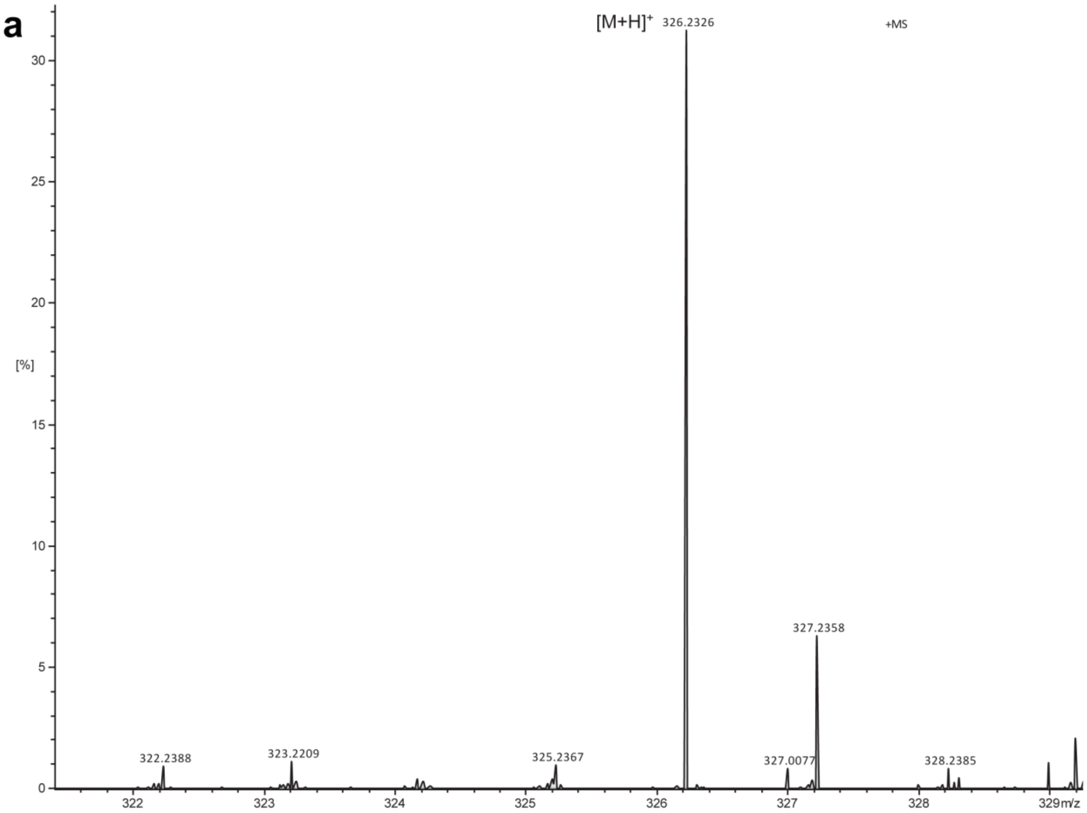
[M+H]+
326.2326
+MS
[%]
30
25
20
15
10
5
0
322.2388
323.2209
325.2367
327.2358
327.0077
328.2385
322
323
324
325
326
327
328
329m/z

**b**

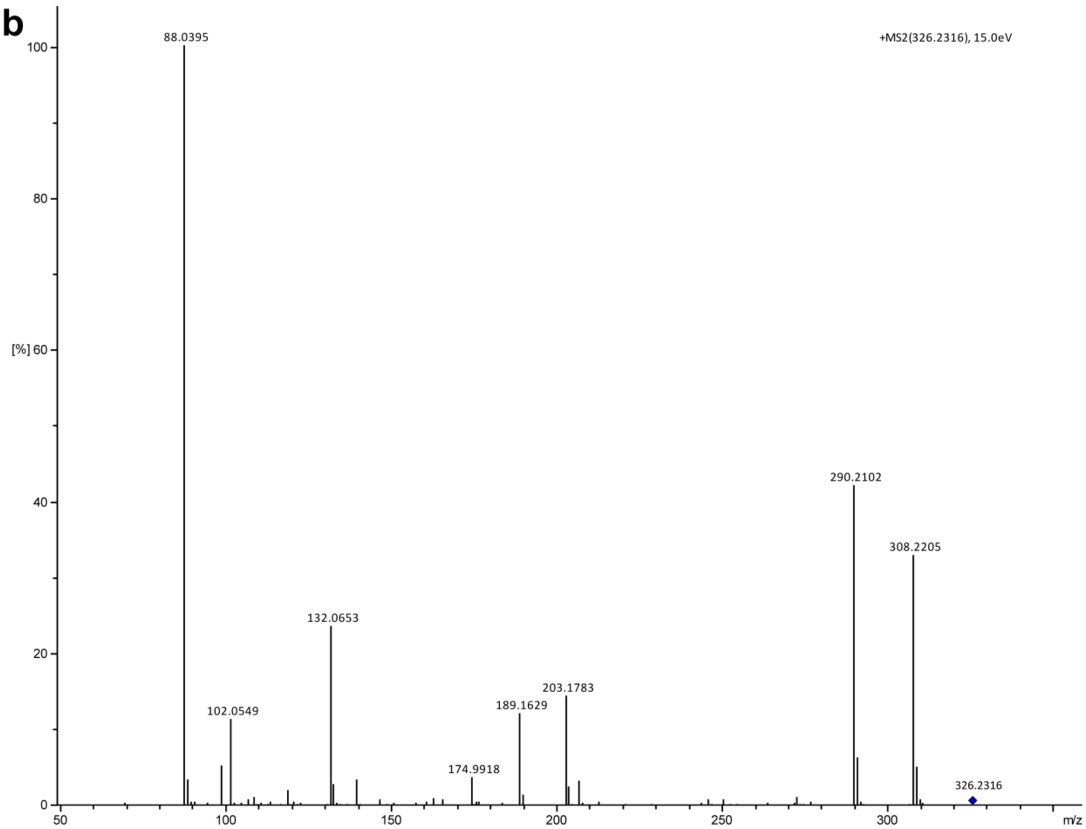
88.0395
+MS2(326.2316), 15.0eV
100
80
[%] 60
40
20
0
290.2102
308.2205
132.0653
102.0549
189.1629
203.1783
174.9918
326.2316
50
100
150
200
250
300
m/z

**c**

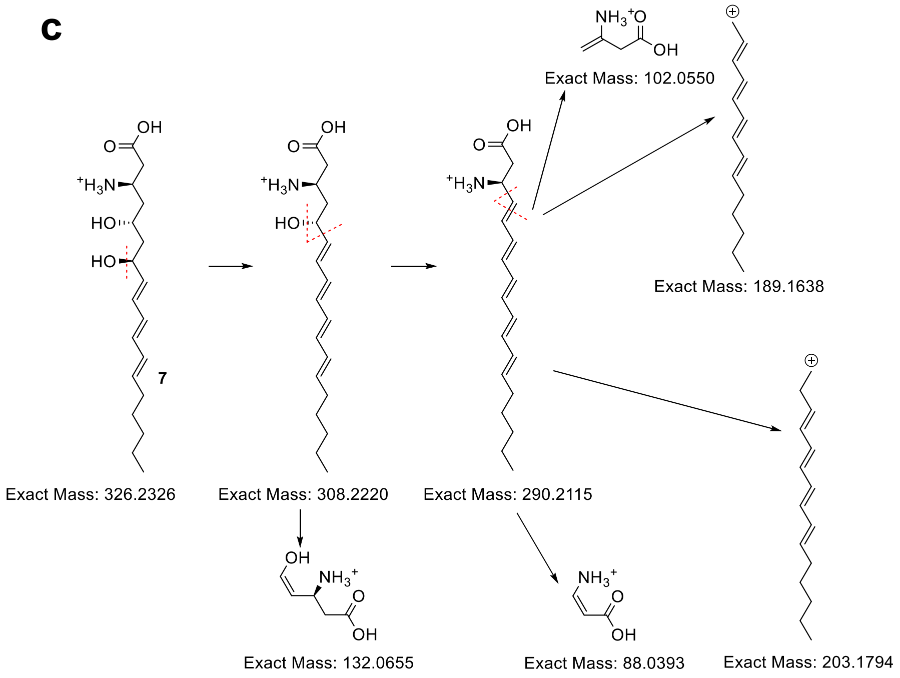


**Supplementary Figure 13. Mass spectrometry analysis of intermediate 7**. (**a**,**b**) HR-LC-MS/MS of intermediate **7**. (**c**) Analysis of the fragmentation patterns.

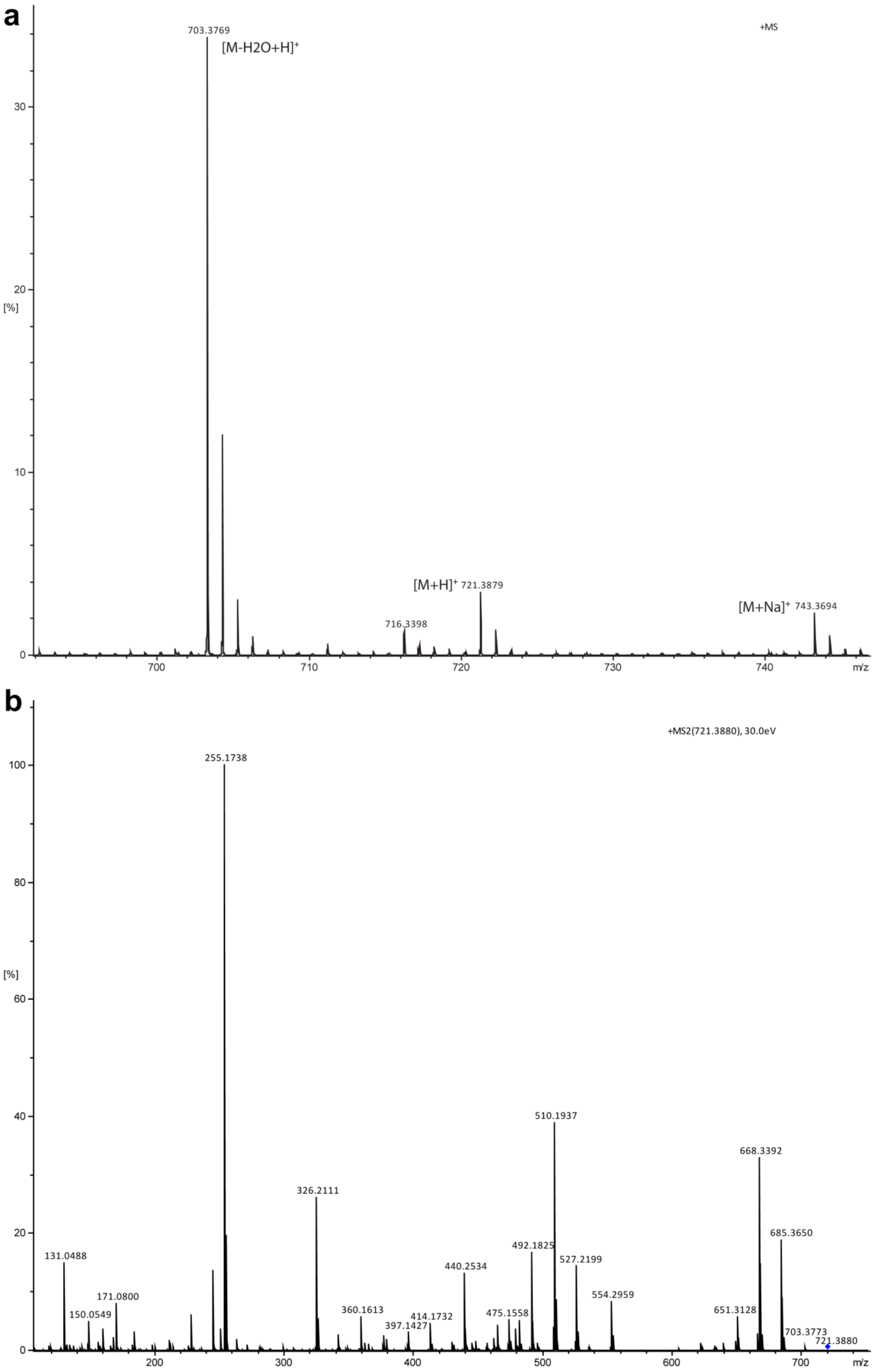
a
+MS
703.3769
[M-H2O+H]+
716.3398
[M+H]+ 721.3879
[M+Na]+ 743.3694
[%]
m/z
b
+MS2(721.3880), 30.0eV
255.1738
326.2111
131.0488
150.0549
171.0800
360.1613
397.1427
414.1732
440.2534
475.1558
492.1825
510.1937
527.2199
554.2959
651.3128
668.3392
685.3650
703.3773
721.3880
[%]
m/z

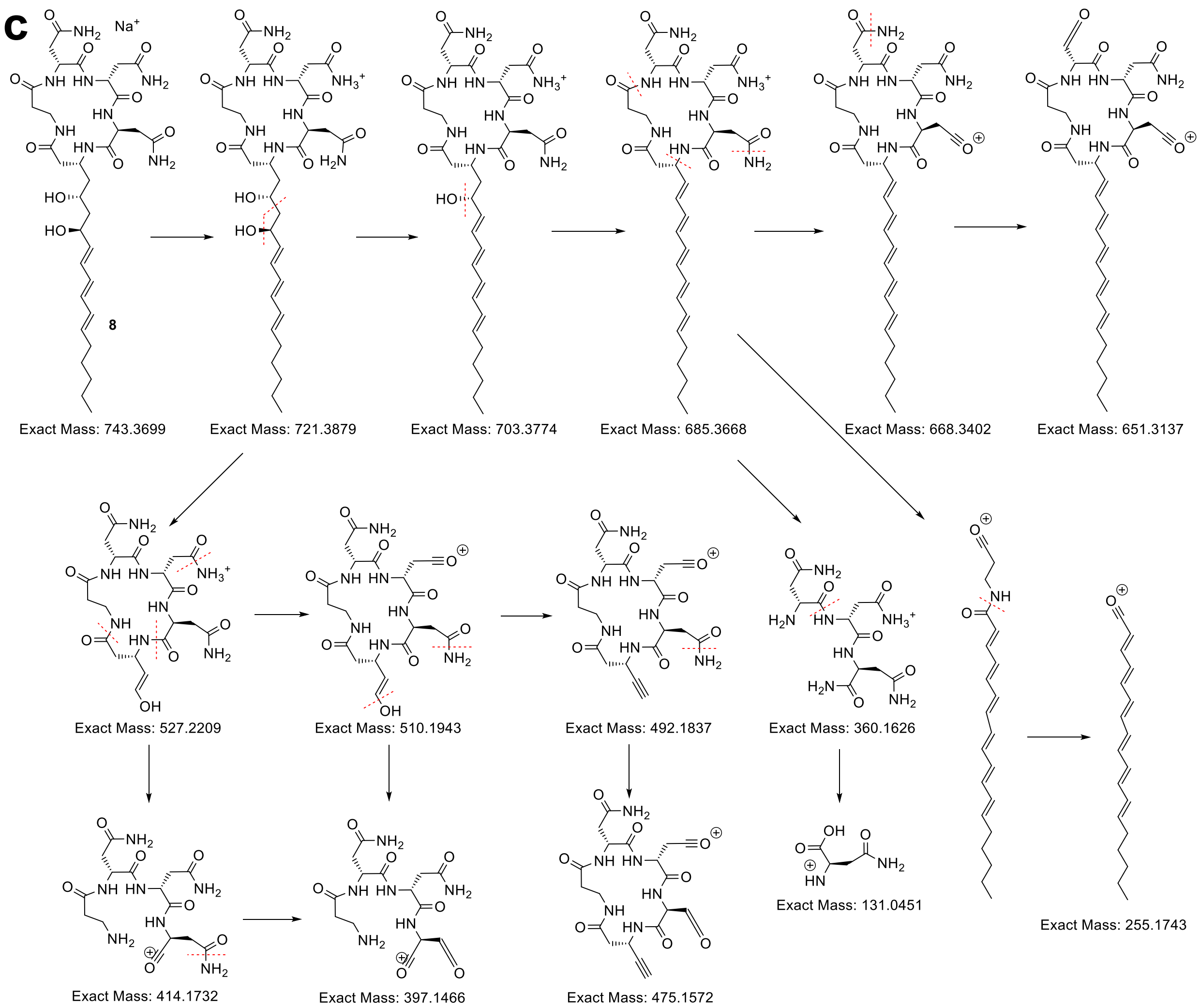


**Supplementary Figure 14. Mass spectrometry analysis of desmethyl-nemamide, 8**. **(a,b)** HR-LC-MS/MS of desmethyl-nemamide, **8**. **(c)** Analysis of the fragmentation patterns.

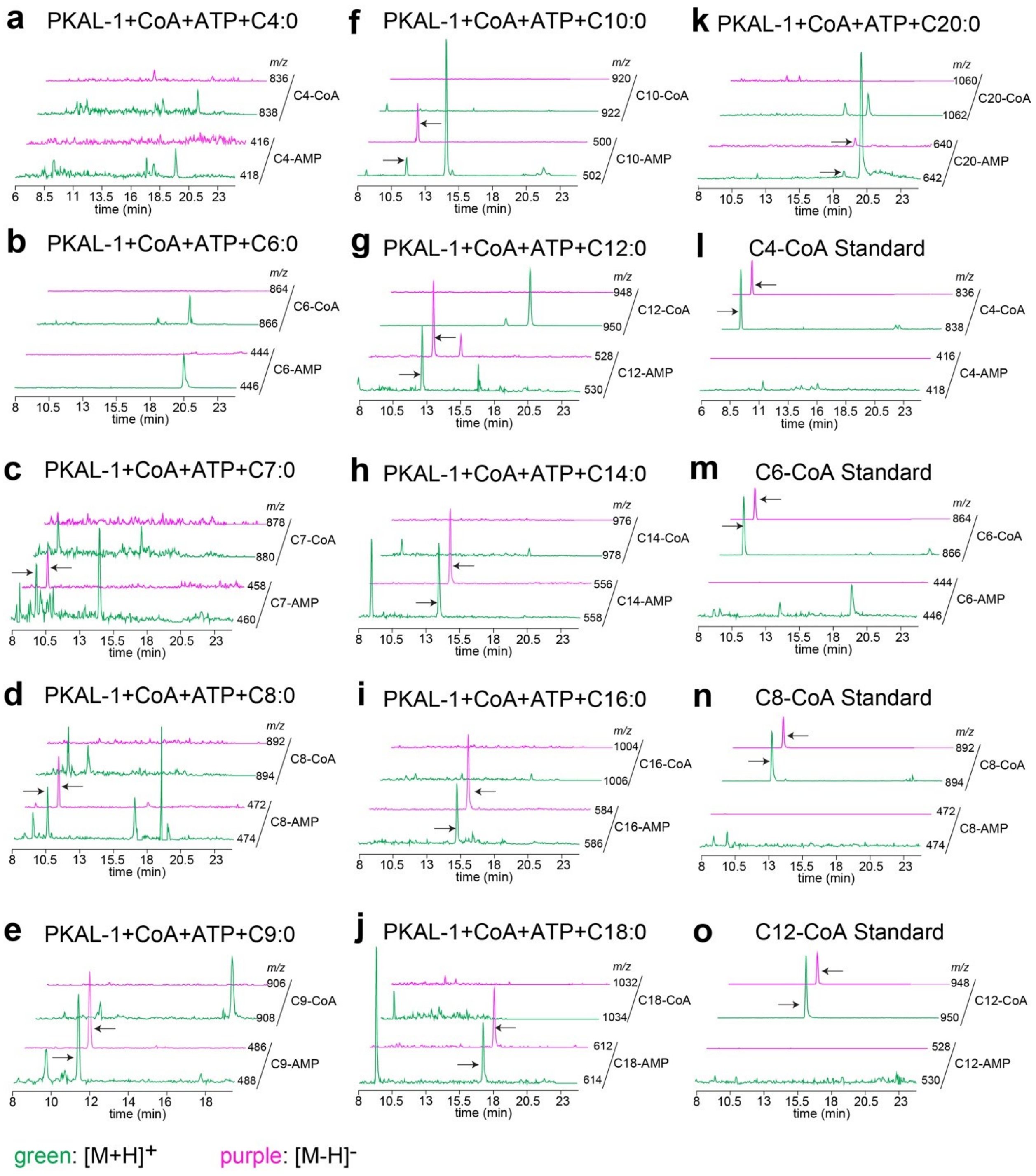


**Supplementary Figure 15. PKAL-1 is not an FACL.** Reaction of PKAL-1 with fatty acids of various lengths, ATP, and CoA did not result in the corresponding fatty acyl-CoAs. Several fatty acyl-CoAs were used as standards.

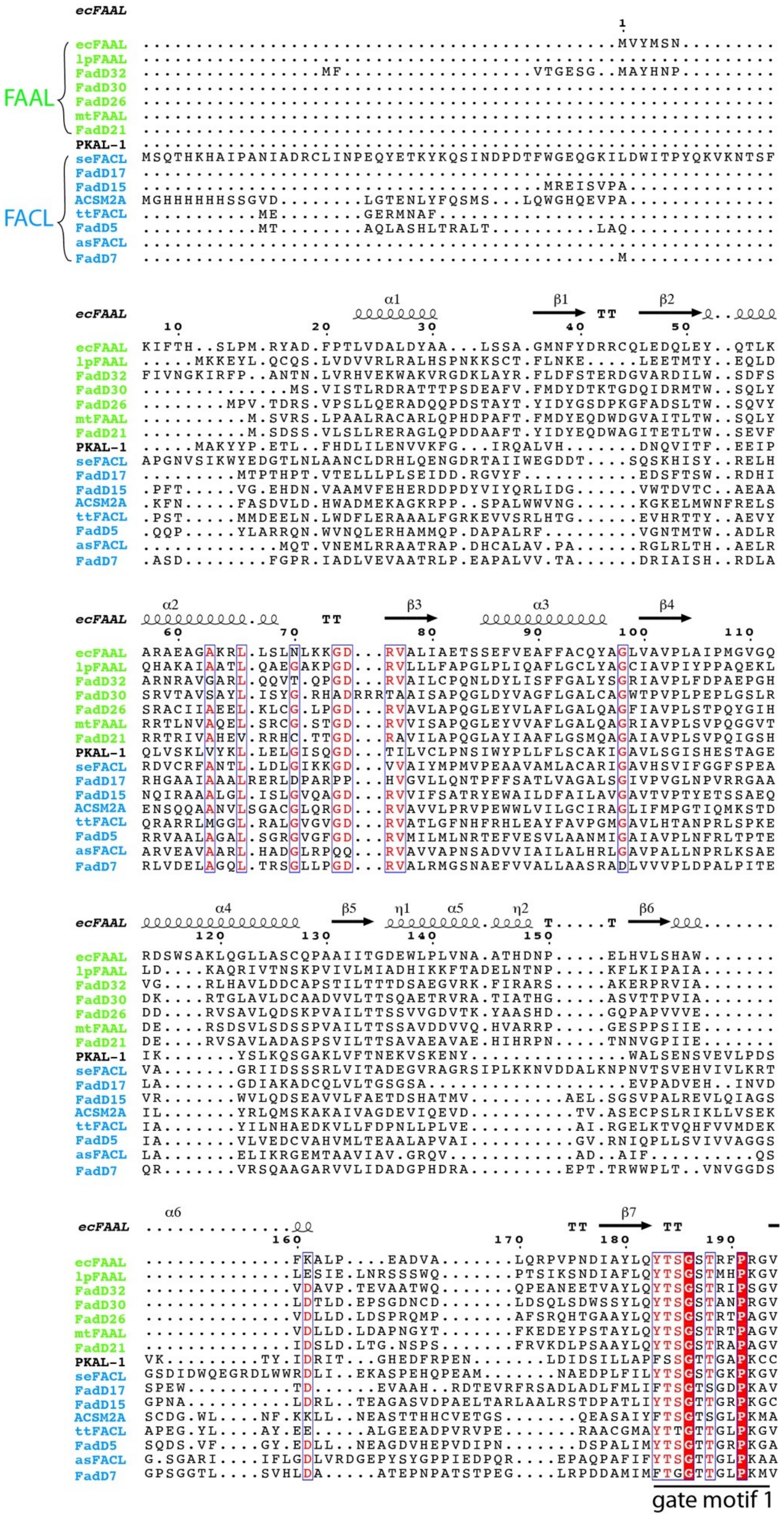
ecFAAL
FAAL
FACL
ecFAAL
lpFAAL
FadD32
FadD30
FadD26
mtFAAL
FadD21
PKAL-1
seFACL
FadD17
FadD15
ACSM2A
ttFACL
FadD5
asFACL
FadD7
gate motif 1

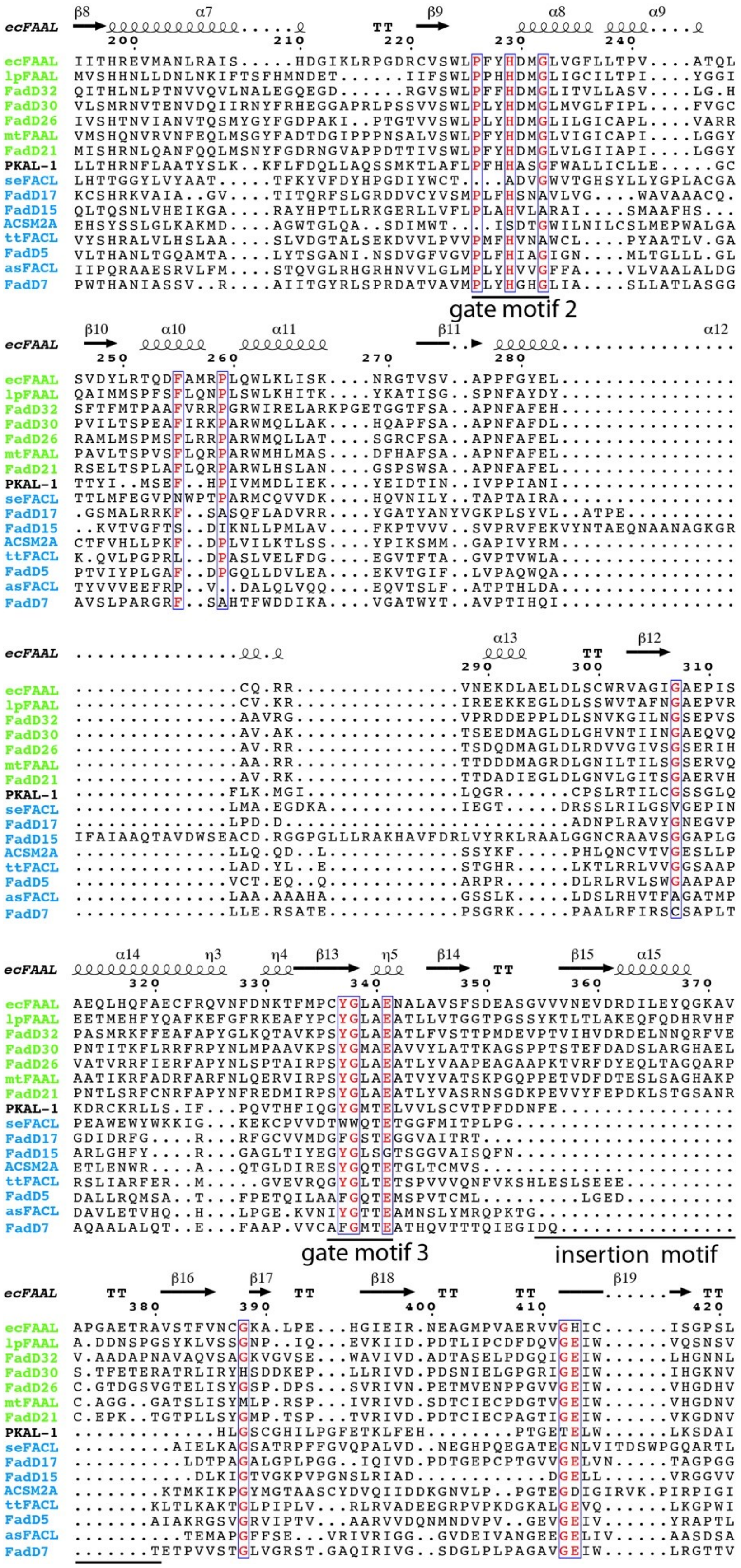
β8 α7 β9 α8 α9
ecFAAL TT 200 210 220 230 240
ecFAAL IITHREVMANLRAIS.....HDGIKLRPGDRCVSWLPFYHDMGLVGFLLTPV....ATQL
lpFAAL MVSHHNLLDNLNKIFTSFHMNDET......IIFSWLPPHHDMGLIGCILTPI....YGGI
FadD32 QITHLNLPTNVVQVLNALEGQEGD......RGVSWLPFFHDMGLITVVLLASV....LG.H
FadD30 VLSMRNVTENVDQIIRNYFRHEGGAPRLPSSVVSWLPLYHDMGLMVGLFIPL....FVGC
FadD26 IVSHTNVIANVTQSMYGYFGDPAKI..PTGTVVSWLPLYHDMGLILGICAPL....VARR
mtFAAL VMSHQNVRVNFEQLMSGYFADTDGIPPPNSALVSWLPFYHDMGLVIGICAPI....LGGY
FadD21 MISHRNLQANFQQLMSNYFGDRNGVAPPDTTIVSWLPFYHDMGLVLGIIAPI....LGGY
PKAL-1 LLTHRNFLAATYSLK..KFLFDQLLAQSSMKTLAFLPFHHASGFWALLICLLE.....GC
seFACL LHTTGGYLVYAAT.....TFKYVFDYHPGDIYWCT...ADVGWVTGHSYLLYGPLACGA
FadD17 KCSHRKVAIA..GV....TITQRFSLGRDDVCYVSMPLFHSNAVLVG....WAVAAACQ.
FadD15 QLTQSNLVHEIKGA....RAYHPTLLRKGERLLVFLPLAHVLARAI....SMAAFHS...
ACSM2A EHSYSSLGLKAKMD....AGWTGLQA..SDIMWT...ISDTGWILNILCSLMEPWALGA
ttFACL VYSHRALVLHSLAA....SLVDGTALSEKDVVLPVVPMFHVNAWCL....PYAATLV.GA
FadD5 VLTHANLTGQAMTA....LYTSGANI.NSDVGFVGVPLFHIAGIGN....MLTGLLL.GL
asFACL IIPQRAAESRVLFM....STQVGLRHGRHNVVLGLMPLYHVVGFFA....VLVAALALDG
FadD7 PWTHANIASSV..R....AIITGYRLSPRDATVAVMPLYHGHGLIA....SLLATLASGG
gate motif 2
β10 α10 α11 β11 α12
ecFAAL 250 260 270 280
ecFAAL SVDYLRTQDFAMRPLQWLKLISK....NRGTVSV..APPFGYEL...............
lpFAAL QAIMMSPFSFLQNPLSWLKHITK....YKATISG..SPNFAYDY...............
FadD32 SFTFMTPAAFVRRPGRWIRELARKPGETGGTFSA..APNFAFEH...............
FadD30 PVILTSPEAFIRKPARWMQLLAK....HQAPFSA..APNFAFDL...............
FadD26 RAMLMSPMSFLRRPARWMQLLAT....SGRCFSA..APNFAFEL...............
mtFAAL PAVLTSPVSFLQRPARWMHLMAS....DFHAFSA..APNFAFEL...............
FadD21 RSELTSPLAFLQRPARWLHSLAN....GSPSWSA..APNFAFEL...............
PKAL-1 TTYI..MSEF..HPIVMMDLIEK....YEIDTIN..IVPPIANI...............
seFACL TTLMFEGVPNWPTPARMCQVVDK....HQVNILY..TAPTAIRA...............
FadD17 .GSMALRRKF..SASQFLADVRR....YGATYANYVGKPLSYVL..ATPE.........
FadD15 ..KVTVGFTS..DIKNLLPMLAV....FKPTVVV..SVPRVFEKVYNTAEQNAANAGKGR
ACSM2A CTFVHLLPKF..DPLVILKTLSS....YPIKSMM..GAPIVYRM...............
ttFACL K.QVLPGPRL..DPASLVELFDG....EGVTFTA..GVPTVWLA...............
FadD5 PTVIYPLGAF..DPGQLLDVLEA....EKVTGIF..LVPAQWQA...............
asFACL TYVVVEEFRP..V..DALQLVQQ....EQVTSLF..ATPTHLDA...............
FadD7 AVSLPARGRF..SAHTFWDDIKA....VGATWYT..AVPTIHQI...............
α13 β12
ecFAAL TT 290 300 310
ecFAAL ...............CQ.RR..............VNEKDLAELDLSCWRVAGIGAEPIS
lpFAAL ...............CV.KR..............IREEKKEGLDLSSWVTAFNGAEPVR
FadD32 ..............AAVRG..............VPRDDEPPLDLSNVKGILNGSEPVS
FadD30 ..............AV.AK..............TSEEDMAGLDLGHVNTIINGAEQVQ
FadD26 ..............AV.RR..............TSDQDMAGLDLRDVVGIVSGSERIH
mtFAAL ..............AA.RR..............TTDDDMAGRDLGNILTILSGSERVQ
FadD21 ..............AV.RK..............TTDADIEGLDLGNVLGITSGAERVH
PKAL-1 ..............FLK.MGI.............LQGR......CPSLRTILCGSSGLQ
seFACL ..............LMA.EGDKA...........IEGT.....DRSSLRILGSVGEPIN
FadD17 ..............LPD.D.......................ADNPLRAVYGNEGVP
FadD15 IFAIAAQTAVDWSEACD.RGGPGLLLRAKHAVFDRLVYRKLRAALGGNCRAAVSGGAPLG
ACSM2A ..............LLQ.QD..L...........SSYKF.....PHLQNCVTVGESLLP
ttFACL ..............LAD.YL..E...........STGHR.....LKTLRRLVVGGSAAP
FadD5 ..............VCT.EQ..Q...........ARPR......DLRLRVLSWGAAPAP
asFACL ..............LAA.AAAHA...........GSSLK.....LDSLRHVTFAGATMP
FadD7 ..............LLE.RSATE...........PSGRK.....PAALRFIRSCSAPLT
α14 η3 η4 β13 η5 β14 β15 α15
ecFAAL TT 320 330 340 350 360 370
ecFAAL AEQLHQFAECFRQVNFDNKTFMPCYGLAENALAVSFSDEASGVVVNEVDRDILEYQGKAV
lpFAAL EETMEHFYQAFKEFGFRKEAFYPCYGLAEATLLVTGGTPGSSYKTLTLAKEQFQDHRVHF
FadD32 PASMRKFFEAFAPYGLKQTAVKPSYGLAEATLFVSTTPMDEVPTVIHVDRDELNNQRFVE
FadD30 PNTITKFLRRFRPYNLMPAAVKPSYGMAEAVVYLATTKAGSPPTSTEFDADSLARGHAEL
FadD26 VATVRRFIERFAPYNLSPTAIRPSYGLAEATLYVAAPEAGAAPKTVRFDYEQLTAGQARP
mtFAAL AATIKRFADRFARFNLQERVIRPSYGLAEATVYVATSKPGQPPETVDFDTESLSAGHAKP
FadD21 PNTLSRFCNRFAPYNFREDMIRPSYGLAEATLYVASRNSGDKPEVVYFEPDKLSTGSANR
PKAL-1 KDRCKRLLS.IF...PQVTHFIQGYGMTELVVLSCVTPFDDNFE................
seFACL PEAWEWYWKKIG...KEKCPVVDTWWQTETGGFMITPLPG....................
FadD17 GDIDRFG....R...RFGCVVMDGFGSTEGGVAITRT.......................
FadD15 ARLGHFY....R...GAGLTIYEGYGLSGTSGGVAISQFN....................
ACSM2A ETLENWR....A...QTGLDIRESYGQTETGLTCMVS.......................
ttFACL RSLIARFER..M.....GVEVRQGYGLTETSPVVVQNFVKSHLESLSEEE..........
FadD5 DALLRQMSA..T...FPETQILAAFGQTEMSPVTCML.........LGED..........
asFACL DAVLETVHQ..H...LPGE.KVNIYGTTEAMNSLYMRQPKTG..................
FadD7 AQAALALQT..E...FAAP.VVCAFGMTEATHQVTTTQIEGIDQ................
gate motif 3
insertion motif
β16 β17 β18 β19
ecFAAL TT TT TT TT TT 380 390 400 410 420
ecFAAL APGAETRAVSTFVNCGKA.LPE...HGIEIR.NEAGMPVAERVVGHIC......ISGPSL
lpFAAL A.DDNSPGSYKLVSSGNP..IQ...EVKIID.PDTLIPCDFDQVGEIW......VQSNSV
FadD32 V.AADAPNAVAQVSAGKVGVSE...WAVIVD.ADTASELPDGQIGEIW......LHGNNL
FadD30 S.TFETERATRLIRYHSDDKEP...LLRIVD.PDSNIELGPGRIGEIW......IHGKNV
FadD26 C.GTDGSVGTELISYGSP.DPS...SVRIVN.PETMVENPPGVVGEIW......VHGDHV
mtFAAL C.AGG..GATSLISYMLP.RSP...IVRIVD.SDTCIECPDGTVGEIW......VHGDNV
FadD21 C.EPK..TGTPLLSYGMP.TSP...TVRIVD.PDTCIECPAGTIGEIW......VKGDNV
PKAL-1 .............HLGSCGHILPGFETKLFEH........PTGETELW......LKSDAI
seFACL .........AIELKAGSATRPFFGVQPALVD..NEGHPQEGATEGNLVITDSWPGQARTL
FadD17 ..........LDTPAGALGPLPGG..IQIVD.PDTGEPCPTGVVGELVN.....TAGPGG
FadD15 ...........DLKIGTVGKPVPGNSLRIAD...........DGELL......VRGGVV
ACSM2A ........KTMKIKPGYMGTAASCYDVQIIDDKGNVLP..PGTEGDIGIRVK.PIRPIGI
ttFACL .......KLTLKAKTGLPIPLV...RLRVADEEGRPVPKDGKALGEVQ......LKGPWI
FadD5 .......AIAKRGSVGRVIPTV...AARVVDQNMNDVPV..GEVGEIV......YRAPTL
asFACL ..........TEMAPGFFSE...VRIVRIGG..GVDEIVANGEEGELIV.....AASDSA
FadD7 .......TETPVVSTGLVGRST.GAQIRIVG..SDGLPLPAGAVGEIW......LRGTTV

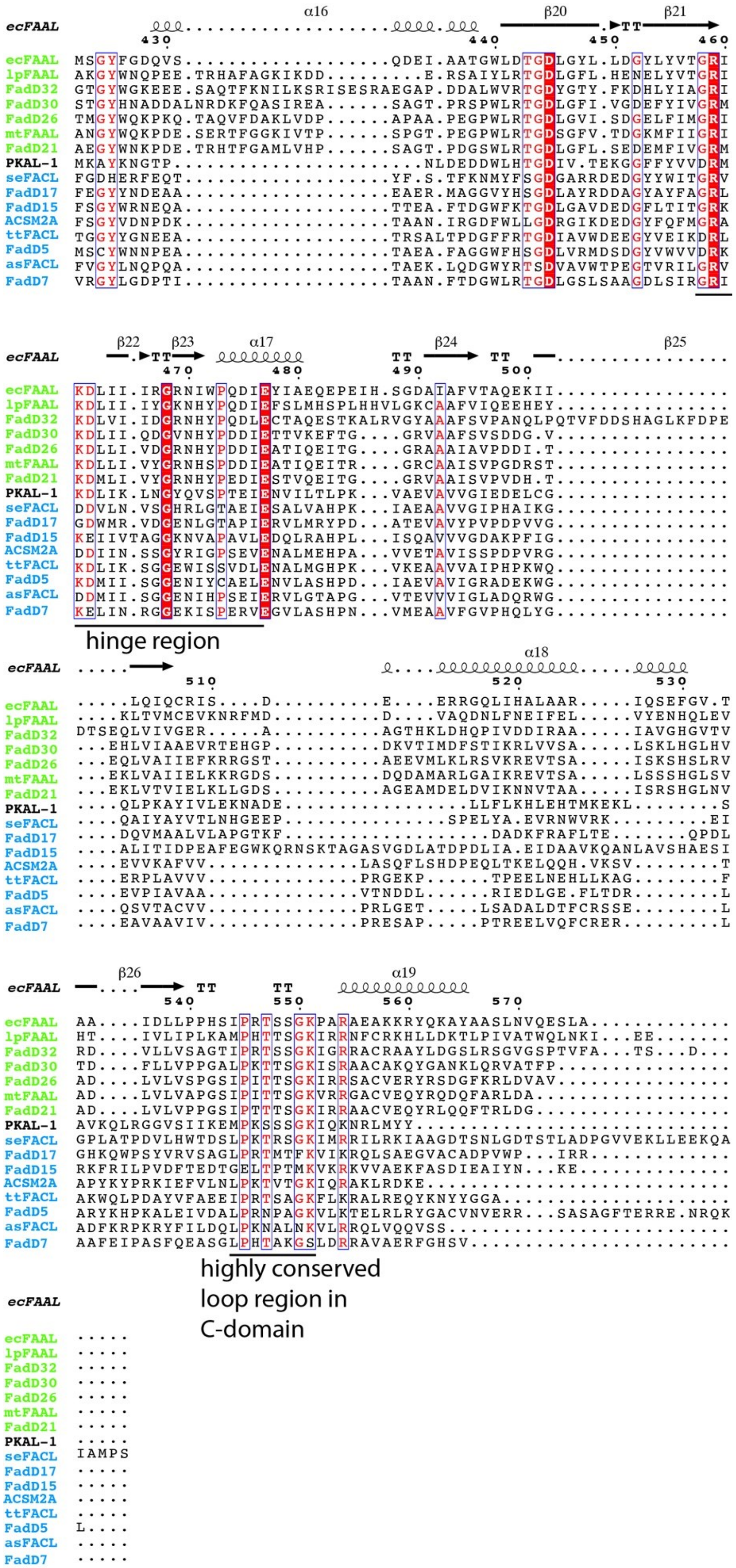
α16 β20 β21
ecFAAL
430 440 450 460
ecFAAL MSGYFGDQVS..................QDEI.AATGWLDTGDLGYL.LDGYLYVTGRI
lpFAAL AKGYWNQPEE.TRHAFAGKIKDD........E.RSAIYLRTGDLGFL.HENELYVTGRI
FadD32 GTGYWGKEEE.SAQTFKNILKSRISESRAEGAP.DDALWVRTGDYGTY.FKDHLYIAGRI
FadD30 STGYHNADDALNRDKFQASIREA.....SAGT.PRSPWLRTGDLGFI.VGDEFYIVGRM
FadD26 TMGYWQKPKQ.TAQVFDAKLVDP......APAA.PEGPWLRTGDLGVI.SDGELFIMGRI
mtFAAL ANGYWQKPDE.SERTFGGKIVTP.....SPGT.PEGPWLRTGDSGFV.TDGKMFIIGRI
FadD21 AEGYWNKPDE.TRHTFGAMLVHP.....SAGT.PDGSWLRTGDLGFL.SEDEMFIVGRM
PKAL-1 MKAYKNGTP......................NLDEDDWLHTGDIV.TEKGGFFYVVDRM
seFACL FGDHERFEQT..................YF.S.TFKNMYFSGDGARRDEDGYYWITGRV
FadD17 FEGYYNDEAA..................EAER.MAGGVYHSGDLAYRDDAGYAYFAGRL
FadD15 FSGYWRNEQA..................TTEA.FTDGWFKTGDLGAVDEDGFLTITGRK
ACSM2A FSGYVDNPDK..................TAAN.IRGDFWLLGDRGIKDEDGYFQFMGRA
ttFACL TGGYYGNEEA..................TRSALTPDGFFRTGDIAVWDEEGYVEIKDRL
FadD5 MSCYWNNPEA..................TAEA.FAGGWFHSGDLVRMDSDGYVWVVDRK
asFACL FVGYLNQPQA..................TAEK.LQDGWYRTSDVAVWTPEGTVRILGRV
FadD7 VRGYLGDPTI..................TAAN.FTDGWLRTGDLGSLSAAGDLSIRGRI
β22 β23 α17 β24 β25
ecFAAL
470 480 490 500
ecFAAL KDLII.IRGRNIWPQDIEYIAEQEPEIH.SGDAIAFVTAQEKII................
lpFAAL KDLII.IYGKNHYPQDIEFSLMHSPLHHVLGKCAAFVIQEEHEY................
FadD32 KDLVI.IDGRNHYPQDLECTAQESTKALRVGYAAAFSVPANQLPQTVFDDSHAGLKFDPE
FadD30 KDLII.QDGVNHYPDDIETTVKEFTG....GRVAAFSVSDDG.V................
FadD26 KDLLI.VDGRNHYPDDIEATIQEITG....GRAAAIAVPDDI.T................
mtFAAL KDLLI.VYGRNHSPDDIEATIQEITR....GRCAAISVPGDRST................
FadD21 KDMLI.VYGRNHYPEDIESTVQEITG....GRVAAISVPVDH.T................
PKAL-1 KDLIK.LNGYQVSPTEIENVILTLPK...VAEVAVVGIEDELCG................
seFACL DDVLN.VSGHRLGTAEIESALVAHPK...IAEAAVVGIPHAIKG................
FadD17 GDWMR.VDGENLGTAPIERVLMRYPD...ATEVAVYPVPDPVVG................
FadD15 KEIIVTAGGKNVAPAVLEDQLRAHPL...ISQAVVVGDAKPFIG................
ACSM2A DDIIN.SSGYRIGPSEVENALMEHPA...VVETAVISSPDPVRG................
ttFACL KDLIK.SGGEWISSVDLENALMGHPK...VKEAAVVAIPHPKWQ................
FadD5 KDMII.SGGENIYCAELENVLASHPD...IAEVAVIGRADEKWG................
asFACL DDMII.SGGENIHPSEIERVLGTAPG...VTEVVVIGLADQRWG................
FadD7 KELIN.RGGEKISPERVEGVLASHPN...VMEAAVFGVPHQLYG................
hinge region
α18
ecFAAL
510 520 530
ecFAAL .....LQIQCRIS....D..........E....ERRGQLIHALAAR.....IQSEFGV.T
lpFAAL ....KLTVMCEVKNRFMD..........D....VAQDNLFNEIFEL.....VYENHQLEV
FadD32 DTSEQLVIVGER.....A..........AGTHKLDHQPIVDDIRAA.....IAVGHGVTV
FadD30 ...EHLVIAAEVRTEHGP..........DKVTIMDFSTIKRLVVSA.....LSKLHGLHV
FadD26 ...EQLVAIIEFKRRGST..........AEEVMLKLRSVKREVTSA.....ISKSHSLRV
mtFAAL ...EKLVAIIELKKRGDS..........DQDAMARLGAIKREVTSA.....LSSSHGLSV
FadD21 ...EKLVTVIELKLLGDS..........AGEAMDELDVIKNNVTAA.....ISRSHGLNV
PKAL-1 ....QLPKAYIVLEKNADE................LLFLKHLEHTMKEKL........S
seFACL ....QAIYAYVTLNHGEEP...............SPELYA.EVRNWVRK.........EI
FadD17 ....DQVMAALVLAPGTKF..................DADKFRAFLTE.......QPDL
FadD15 ....ALITIDPEAFEGWKQRNSKTAGASVGDLATDPDLIA.EIDAAVKQANLAVSHAESI
ACSM2A ....EVVKAFVV..............LASQFLSHDPEQLTKELQQH.VKSV........T
ttFACL ....ERPLAVVV..............PRGEKP......TPEELNEHLLKAG........F
FadD5 ....EVPIAVAA..............VTNDDL......RIEDLGE.FLTDR........L
asFACL ....QSVTACVV..............PRLGET.....LSADALDTFCRSSE........L
FadD7 ....EAVAAVIV..............PRESAP.....PTREELVQFCRER.........L
β26 α19
ecFAAL
540 550 560 570
ecFAAL AA....IDLLPPHSIPRTSSGKPARAEAKKRYQKAYAASLNVQESLA.............
lpFAAL HT....IVLIPLKAMPHTTSGKIRRNFCRKHLLDKTLPIVATWQLNKI...EE.......
FadD32 RD....VLLVSAGTIPRTSSGKIGRRACRAAYLDGSLRSGVGSPTVFA...TS...D...
FadD30 TD....FLLVPPGALPKTTSGKISRAACAKQYGANKLQRVATFP................
FadD26 AD....LVLVSPGSIPITTSGKIRRSACVERYRSDGFKRLDVAV................
mtFAAL AD....LVLVAPGSIPITTSGKVRRGACVEQYRQDQFARLDA..................
FadD21 AD....LVLVPPGSIPTTTSGKIRRAACVEQYRLQQFTRLDG..................
PKAL-1 AVKQLRGGVSIIKEMPKSSSGKIQKNRLMYY.............................
seFACL GPLATPDVLHWTDSLPKTRSGKIMRRILRKIAAGDTSNLGDTSTLADPGVVEKLLEEKQA
FadD17 GHKQWPSYVRVSAGLPRTMTFKVIKRQLSAEGVACADPVWP...IRR.............
FadD15 RKFRILPVDFTEDTGELTPTMKVKRKVVAEKFASDIEAIYN...KE..............
ACSM2A APYKYPRKIEFVLNLPKTVTGKIQRAKLRDKE............................
ttFACL AKWQLPDAYVFAEEIPRTSAGKFLKRALREQYKNYYGGA.....................
FadD5 ARYKHPKALEIVDALPRNPAGKVLKTELRLRYGACVNVERR...SASAGFTERRE.NRQK
asFACL ADFKRPKRYFILDQLPKNALNKVLRRQLVQQVSS..........................
FadD7 AAFEIPASFQEASGLPHTAKGSLDRRAVAERFGHSV........................
highly conserved loop region in C-domain
ecFAAL
ecFAAL .....
lpFAAL .....
FadD32 .....
FadD30 .....
FadD26 .....
mtFAAL .....
FadD21 .....
PKAL-1 .....
seFACL IAMPS
FadD17 .....
FadD15 .....
ACSM2A .....
ttFACL .....
FadD5 L....
asFACL .....
FadD7 .....

**Supplementary Figure 16. Sequence alignment of PKAL-1 with FAAL enzymes.** The FAAL enzymes (in green) are ecFAAL from *Escherichia coli*, lpFAAL from *Legionella pneumophila*, mtFAAL from *Mycobacterium tuberculosis*, and FadD21, FadD26, FadD30, and FadD32 from *M. tuberculosis*, and FACL enzymes (in blue), including seFACL from *Salmonella enterica*, ttFACL from *Thermus thermophilus*, asFACL from *Alcaligenes sp.*, ACSM2A from human, and FadD5, FadD7, FadD15, and FadD17 from *M. tuberculosis*. The conserved motifs, including the insertion motif that is present in the FAAL enzymes, but missing in PKAL-1 and FACL enzymes, are indicated.

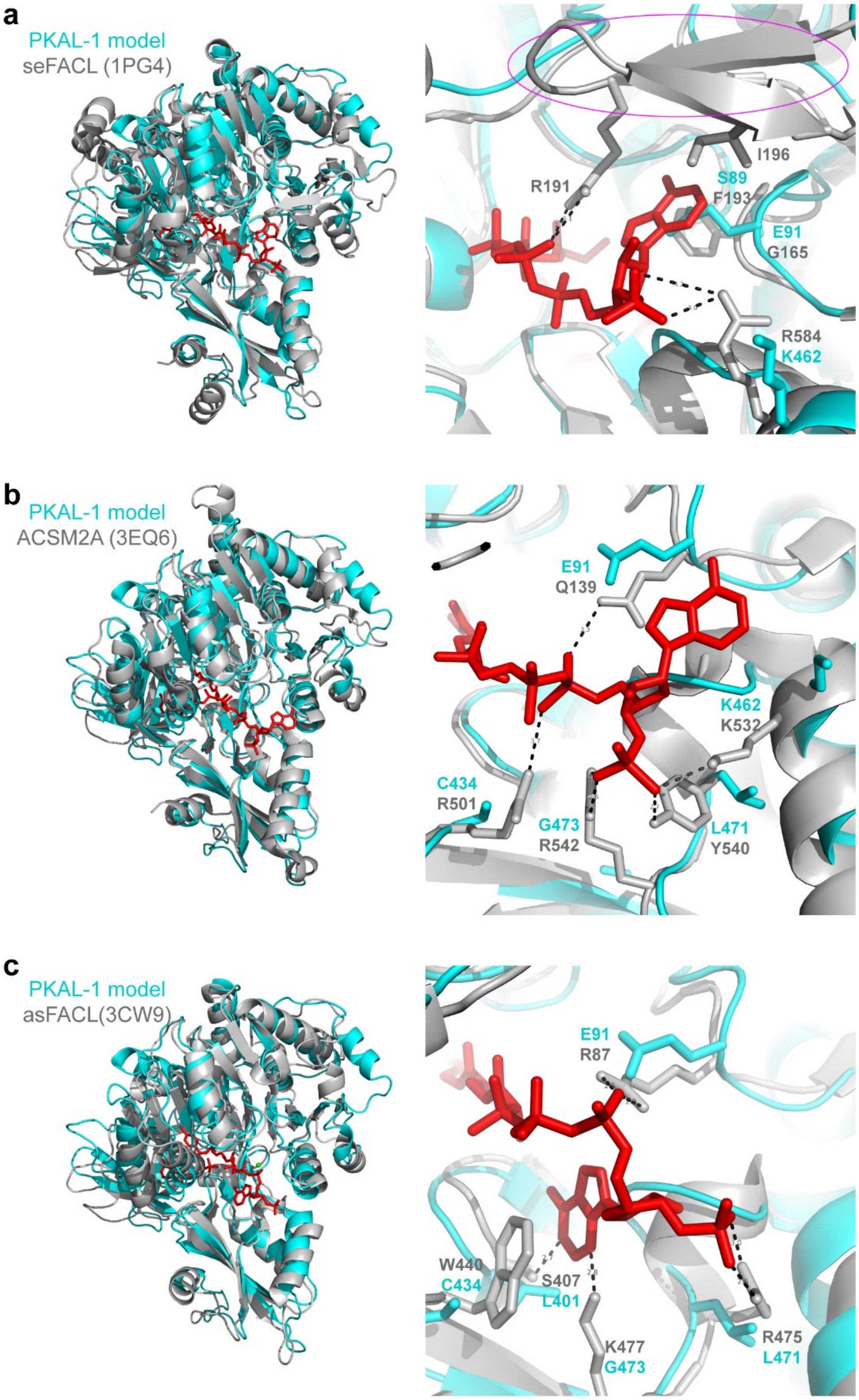


**Supplementary Figure 17. Comparison of the modeled PKAL-1 structure to the structures of FACL enzymes bound to CoA substrates.** PKAL-1 was modeled using Phyre2, and six templates 5GXD, 6EQO, 6PLJ, 5ES8, 6MFZ, and 5U89 were selected for modeling, enabling 100% of residues to be modeled at >90% confidence. The PKAL-1 structural model was then overlaid with the structures of three FACL enzymes in the thioester-forming conformation: (**a**) 1PG4, the structure of acetyl-CoA synthetase from *Salmonella enterica* (seFACL), bound to adenosine-5'-propylphosphate and CoA (in red), (**b**) 3EQ6, the structure of the human medium-chain acyl-CoA synthetase (ACSM2A) bound to AMP and butyryl-CoA (in red), and (**c**) 3CW9, the structure of 4-chlorobenzoate:CoA ligase from *Alcaligenes sp.* (asFACL) bound to 4-chlorophenyacyl-CoA (in red). In (**a**), the seFACL structure has a beta hairpin (circled in pink), which contains R191 that binds CoA and which is missing in the PKAL-1 model. A hydrophobic pocket for the adenine ring of CoA that is formed by I196 in the beta hairpin and F193 and G165 is also missing in the PKAL-1 model. In (**b**), the ACSM2A structure has several residues, including R501, R542, Y540, and Q139, which are important for binding of the CoA substrate and which are replaced with C434, G473, L471, and E91, respectively, in the PKAL-1 model. In (**c**), the asFACL structure has several residues, including W440, S407, K477, R475, and R87, which are important for binding of the CoA substrate and which are replaced with C434, L401, G473, L471, and E91, respectively, in the PKAL-1 model.

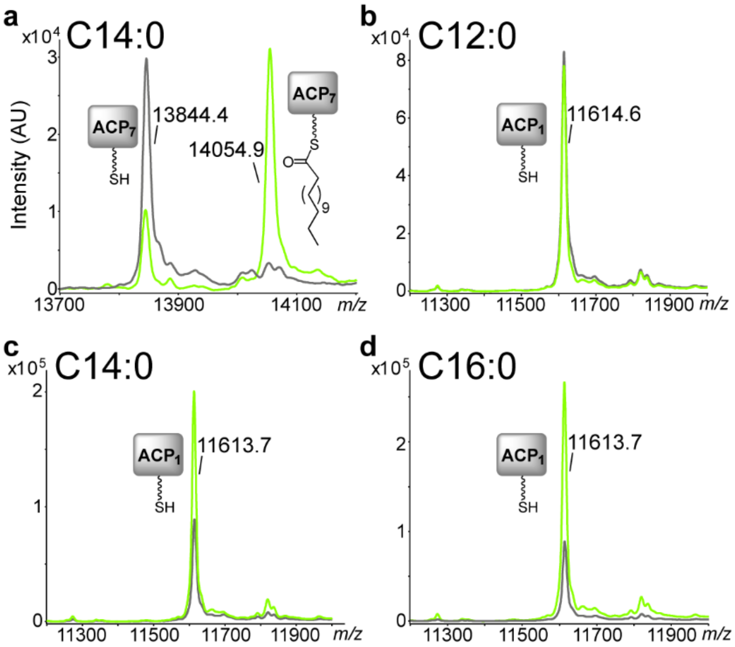


**Supplementary Figure 18. Loading of PKS-1_ACP$_1$ or NRPS-1_ACP$_7$ by PKAL-1.** To determine the carrier protein specificity of PKAL-1, PKAL-1 was incubated with (**a**) holo-ACP$_7$ (positive control), ATP, and C14:0 fatty acid, or with (**b**-**d**) holo-ACP$_1$ from PKS-1, ATP, and C12:0 fatty acid (**b**), C14:0 fatty acid (**c**), or C16:0 fatty acid (**d**). Samples were analyzed by MALDI to determine if the fatty acid substrates were loaded onto the respective carrier proteins.

**Supplementary Table 1. A domain selectivity codes for the PKS-1 $A_1$ domain from various nematode species.**

| Species | 235 | 236 | 239 | 278 | 299 | 301 | 322 | 330 | 331 | 517 |
|---|---|---|---|---|---|---|---|---|---|---|
| *C. elegans* | D | V | S | F | T | G | I | I | W | K |
| *C. angaria* | D | V | S | F | T | G | I | I | W | K |
| *C. japonica* | D | V | A | F | T | G | I | V | W | K |
| *C. brenneri* | D | V | S | F | T | G | I | I | W | K |
| *C. remanei* | D | V | S | F | T | G | I | V | W | K |
| *C. briggsae* | D | V | S | F | T | G | I | V | W | K |
| *C. tropicalis* | D | V | S | F | T | G | I | I | W | K |
| *A. suum* | D | V | M | Y | F | G | I | I | W | K |
| *T. canis* | D | V | M | F | F | G | I | I | W | K |
| *D. immitis* | D | V | M | F | Y | G | I | I | W | K |
| *O. volvulus* | D | V | V | F | Y | G | I | V | W | K |
| *L. loa* | D | V | V | F | Y | G | I | V | W | K |
| *B. malayi* | D | V | M | F | F | G | I | I | W | K |
| *P. pacificus* | D | V | F | F | I | G | I | I | W | K |
| *P. exspectatus* | D | V | F | F | I | G | I | I | W | K |
| *S. carpocapsae* | D | V | F | F | Y | G | I | I | W | K |
| *B. xylophilus* | D | V | F | F | I | G | I | I | W | K |
| *A. ceylanicum* | D | V | M | F | F | G | I | V | W | K |
| *A. duodenale* | D | V | M | F | L | G | I | I | W | K |
| *O. dentatum* | D | V | L | F | F | G | I | V | W | K |
| *N. americanus* | D | V | F | F | V | G | I | V | W | K |
| *H. bacteriophora* | D | V | V | F | F | G | I | V | W | K |
| *H. contortus* | D | V | F | F | F | G | I | V | W | K |

**Supplementary Table 2. Strains used in this study.**

| Strain | Genotype | Mutation | Background | Resource |
|---|---|---|---|---|
| N2 | wild type | | | CGC |
| RAB43 | *nrps-1(gk186409[$C_4$_S1934N]; gk186410[$C_4$_D1971N])* III | Located in NRPS-1_$C_4$ | VC20469 (4X outcrossing) | CGC |
| RAB45 | Y71H2B.1*(gk712674)* III | W136Opal | VC40597(2X outcrossing) | CGC |
| RAB51 | *nrps-1(reb8[$ACP_7$_S307V])* III | NRPS-1_$ACP_7$ | N2 | CRISPR-Cas9 |
| RAB52 | *pks-1(reb9[$C_1$_H6685A])* X | PKS-1_$C_1$ | N2 | CRISPR-Cas9 |
| RAB53 | *nrps-1(reb10[$C_3$_H1486A])* III | NRPS-1_$C_3$ | N2 | CRISPR-Cas9 |
| RAB54 | *pks-1(reb11[$TE_1$_S7593A])* X | PKS-1_$TE_1$ | N2 | CRISPR-Cas9 |
| RAB55 | *nrps-1(reb12[$TE_2$_S2803A])* III | NRPS-1_$TE_2$ | N2 | CRISPR-Cas9 |
| RAB56 | *pks-1(reb13[$TE_1$_S7593C]; reb14[$TE_1$_G7596A])* X | PKS-1_$TE_1$ | N2 | CRISPR-Cas9 |
| RAB57 | *nemt-1(reb15)* IV | 306 bp deletion with 27 bp insertion | N2 | CRISPR-Cas9 |
| RAB58 | *pkal-1(reb21)* X | 154 bp deletion with single base 'T' insertion | N2 | CRISPR-Cas9 |
| RAB59 | *pkal-1(reb28)* X | 552 bp deletion | N2 | CRISPR-Cas9 |
| RAB60 | *C32E8.6(reb23)* I | 991 bp deletion with 10 bp insertion | N2 | CRISPR-Cas9 |
| RAB61 | *C32E8.6(reb24)* I | 993 bp deletion | N2 | CRISPR-Cas9 |
| RAB62 | C24A3.4*(reb16)* X | 1361 bp deletion | N2 | CRISPR-Cas9 |
| RAB67 | *pkal-1(reb28)*; *pks-1(reb11[$TE_1$_S7593A])* | | | Outcrossing |
| RAB68 | *pkal-1(reb28)*; *nemt-1(reb15)* | | | Outcrossing |
| RAB69 | *nemt-1(reb15); pks-1(reb11[$TE_1$_ S7593A])* | | | Outcrossing |
| RAB72 | *rebEx15 (Pnemt-1::gfp, 50 ng/µL; CAN::mcherry, 50 ng/µL)* | | N2 | Transgenesis |
| RAB73 | *rebEx16 (Ppkal-1::gfp, 50 ng/µL; CAN::mcherry, 50 ng/µL)* | | N2 | Transgenesis |
| RAB74 | *rebEx17 (PC32E8.6::gfp, 50 ng/µL; CAN::mcherry, 50 ng/µL)* | | N2 | Transgenesis |
| RAB76 | *nemt-1(reb15); rebEx19(Pnemt-1::nemt-1::sl2::mcherry, 50 ng/µL)* | | | Transgenesis |
| RAB77 | *pkal-1(reb28); rebEx20(Ppkal-1::pkal-1::sl2::mcherry, 50 ng/µL)* | | | Transgenesis |
| RAB78 | *C32E8.6(reb24); rebEx21(PC32E8.6::C32E8.6::sl2::mcherry, 50 ng/µL)* | | | Transgenesis |

| | | | | |
|---|---|---|---|---|
| RAB79 | *C32E8.6(reb24); rebEx20(Ppkal-1::pkal-1::sl2::mcherry, 50 ng/µL)* | | | Transgenesis |
| RAB80 | *pkal-1(reb28); rebEx21(PC32E8.6::C32E8.6::sl2::mcherry, 50 ng/µL)* | | | Transgenesis |
| RAB81 | Y71H2B.1*(gk712674); rebEx22(P*Y71H2B.1*::*Y71H2B.1*::sl2::mcherry, 50 ng/µL)* | | | Transgenesis |
| RAB82 | C24A3.4(*reb16); rebEx23(PC24A3.4::C24A3.4::sl2::mcherry, 50 ng/µL)* | | | Transgenesis |
| RAB89 | *pks-1(reb22[$A_1$_G7106E])* X | PKS-1_$A_1$ | N2 | CRISPR-Cas9 |
| RAB103 | *pks-1(reb29[$PCP_2$_S7463A])*X | PKS-1_$PCP_2$ | N2 | CRISPR-Cas9 |

**Supplemental Table 3. Single worm PCR primers for mutant strains in this study.**

| Strain | Genotype | Primers |
|---|---|---|
| RAB43 | *nrps-1 (gk186409[$C_4$_S1934N]; gk186410[$C_4$_D1971N])* III | Forward_CTGAAGCCTTTATTCAGTGCCAAG<br>Reverse_CTTGCACTGCTAGAGCTAAGCTTC |
| RAB45 | Y71H2B.1*(gk712674)* III | Forward_GGAAAGCACGGAGATTTTGAAG<br>Reverse_AGTGATGGGAATGGTCTCTGTT |
| RAB51 | *nrps-1(reb8[$ACP_7$_S307V])* III | Forward_GAAGGAGCAGCAAACATCGAGAA<br>Reverse_ATCTGAGTGACCTGCTTTCAGAG |
| RAB52 | *pks-1(reb9[$C_1$_H6685A])* X | Forward_CATCTGTAAACCCTGCAGATATTGC<br>Reverse_CGGCATCGCAGAAAACTGATAATGC |
| RAB53 | *nrps-1(reb10[$C_3$_H1486A])* III | Forward_GAAGCTGGTGGAGTTGTCCAATGCT<br>Reverse_GAAACTGTATCCCAGTTCTCTGGAG |
| RAB54 | *pks-1(reb11[$TE_1$_S7593A])* X | Forward_GGTGATTAAATCTGGAGTAC<br>Reverse_TAGTCCAGAGAAGACGTACT |
| RAB55 | *nrps-1(reb12[$TE_2$_S2803A])* III | Forward_TCGAGACCAAACTCGGAATC<br>Reverse_TCTGAGAAAATGTTCACCGG |
| RAB56 | *pks-1(reb13[$TE_1$_S7593C]; reb14[$TE_1$_G7596A])* X | Forward_GAGGTGATTAAATCTGGAGTACGGC<br>Reverse_TCACTATCCGGTAGTCCAGAGAAG |
| RAB57 | *nemt-1(reb15)* IV | Forward_AGTGGCTTTGCCTTTCCTCCTT<br>Reverse_AGCCCTCAACTACTTCATCAGTG |
| RAB58 | *pkal-1(reb21)* X | Forward_GAGCTCGGGATTTCTCAAGGT<br>Reverse_CAATTCTGCAACACAGAATGTCG |
| RAB59 | *pkal-1(reb28)* X | Forward_GAGCTCGGGATTTCTCAAGGT<br>Reverse_CAATTCTGCAACACAGAATGTCG |
| RAB60 | *C32E8.6(reb23)* I | Forward_GCTTCAACTCCAGAGAATCAGG<br>Reverse_CAACGGCTCTCCGCTCTTAAG |
| RAB61 | *C32E8.6 (reb24)* I | Forward_GCTTCAACTCCAGAGAATCAGG<br>Reverse_CAACGGCTCTCCGCTCTTAAG |
| RAB62 | C24A3.4*(reb16)* X | Forward_CTCTGCCGTACCAGTGATGTTCTA<br>Reverse_CTATCCATGTGCTACCAAACTTGTC |
| RAB89 | *pks-1(reb22[$A_1$_G7106E])* X | Forward_CACCACTATACCAATTCGAAGAACTG<br>Reverse_AGTGACTTGTCAACTTTCCCACTTG |
| RAB103 | *pks-1(reb29[$PCP_2$_S7463A])*X | Forward_GAGACTCACTGAGCAATGAAACTTG<br>Reverse_TCCAGATTTAATCACCTCTTCAGC |

**Supplemental Table 4. Single-worm PCR information used to confirm genotype of wild-type and mutant worm strains used in this study.**

| Strain | Genotype | Wild type | Mutant | Enzyme digestion |
|---|---|---|---|---|
| RAB43 | *nrps-1 (gk186409[$C_4$_S1934N]; gk186410[$C_4$_D1971N])* III | 445 bp | 445 bp | Wild type is cut by BtsIMutI to 265 bp+180 bp; no cut for mutant |
| RAB45 | Y71H2B.1*(gk712674)* III | 700 bp | 700 bp | Wild type is cut by Hinfl into 450 bp+200 bp+50 bp; mutant is cut by Hinfl into 500 bp+200 bp |
| RAB51 | *nrps-1(reb8[$ACP_7$_S307V])* III | 820 bp | 820 bp | Mutant is cut by AatII into 438 bp+382 bp |
| RAB52 | *pks-1(reb9[$C_1$_H6685A])* X | 1098 bp | 1098 bp | Mutant is cut by SphI into 750 bp+348 bp |
| RAB53 | *nrps-1(reb10[$C_3$_H1486A])* III | 993 bp | 993 bp | Mutant is cut by AseI into 658 bp+335 bp |
| RAB54 | *pks-1(reb11[$TE_1$_S7593A])* X | 562 bp | 562 bp | Mutant is cut by SphI into 355 bp+207 bp |
| RAB55 | *nrps-1(reb12[$TE_2$_S2803A])* III | 539 bp | 539 bp | Mutant is cut by NheI into 318 bp+221 bp |
| RAB56 | *pks-1(reb13[$TE_1$_S7593C]; reb14[$TE_1$_G7596A])* X | 574 bp | 574 bp | Mutant is cut by KasI into 364 bp+210 bp |
| RAB57 | *nemt-1(reb15)* IV | 981 bp | 702 bp | |
| RAB58 | *pkal-1(reb21)* X | 1723 bp | 1570 bp | |
| RAB59 | *pkal-1(reb28)* X | 1723 bp | 1171 bp | |
| RAB60 | *C32E8.6(reb23)* I | 1405 bp | 424 bp | |
| RAB61 | *C32E8.6(reb24)* I | 1405 bp | 412 bp | |
| RAB62 | C24A3.4*(reb16)* X | 1863 bp | 502 bp | |
| RAB89 | *pks-1(reb22[$A_1$_G7106E])* X | 1350 bp | 1350 bp | Mutant is cut by SalI into 914 bp+436 bp |
| RAB103 | *pks-1(reb29[$PCP_2$_S7463A])*X | 1057bp | 1057bp | Mutant is cut by SalI into 914 bp+436 bp |

**Supplementary Table 5. sgRNA sequences used for CRISPR-Cas9 in this study.**

| Strain | Genotype (alleles) | sgRNA (20 bases+NGG, and imported vector) |
|---|---|---|
| RAB51 | *nrps-1(reb8[$ACP_7$_S307V])* III | CTCCAGCTCGGCGAGTCTTA AGG (pTM 55-FE*) |
| RAB52 | *pks-1(reb9[$C_1$_H6685A])* X | ATCATATTTTAACTGATGGT TGG (pTM 55-FE*) |
| RAB53 | *nrps-1(reb10[$C_3$_H1486A])* III | GGCTTCTACCATCGCAGATC AGG (pTM 55-FE*) |
| RAB54 | *pks-1(reb11[$TE_1$_S7593A])* X | TTCGTTATGGGGCACTCGAT GGG (pTM 55) |
| | | CTTCGTTATGGGGCACTCGA TGG (pTM 55) |
| | | GTTATGGGGCACTCGATGGG TGG (pTM 55) |
| RAB55 | *nrps-1(reb12[$TE_2$_S2803A])* III | ACCTCTAAATTGGTGTTCAT TGG (pTM 55) |
| | | TTCATTGGCGCCTCGTCTGC TGG (pTM 55) |
| RAB56 | *pks-1(reb13[$TE_1$_S7593C];* | TTCGTTATGGGGCACTCGAT GGG (pTM 55) |
| | *reb14[$TE_1$_G7596A])* X | CTTCGTTATGGGGCACTCGA TGG (pTM 55) |
| | | GTTATGGGGCACTCGATGGG TGG (pTM 55) |
| RAB57 | *nemt-1(reb15)* IV | TATTACTACAGTTATGGCTT TGG (pTM 55-FE*) |
| | | CGAGAAATATGGAACACAGG TGG (pTM 55-FE*) |
| RAB58 | *pkal-1(reb21)* X | AAACTATTGGGCACTTTCGG AGG (pTM 55-FE*) |
| | | CCCAGAATCCAGATGCATGG TGG (pTM 55-FE*) |
| | | TCTTGTGGACATATTCTGCC AGG (pTM 55-FE*) |
| RAB59 | *pkal-1(reb28)* X | AAACTATTGGGCACTTTCGG AGG (pTM 55-FE*) |
| | | CCCAGAATCCAGATGCATGG TGG (pTM 55-FE*) |
| | | TCTTGTGGACATATTCTGCC AGG (pTM 55-FE*) |
| RAB60 | *C32E8.6(reb23)* I | CTTCCATTCTTCCATGCGGG TGG (pTM 55-FE*) |
| | | GACGTGATCCGGAAAGTGGA GGG (pTM 55-FE*) |
| RAB61 | *C32E8.6(reb24)* I | CTTCCATTCTTCCATGCGGG TGG (pTM 55-FE*) |
| | | GACGTGATCCGGAAAGTGGA GGG (pTM 55-FE*) |
| RAB89 | *pks-1(reb22[$A_1$_G7106E])* X | AGGGACACCTGTTGAGCCAC TGG(pTM 55-FE*) |
| RAB103 | *pks-1(reb29[$PCP_2$_S7463A])X* | AGCAATTTGAATAGCATTGA GGG(pTM 55-FE*) |

* pTM 55-FE is a modified version of pTM 55 (a gift of Patrick McGrath) and was mutated to enable a higher level of recognition efficiency[5] by Cas9.

**Supplementary Table 6. Repair templates used for CRISPR-Cas9 in this study.**

| Strain | Genotype (alleles) | Repair template* |
|---|---|---|
| RAB51 | *nrps-1(reb8[ACP$_7$_S307V])* III | ATGAAGTTGAAACCACTCCTCTACCATACCTCGGAATCGACGTCTTAAGACTCGCCGAGCTGGAGTACCACGTGGCTAGT (underlined: AatII; S307V: TCC»GTC ) |
| RAB52 | *pks-1(reb9[C$_1$_H6685A])* X | TGATAACAGTCGAATTCACATCGTTTTCAATCAGCATGCAATTTTAACTGATGGTTGGTCAATGACTGTTCTTTCTGACACTGT (underlined: SphI; H6685A: CAT»GCA ) |
| RAB53 | *nrps-1(reb10[C$_3$_H1486A])* III | CTGGATGAGTAGCAAAAATAAGTTATTGACAATTTCCATTCACGCATTAATCTGCGATGGTAGAAGCCTGCAGATTCTCGAG (underlined: AseI; H1486A: CAC»GCA ) |
| RAB54 | *pks-1(reb11[TE$_1$_S7593A])* X | TGCCGCACATGCCGGAAACAAGAGAATCTTCGTTATGGGGCATGCGATGGGTGGAATAATGAGTCGCGAAATAGTGGCTGAGCTCAAAAT (underlined: SphI; S7593A: TCG»GCG ) |
| RAB55 | *nrps-1(reb12[TE$_2$_S2803A])* III | GTGCTGAAAATATTGAAACCTCTAAATTGGTGTTCATTGGCGCCGCTAGCGCTGGTACTTTTGCATTTTCCACGTCACAACTTTTTG (underlined: NheI; S2803A: TCG»GCT ) |
| RAB56 | *pks-1(reb13[TE$_1$_S7593C]; reb14[TE$_1$_G7596A])* X | TGCCGCACATGCCGGAAACAAGAGAATCTTCGTTATGGGACATTGCATGGGCGCCATAATGAGTCGCGAAATAGTGGCTGAGCTCAAAAT (underlined: KasI; S7593C/G7596A: TCG» TGC/ GGA»GCC) |
| RAB89 | *pks-1(reb22[A$_1$_G7106E])* X | AGAACTCAATTTGGAAGTATTTACTCCATATTCACCAGTGAGTCGACAGGTGTCCCTAAAGGAGTTTGATGGCGGAACAGTCA(underlined: SalI; G7106E: GGC»GAG) |
| RAB103 | *pks-1(reb29[PCP$_2$_S7463A])*X | GTGGCGCCGACAGATAAATTTGAAAGTATTGGTGGAAACGCGTTAAATGCTATTCAAATTGCTCATCGGTTGGCTGAAGAG(underlined: MluI; S7463A: TCC»GCG) |

* The underlined bases indicate restriction sites designed for screening worms containing desired mutations, and the bases labeled in red are coding for the amino acid that was mutated.

**Supplementary Table 7. Primers for construction of translational reporter lines.**

| Strain | Genotype | Primers* |
|---|---|---|
| RAB72 | *rebEx15* | *nemt-1p*_SalI_Fwd_ |
| | | GCGCGTCGACCGAAAATCTCAAGTCTTGTCTTAA |
| | | *nemt-1p*_NotI_Rev_ |
| | | CATGGCGGCCGCAGTGAATATGTGTTTACGAGTAAATG |
| RAB73 | *rebEx16* | *pkal-1p*_SalI_Fwd_ |
| | | GCGCGTCGACGCTATAAATGGGTACCTGGCCGTAA |
| | | *pkal-1p*_NotI_Rev_ |
| | | CATGGCGGCCGCCGTAGAGAAGAAACTGTGACAGTTC |
| RAB74 | *rebEx17* | *C32E8.6p*_SalI_Fwd_ |
| | | GCGCGTCGACGTACCGGGAATCGAAAAATTGTCC |
| | | *C32E8.6p*_NotI_Rev_ |
| | | CATGGCGGCCGCTCTTATTGTACAGAATGTTTCTTTCC |
| RAB76 | *rebEx19* | *nemt-1SL2*_SalI_Fwd_ |
| | | GCGCGTCGACCGAAAATCTCAAGTCTTGTCTTAA |
| | | *nemt-1SL2*_NotI_Rev_ |
| | | CATGGCGGCCGCAGTGAATATGTGTTTACGAGTAAATG |
| RAB77 | *rebEx20* | *pkal-1SL2*_PstI_Fwd_ |
| | | CATGCTGCAGGCTATAAATGGGTACCTGGCCGTAA |
| | | *pkal-1SL2*_NotI_Rev_ |
| | | CATGGCGGCCGCTCAATAGTACATTAGCCTATTCTTTG |
| RAB78 | *rebEx21* | *C32E8.6SL2*_SalI_Fwd_ |
| | | GCGCGTCGACGTACCGGGAATCGAAAAATTGTCC |
| | | *C32E8.6SL2*_NotI_Rev_ |
| | | CATGGCGGCCGCTCAATGCAACATATGACGGACTAG |
| RAB81 | *rebEx22* | Y71H2B.1*SL2*_PstI_Fwd_ |
| | | CATGCTGCAGCCAGATGAAGAAAACGATGGAACT |
| | | Y71H2B.1*SL2*_NotI_Rev_ |
| | | CATGGCGGCCGCTTAAACCCATCCCTCTGGAGGATT |
| RAB82 | *rebEx23* | C24A3.4*SL2*_SalI_Fwd_ |
| | | GCAGGTCGACGGTCTCTACAGTGATGACACTCATT |
| | | C24A3.4*SL2*_NotI_Rev_ |
| | | CATGGCGGCCGCTCACAGCTTGGACCGCGCCGCAAA |

* The underlined bases indicate restriction sites used for plasmid construction.
